# A comprehensive framework for handling location error in animal tracking data

**DOI:** 10.1101/2020.06.12.130195

**Authors:** C. H. Fleming, J. Drescher-Lehman, M. J. Noonan, T. S. B. Akre, D. J. Brown, M. M. Cochrane, N. Dejid, V. DeNicola, C. S. DePerno, J. N. Dunlop, N. P. Gould, A.-L. Harrison, J. Hollins, H. Ishii, Y. Kaneko, R. Kays, S. S. Killen, B. Koeck, S. A. Lambertucci, S. D. LaPoint, E. P. Medici, B.-U. Meyburg, T. A. Miller, R. A. Moen, T. Mueller, T. Pfeiffer, K. N. Pike, A. Roulin, K. Safi, R. Séchaud, A. K. Scharf, J. M. Shephard, J. A. Stabach, K. Stein, C. M. Tonra, K. Yamazaki, W. F. Fagan, J. M. Calabrese

## Abstract

Animal tracking data are being collected more frequently, in greater detail, and on smaller taxa than ever before. These data hold the promise to increase the relevance of animal movement for understanding ecological processes, but this potential will only be fully realized if their accompanying location error is properly addressed. Historically, coarsely-sampled movement data have proved invaluable for understanding large scale processes (e.g., home range, habitat selection, etc.), but modern fine-scale data promise to unlock far more ecological information. While GPS location error can often be ignored in coarsely sampled data, fine-scale data require more care, and tools to do this have not kept pace. Current approaches to dealing with location error largely fall into two categories—either discarding the least accurate location estimates prior to analysis or simultaneously fitting movement and error parameters in a hidden-state model. In some cases these approaches can provide a level of correction, but they have known limitations, and in some cases they can be worse than doing nothing. Here, we provide a general framework to account for location error in the analysis of triangulated and trilatcralizcd animal tracking data, which includes GPS, Argos Doppler-shift, triangulated VHF, trilatcralized acoustic and cellular location data. We apply our error-modelselection framework to 190 GPS, cellular, and acoustic devices representing 27 models from 14 manufacturers. Collectively, these devices were used to track a wide range of taxa comprising birds, fish, reptiles, and mammals of different sizes and with different behaviors, in urban, suburban, and wild settings. In almost half of the tested device models, error-model selection was necessary to obtain the best performing error model, and in almost a quarter of tested device models, the reported DOP values were actually misinformative. Then, using empirical tracking data from multiple species, we provide an overview of modern, error-informed movement analyses, including continuous-time path reconstruction, home-range distribution, home-range overlap, speed, and distance estimation. Adding to these techniques, we introduce new error-informed estimators for outlier detection and autocorrelation visualization. Because error-induced biases depend on many factors—sampling schedule, movement characteristics, tracking device, habitat, etc.—differential bias can easily confound biological inference and lead researchers to draw false conclusions. We demonstrate how error-informed analyses on calibrated tracking data can provide more accurate estimates are that are insensitive to location error, and allow researchers to use all of their data.

## 1 Introduction

Technological advances have ushered movement ecology into a new frontier of high-resolution tracking data on a broad range of taxa (Kays et al., 2015). It is now not uncommon to see tracking data with sampling frequencies of 1 Hz (Munden et al., 2021; Klarevas-Irby et al., 2021). Animal movement data have become central to ecological research and can shed light on previously unanswerable questions relating to behavior, population dynamics, and ecosystem function (Bestley et al., 2015; Kays et al., 2015; Noonan et al., 2015; Strandburg-Peshkin et al., 2015; Lennox et al., 2017; McKinnon and Love, 2018; Noonan et al., 2018; Abernathy et al., 2019). As the quality and quantity of tracking data continue to improve, the scope of inference they provide can expand, provided that analytic methods keep pace (Johnson et al., 2008; Kranstauber et al., 2012; Calabrese et al., 2016; McClintock and Michelot, 2018; Gurarie et al., 2017a; Jonsen et al., 2019). Already, modern movement data are informing on activity and energy budgets (Williams et al., 2014; Noonan et al., 2019), fine-scale habitat use and selection (Ewald et al., 2014), species interactions and encounter processes (Martinez-Garcia et al., 2020; Noonan et al., 2021), but the ability to account for location error is an emerging bottleneck in fine-scale inference (Noonan et al., 2019).

With tracking technology, there are often tradeoffs between device size, battery life, and location estimate precision, with Global Positioning System (GPS; Table 1) and other global navigation satellite system (GNSS) location estimates typically having location errors on the order of meters, Argos Doppler-shift location estimates having errors on the order of kilometers^1^, and light-level geolocator-derived location estimates having errors on the order of hundreds of kilometers (Kaplan and Hegarty, 2006; CLS, 2016; McKinnon and Love, 2018). However, many factors can influence GPS precision, including satellite reception, the specific hardware and software used in a tracking device, and the time-lag between fixes, with more frequent fixes potentially leading to more accurate post-processed estimates (sub-meter with integrated Kálmán filtering, Kreye et al., 2004). Thus, a universal, one-size-fits-all estimate for the location error representing a single class of technology would neglect a large amount of variation.

**Table 1:**
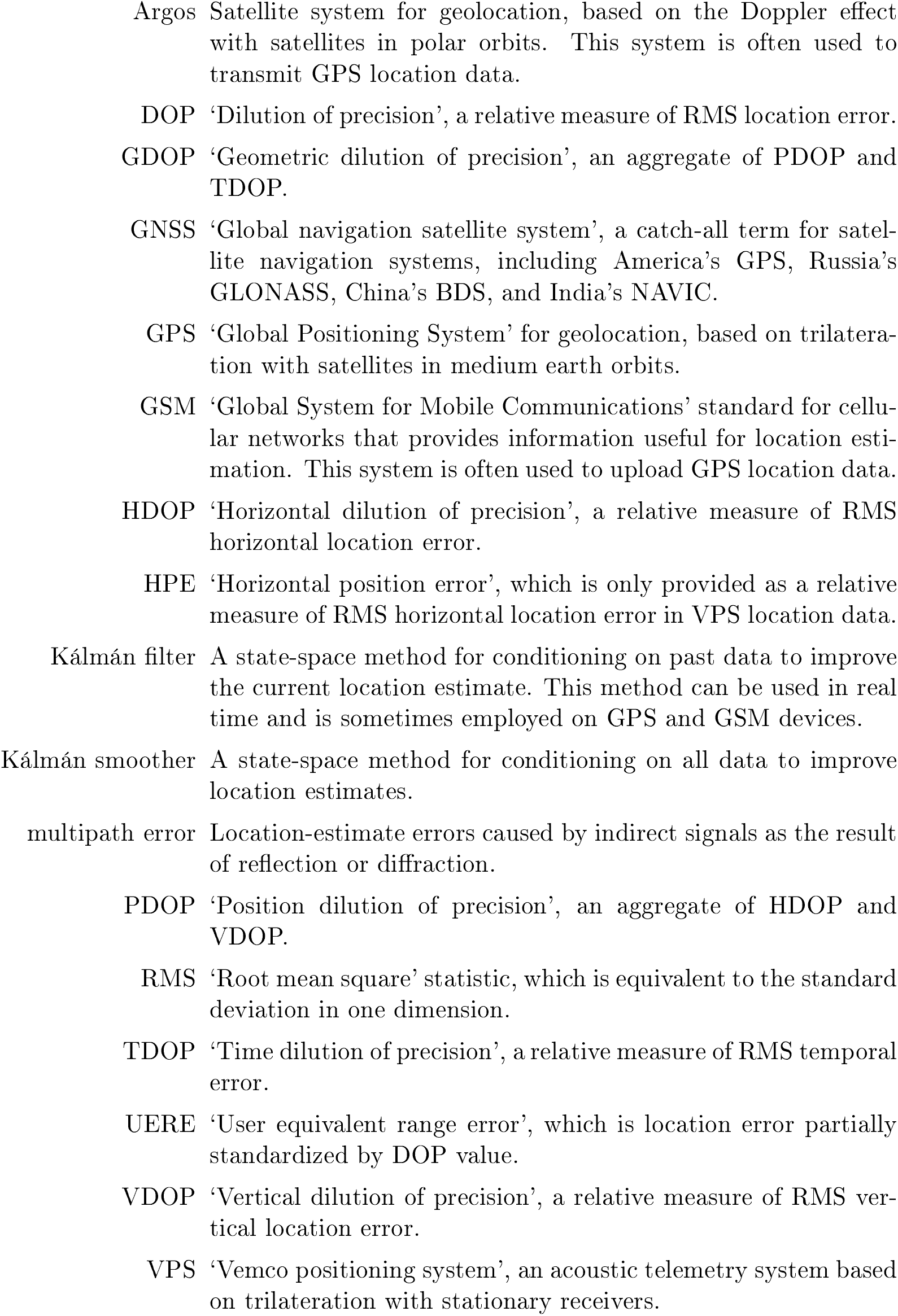
Glossary of terms

Here we highlight the fact that having accurate measures of location error can be essential in providing accurate measurements of animal movement, depending on the relative scales involved. This point has long been recognized by those using less accurate technologies such as Argos Doppler-shift tags, where state-space models have been used to account for location error when tracking animal movement (e.g., Jonsen et al., 2007; Johnson et al., 2008; McClintock et al., 2014). However, if not properly accounted for, we argue that typical location errors of GPS units can also result in mistaken conclusions. For example, Noonan et al. (2019) showed that simple measures of speed are particularly sensitive to location error as step-lengths are overtaken by error. Ross-Smith et al. (2016) and Péron et al. (2017) found it necessary to account for vertical errors in the estimation of flight-altitude distributions. Location error is also important when estimating home ranges from triangulated VHF (Gerber et al., 2018) and Argos Doppler-shift (Poessel et al., 2020) tracking data, and so GPS errors should be expected to especially impact the home-range estimates of animals with smaller home ranges. In all of these cases, biological inference is compromised when location error becomes comparable to the relevant movement scales, and the biases due to mishandling or ignoring location error can easily outstrip the truth (e.g., the more than 10-fold overestimation of wood turtle speed noted by Noonan et al., 2019). As we will elaborate on, these differential biases can cause researchers to draw mistaken conclusions, such as inferring that a species travels faster and more tortuously under dense canopy than in an open habitat.

Aside from outlier classification, which we will consider separately, most animal movement analyses that work with GPS location data simply ignore the issue of location error. Beyond that, conventional solutions to mitigating against location error involve either discarding the most erroneous locations according to a threshold (e.g., Bjørneraas et al., 2010; Poessel et al., 2018*a*,*b*) or simultaneously modeling the movement and error with a hidden state model (e.g., Jonsen et al., 2005; Ross-Smith et al., 2016; Péron et al., 2017). As we detail, the former solution is at best incomplete, while the latter cannot reliably distinguish between variance due to movement and variance due to location error (Auger-Méthé et al., 2016). Related to these issues is the problem of outlier detection, which has, until now, relied on metrics assuming no location error, in accordance with the above principle of thresholding.

Here we present comprehensive methods and guidelines for dealing with location error (Fig. 1), largely focused on GPS, but also including Argos Doppler-shift, Global System for Mobile Communications (GSM), acoustic trilateralization, and other tracking technologies that provide location estimates. First, we put forward the need and best practices for the error calibration of tags and encourage researchers to obtain this information before they start collecting tracking data. Second, we provide a statistically efficient framework for quantifying location error, showing how to create and select among error models with 190 example calibration datasets. Finally, we demonstrate how calibrated tracking data then serve as ideal inputs for error-informed movement analyses, including path reconstruction, connectivity quantification, home-range estimation, and distance estimation.

**Figure 1:**
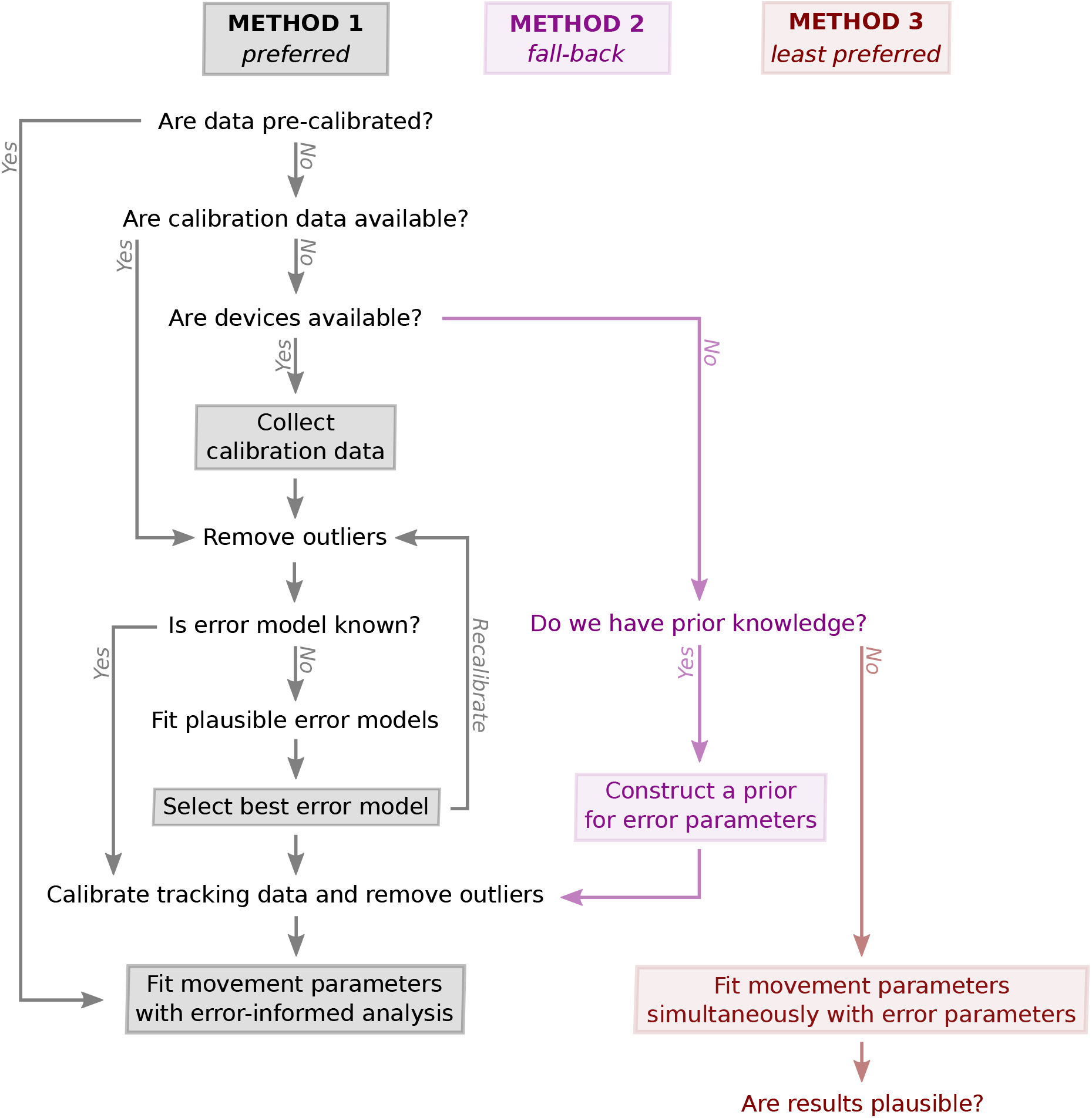
Proposed workflow to account for location error in animal movement analysis, with the primary method (1) that requires calibration data in black, the secondary method (2) that requires prior knowledge of calibration parameters in magenta, and the tertiary method (3) of simultaneous estimation in red, which is the conventional method (e.g., Jonsen et al., 2005) and is vulnerable to issues of statistical identifiability (Auger-Méthé et al., 2016). We note that most of these steps are avoidable. In particular, if the tracking data are supplied precalibrated, then researchers can proceed immediately to error-informed movement analysis.

### Tracking data and their error information

Satellite location data are accompanied by variable degrees of location error—on the order of 1–10 meters for differential and tri-lane GPS location error, 10–100 meters for conventional GPS horizontal error, several times that for GPS vertical error, and 100–10,000 meters for Argos Doppler-shift horizontal error (Kaplan and Hegarty, 2006; Parkinson and Enge, 1996; CLS, 2016). Location errors are generally *heteroskedastic*—meaning that their variances can change in time, depending on the reception and spatial configuration of the satellites and tracking device. While older Argos Doppler-shift location data, prior to Lopez et al. (2014), only come with error classes that must be estimated (Vincent et al., 2002) or fit simultaneously with a movement model (e.g., Jonsen et al., 2005; Johnson et al., 2008), modern Argos data come with 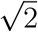-standard-deviation error ellipses, estimated by a custom multiple-model Kálmán filter (Lopez et al., 2014). Error ellipses are more necessary with Argos Doppler-shift location estimates because the polar orbits of Argos satellites provide better resolution of latitudes than longitudes (Costa et al., 2010). Argos Doppler-shift location data were also the first to be modeled via state-space models incorporating both continuous-time non-Markovian movement processes and heteroskedastic errors (Johnson et al., 2008).

GPS device manufacturers do not generally provide calibrated error circles of known quantile or standard deviation. Instead, GPS location estimates typically come with unitless and device-specific “horizontal dilution of precision” (HDOP) and “vertical dilution of precision” (VDOP) values. Ideally, DOP values are designed to account for the order-of-magnitude heteroskedasticity stemming from satellite reception, geometry, latitude, and dynamics (Kaplan and Hegarty, 2006; Yahya and Kamarudin, 2008), which, for the purpose of animal tracking, includes the effects of local habitat, canopy, cover, burrowing, device orientation, speed, and most everything else short of space-weather events (Coster and Komjathy, 2008). DOP values provide the time-dependent transformation

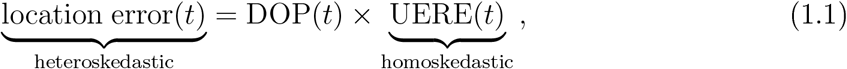

between heteroskedastic location error and homoskedastic “user equivalent range error” (UERE). UEREs can be thought of as partially standardized location errors, in that they are proportional to the true location errors, but have a fixed (homoskedastic) variance. In the null model (1.1), UEREs are assumed to be a mean-zero, stationary Gaussian process (Kaplan and Hegarty, 2006). By quantifying the relatively simple distribution of UEREs that can often be described by a single scale parameter, we can back-transform with DOP values to quantify the time-dependent distribution of location errors. Traditionally, the UERE scale parameter estimated is the root-mean-square (RMS) UERE. We refer to the direct estimation of UERE parameters—and subsequent back-transformation via relation (1.1)—as error ‘calibration’. However, GPS location data are sometimes missing relevant DOP information and may exhibit more complex error structures than present in the null model (1.1), which can necessitate error-model selection.

Triangulated VHF location errors tend to be intermediate to GPS and Argos Doppler-shift errors in magnitude, and their location estimates come equipped with error ellipses when using the method of Lenth (1981), such as from `R` package `sigloc` (Berg, 2015). (Also, see Gerber et al. (2018) for full posterior estimation of the location-error distributions.) Therefore, operationally speaking, triangulated VHF tracking data can be treated analogously to modern Argos Doppler-shift data.

Acoustic trilateralization is more popular in marine systems, where electromagnetic signals rapidly attenuate. These systems operate much like GPS, but via acoustic signals with receivers fixed underwater instead of electromagnetic signals with satellites in orbit. In particular, Vemco positioning system (VPS) location estimates come with “horizontal position error” (HPE) values, which Vemco documentation describes as being analogous to GPS HDOP, in that they are proportional to error-circle radii and must be calibrated on a per-network basis (Smith, 2013).

‘Global System for Mobile Communications’ (GSM) cellular networks provide a number of data sources that devices can use to estimate location, such as time-of-arrival, angle-of-arrival (azimuth), and signal strength with respect to multiple cellular towers (Martinez Hernandez et al., 2019). Resulting location estimates can vary in quality, depending on the techniques employed and network coverage, and may return error-parameter estimates analogous to GPS, though error ellipses may be more desirable. GSM-derived location errors tend to be larger than those of GPS, due to the smaller number of cellular towers that are typically within range.

In summary and irrespective of how they are collected, tracking data that provide location estimates usually come with either error-circle (GPS, GSM, VPS) or error-ellipse (triangulated VHF, Argos Doppler-shift, geolocator) information. Sometimes this information is only proportional to the location error’s variance (GPS, GSM, VPS), and further estimation is required to fully quantify the error distribution.

### When can location error be reasonably ignored?

It can be reasonable and pragmatic to simply ignore location error in a study when the scales of error are far smaller than all relevant scales of movement. For example, 10-100 meter GPS error can be considered insubstantial in comparison to a long-distance migration sampled on a daily schedule, where individuals travel many kilometers between location fixes. In situations where this comparison is less clear cut, researchers can perform a sensitivity analysis to approximate the impact of ignoring location error via simulation (App. S1). Because these sensitivity estimates are approximate and require as much effort as more-exact, error-informed analysis, we only recommend this procedure in cases where there is no error-informed analysis available.

### Conventional approaches to handling location error

Beyond outlier detection, the most common approach in the literature to deal with location error is simply to discard the most erroneous location estimates, either with an arbitrary HDOP threshold for GPS data or a location-class threshold for Argos Doppler-shift data (Bjørneraas et al., 2010). This threshold is especially arbitrary for GPS data, as HDOP values do not have the same meaning across device models, even when assuming the null model of location error = DOP × UERE (1.1), and complicated by the fact that most manufacturers do not provide UERE parameters for their wildlife telemetry devices. Consider, for example, that if the RMS UERE of device model A is five times that of device model B, which we will show to be a realistic hypothetical, then an HDOP value of 10 on device model B is equivalent to an HDOP value of 2 on device model A. As an improved criterion, Meckley et al. (2014) argued for objective thresholding, based on calibrated error estimates (in meters), rather than unitless precision estimates that vary by device model. However, thresholding the data still involves a trade off, whereby reduced bias comes at the cost of a smaller sample size. At both extremes—large included location errors and small sample sizes—there can be substantial bias from too much location error or not enough data, and an intermediate threshold that can sufficiently mitigate both biases may not exist (Noonan et al., 2019).

Rather than discarding data, state-space models consider the location fix as the combination of both movement and location error, with unknown parameters for each model. State-space models were first introduced to wildlife tracking as a rigorous method to account for location error by Anderson-Sprecher and Ledolter (1991), and their popularity grew rapidly with the application to geolocation (Sibert et al., 2003) and Argos Doppler-shift (Jonsen et al., 2003) tracking data, and with capable software packages, such as `bsam` (Jonsen et al., 2003, 2005) and `crawl` (Johnson et al., 2008). State-space models are another common approach to account for location error, and are more-than-appropriate for this task, but they are not without their challenges. As has been pointed out by Auger-Méthé et al. (2016), state-space models can misappropriate variance due to movement and location error when the two sets of parameters are fit simultaneously to tracking data, as is common practice in ecology (Fig. 2). Fundamentally, this misappropriation is due to a lack of statistical *identifiability—*if movement and error models can produce similar outputs, then their parameters (and influences) cannot be reliably teased apart (Hilborn and Mangel, 1997). In application, the movement and location-error parameters—and, in particular, the location-error variances—are almost always simultaneously estimated from tracking data (e.g., Jonsen et al., 2005; Kuhn et al., 2009; Swimmer et al., 2009; Howell et al., 2010; Sulikowski et al., 2010; Vermard et al., 2010; Matthews et al., 2011; Blanco et al., 2012; Carman et al., 2012; Hückstädt et al., 2014; Dalleau et al., 2014; Kennedy et al., 2014; Ross-Smith et al., 2016; Afonso et al., 2017; Péron et al., 2017), which means that the identifiability issue is a pressing challenge.

**Figure 2:**
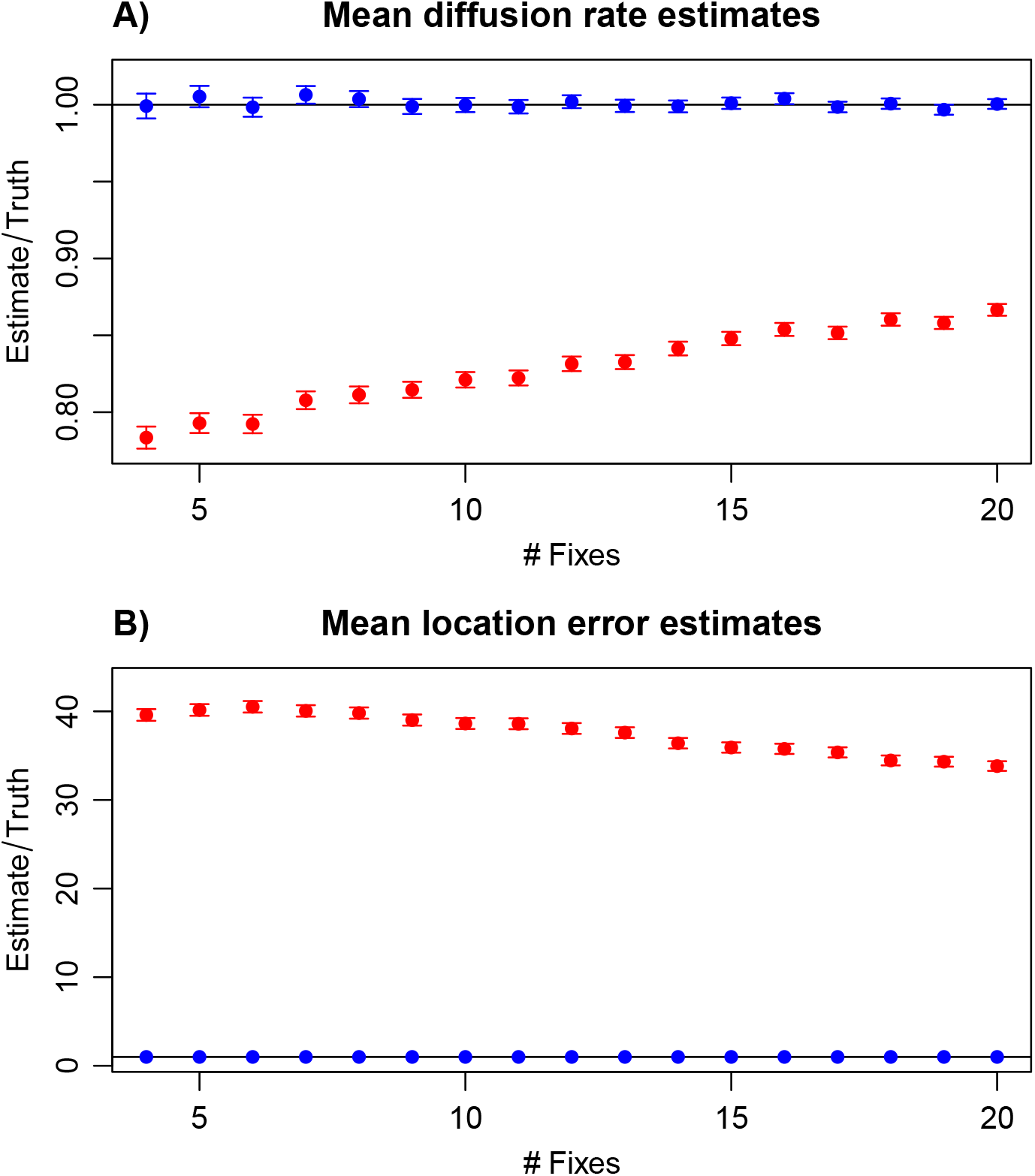
Here we estimate the diffusion rate (**A**) and root-mean-square (RMS) location error (**B**) of a Brownian motion process with RMS location error one-hundredth that of the movement per unit timestep (e.g., 1 km steps and 10 m error), via our proposed method (blue) and the conventional method (red). In the conventional method (red), the movement and location-error parameters are fit simultaneously to tracking data. Because of the identifiability issues pointed out by Augcr-Mćthć et al. (2016), the conventional method attributes a large portion of the movement to location error. In our proposed method (blue), all error parameters are estimated from calibration data before fitting to tracking data. Each mean estimate is calculated from 20,000 replicates and the resulting 95% confidence intervals are shown.

Here we build on the state-space modeling framework and show that performing an integrated analysis, on both tracking data and calibration data, offers a straightforward solution to the problem of statistical identifiability (Auger-Méthé et al., 2016), and the compounding problem of movement-model misspecification. Calibration data are routinely employed with geolocation data (Thiebot and Pinaud, 2010; Lisovski et al., 2020), and were used by Jonsen et al. (2005) to provide Argos Doppler-shift location classes with approximate DOP-value ellipses, which is a practice that remains popular today, though still with the simultaneous estimation of movement and location-error variance parameters. Citing the lack of identifiability between movement and location-error parameters, Block et al. (2011) estimated geolocation errors from calibration data and assumed the resulting estimates to be fixed in their state-space model. Here we also use calibration data to inform difficult-to-identify parameters in state-space models, by performing an integrated analysis on animal tracking and calibration data, without assuming that their location errors are of the same variance. When calibration data are abundant, this results in the location-error parameters being essentially fixed by the calibration data (similar to Block et al., 2011), while small amounts of calibration data or prior knowledge can still improve identifiability. Thus, an integrated analysis provides a general solution to an important statistical challenge compromising the use of state-space models on animal movement data. For the sake of brevity, we will simply refer to the use of calibration data to provide informative priors and model selection criteria for all location-error model parameters as “calibration”, while acknowledging preceding uses of calibration data (Jonsen et al., 2005; Block et al., 2011). Additionally, there is an abundance of historical tracking data for which no calibration data exist, and the simultaneous fitting of movement and location-error parameters performs better in some situations than others. Therefore, it is still useful to understand under what circumstances simultaneous fitting (Jonsen et al., 2005) can perform well.

Finally, in addition to differences in how location error can be modeled and estimated, there are also differences in how these error models are propagated through movement analyses. The most convenient and approximate correction is to “smooth” the data (Auger-Méthé et al., 2021)—which involves calculating predictions of the true locations, conditional on the entire track—before feeding the smoothed data into movement analyses that assume no location error. However, as smoothing does not fully resolve the true locations, there will be a degree of bias and unpropagated uncertainty in the smoothed analysis (for some comparisons, see Noonan et al., 2019). Going one step further, posterior inference (Johnson et al., 2011; Gerber et al., 2018; Noonan et al., 2019) can offer more rigorous solutions, by analyzing ensembles of conditional simulations, rather than analyzing the smoothed data, which are only the conditional predictions. Even when fed into movement analyses that cannot account for location error on their own, first-order bias and uncertainty corrections can be calculated via simulation, as we detail in App. S1. Ideally, however, movement analyses should be fully error informed—meaning that in no stage of the calculation is location error assumed to be absent, which is the case for all of the examples we cover here.

### A comprehensive framework for addressing location error

Our proposed framework to account for location error in animal tracking data, of the trilaterlized, triangulated, or directly estimated variety, is a general approach with three basic steps: 1.) calibration-data collection, 2.) error-model selection, and 3.) error-informed analysis. This approach is summarized in Fig. 1 and briefly described below. It builds on the common state-space formalism (Anderson-Sprecher and Ledolter, 1991) and the general concept of calibration (e.g., Block et al., 2011), while the error-model selection and specifics of the “calibration” step are new ingredients that we demonstrate are necessary for the accurate accounting of location error—particularly in GPS data and in considering the results of Auger-Méthé et al. (2016). Calibration-data collection is detailed more thoroughly in Sec. 2.1. Error-model selection is then covered in Sec. 2.2, with derivations given in App. S2–S3, and 27 error-model-selection examples provided in App. S4. Finally, Sec. 3 presents empirical examples of error-informed movement analyses on tracking data, where accounting for location error impacts inference.

#### 1.) Calibration-data collection

First, we address the lack of statistical identifiability between movement and location-error parameters when estimating the two simultaneously in a state-space framework. Instead of simultaneous estimation, we propose calibrating all tracking data before movement analysis. In the calibration step, researchers collect location data that have a known movement model, which we refer to as calibration data. Most simply, we suggest recording location data at fixed locations, so that all variance in the calibration data is solely due to location error. Error-model parameters can then be fit to the calibration data, which addresses the identifiability issues pointed out by Augcr-Mćthć et al. (2016). Importantly, relation (1.1) does not assume that the location errors of the tracking data and calibration data have the same distribution and, instead, it explains their differences. Finally, the location-error parameter estimates can be updated in an integrated model of the calibration and tracking data, which also propagates their uncertainties into the movement analysis.

#### 2.) Error-model selection

Second, we introduce a novel error-model selection framework, including a number of plausible candidate models and a statistically efficient modelselection criterion. Model selection is standard practice in ecology (Burnham and Anderson, 2002), but it has not been applied to location-error models, in practice. Existing studies have typically assumed the null model (1.1) or a model of homoskedastic location errors, and sometimes other covariates are more informative. To construct plausible location-error models, we discuss relevant predictors in App. S3—including DOP values, the number of satellites in reception, and various location-class proxies—and apply these to empirical data in App. S4. Then, given multiple candidate error models, we introduce a new model-selection framework based on corrected Akaike information criterion (AIC_C_) values (App. S2). AIC-based model selection achieves asymptotically optimal predictions (Yang, 2005), and is known to perform well when selecting among a small number of plausible candidate models, while ‘corrected’ AIC_C_ values are adjusted for small-sample-size bias (Burnham and Anderson, 2002), which is necessary when the calibration data are limited. Finally, to objectively gauge the selected model’s performance, we derive a goodness-of-fit statistic tailored specifically to the needs of error-model evaluation (App. S2.1.2).

#### 3.) Error-informed analyses

The third and final step of our proposed framework is to feed these calibrated data into error-informed—and preferably, continuous-time—analyses. Both the `crawl` (Johnson, 2008; Johnson et al., 2008) and `ctmm` (Fleming and Calabrese, 2015; Calabrese et al., 2016) `R` packages are capable of performing a large number of continuous-time movement analyses while accounting for (calibrated) heteroskedastic location errors in a state-space framework, including path reconstruction, conditional simulation, and the calculation of utilization distributions (Fleming et al., 2015, 2016, 2017). The family of Brownian-bridge methods within `R` package `move` (Kranstauber et al., 2012) can also incorporate calibrated location errors, as can the behavioral-segmentation methods in `R` packages `momentuHMM` (McClintock and Michelot, 2018) and `smoove` (Gurarie et al., 2017*b*)—both by leveraging `crawl`.

Here, we expand on these methods by introducing error-informed statistics for outlier detection and variogram-based autocorrelation visualization. We provide a thoroughly tested and documented software implementation for all analyses in the CRAN-hosted `ctmm R` package, complete with both help files and long-form documentation via `vignette (‘error’)`. Finally, we demonstrate the appropriateness and utility of these methods with several empirical examples, where properly accounting for location error makes the difference between accurate inference and qualitatively incorrect inference.

## 2 Calibration-data collection and error-model selection

### 2.1 How to collect calibration data

In the simplest case, ‘calibration data’ are location fixes obtained when a tracking device is not moving. Error-model parameters are best estimated when the true movement model is known, to avoid the potential for misspecification. The simplest movement model possible is one of no movement. Therefore, to collect calibration data, tracking devices should be left at fixed locations over an extended period of time—preferably for days. The true locations do not need to be known a priori; these can be estimated simultaneously with the UERE parameters, in a statistically efficient manner.

#### How many tracking devices should be tested?

Ideally, only one device per model is needed to collect calibration data, as all devices of the same make and model should have similar UERE distributions. However, multiple devices of the same model can be deployed and the assumption of identical devices can be tested via model selection. If UERE parameter estimates are consistent, then those calibration data can be integrated together to obtain more accurate estimates. If calibration data come at the cost of limited battery life that would otherwise be used to collect animal tracking data, we suggest a two-stage calibration deployment when device heterogeneity is a concern. In the first round of deployments, a small amount of pilot calibration data are collected for each device, such that the total number of sampled locations can meet the target accuracy, were the UERE distributions identical. Discrepancy among devices can then be tested for and, if found significant, further per-device calibration data can be collected in a second round of deployments. Otherwise, if UERE parameter estimates are found to be consistent, the single estimate is sufficient.

#### How much calibration data should be collected?

UERE parameters are estimated from data, and so increasing the sample size will increase their precision. At minimum, calibration data should be collected over the course of a day, to sample a wide range of satellite configurations and DOP values. Researchers may want to collect many days of calibration data if space weather events are of concern (Coster and Komjathy, 2008). More location fixes will provide more precise UERE statistics, but the sampling frequency should not be so high as to induce location-error autocorrelation not present in the tracking data. For GPS devices featuring on-board location-error filtering, the RMS UERE may shrink at higher sampling rates (~1 Hz) and, therefore, the sampling schedule of the calibration data should be the same as that of the tracking data. If the same device has been used at disparate sampling rates, then the presence of an on-board error filter can be tested with calibration data collected over the same range of values.

Relative error in a single mean-square (MS) UERE estimate will be of standard deviation 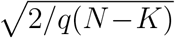 (c.f., Eq. S2.7), where *q* is the number of spatial dimensions (*q* = 2 for horizontal UEREs and *q* = 1 for vertical UEREs), *N* is the total number of sampled locations, and *K* is the number of unknown locations where calibration data were collected. With only one calibration deployment per tracking device, *K* is simply the number of devices. Therefore, given some relative error tolerance 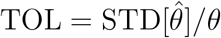 in the MS UERE estimate 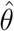, the *total* number of calibration fixes necessary to meet that tolerance in the MS UERE estimate is

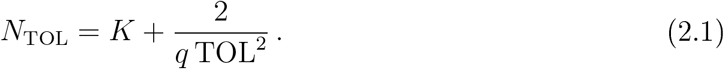

E.g., for 10 devices in two dimensions and deployed once each, the requisite number of calibration fixes *per device* is 41 (or 1001) for relative error tolerances of 5% (or 1%). If the 10 devices are found to have significantly different UERE parameters, then in the second round of deployments an additional 361 (or 9001) calibration fixes must be collected per device, to meet the same relative error tolerance in the individualized parameter estimates.

#### Where should calibration data be collected?

For newer tracking devices that estimate reliable DOP values and mitigate against multipath errors (Table 1), it should not be necessary to sample calibration data in multiple habitats or latitudes. The effect of variable signal reception across different habitat types should be accounted for in the DOP values (Kaplan and Hegarty, 2006; Yahya and Kamarudin, 2008). But, again, this assumption can be challenged, by first collecting a small amount calibration data in each habitat, testing for discrepancy, and then collecting more calibration data if necessary. For devices that feature multiple ‘location classes’—location estimates of substantially different precision, regardless of DOP value—it may be important to sample in areas of poorer satellite reception, because if only the best location classes are observed in the calibration data, then the worst location classes cannot be reliably calibrated. Furthermore, to test the veracity of a device’s DOP values, less-than-ideal signal reception may be preferable to magnify their presumed impact.

#### Opportunistic calibration data

For studies where calibration data were not collected and cannot be collected after the fact, it still may be possible to assemble calibration data opportunistically. Segments of time when the individual was deceased, at a rest site, or when the tracking device was detached may be subsetted and treated as calibration data. Researchers should be especially careful that opportunistic calibration data are free of movement. Methods to test for the presence of movement in relocation data include correlogram analysis (Fig. S4.1) and attempting to fit an autocorrelated movement model (simultaneous with unknown UERE parameters) to the data. However, both of these approaches assume that the data are sampled finely enough that movement and error are easily distinguishable, yet coarsely enough that the location error itself is not autocorrelated. This will typically be the case for GPS location estimates sampled at least 1–2 minutes apart (Tao and Bonnifait, 2015).

### 2.2 An error-model-selection framework

After error calibration data have been collected, an appropriate error model can be selected. While model selection is ubiquitous across ecology (Burnham and Anderson, 2002), it has not been traditionally applied to location error models. In the null model of GPS location error (1.1), DOP values contain all dependence on satellite geometry, including the number of satellites in communication. On the other hand, UEREs are generally assumed to be a mean-zero Gaussian process (Kaplan and Hegarty, 2006), characterized by its scale parameter (e.g., RMS UERE), even though the location errors may have a heavy-tailed distribution (Fig. 3). However, complications may arise, such as if DOP values are not recorded by the device or there are multiple location classes with different UERE distributions. Different error models may be appropriate in different situations, and error-model selection is frequently necessary. Therefore, in App. S2 we derive RMS UERE estimators, AIC_C_ values, and goodness-of-fit statistics for location-error models fit to calibration data. We have focused on the statistical efficiency of our estimators, which have ideal performance for one location class and normally distributed UEREs, including for the null model (1.1), and better performance than maximum likelihood more broadly. Unbiased estimators are important when pooling small amounts of calibration data, and when calibrating multiple location classes where some fix types rarely occur (c.f. S4.11).

**Figure 3:**
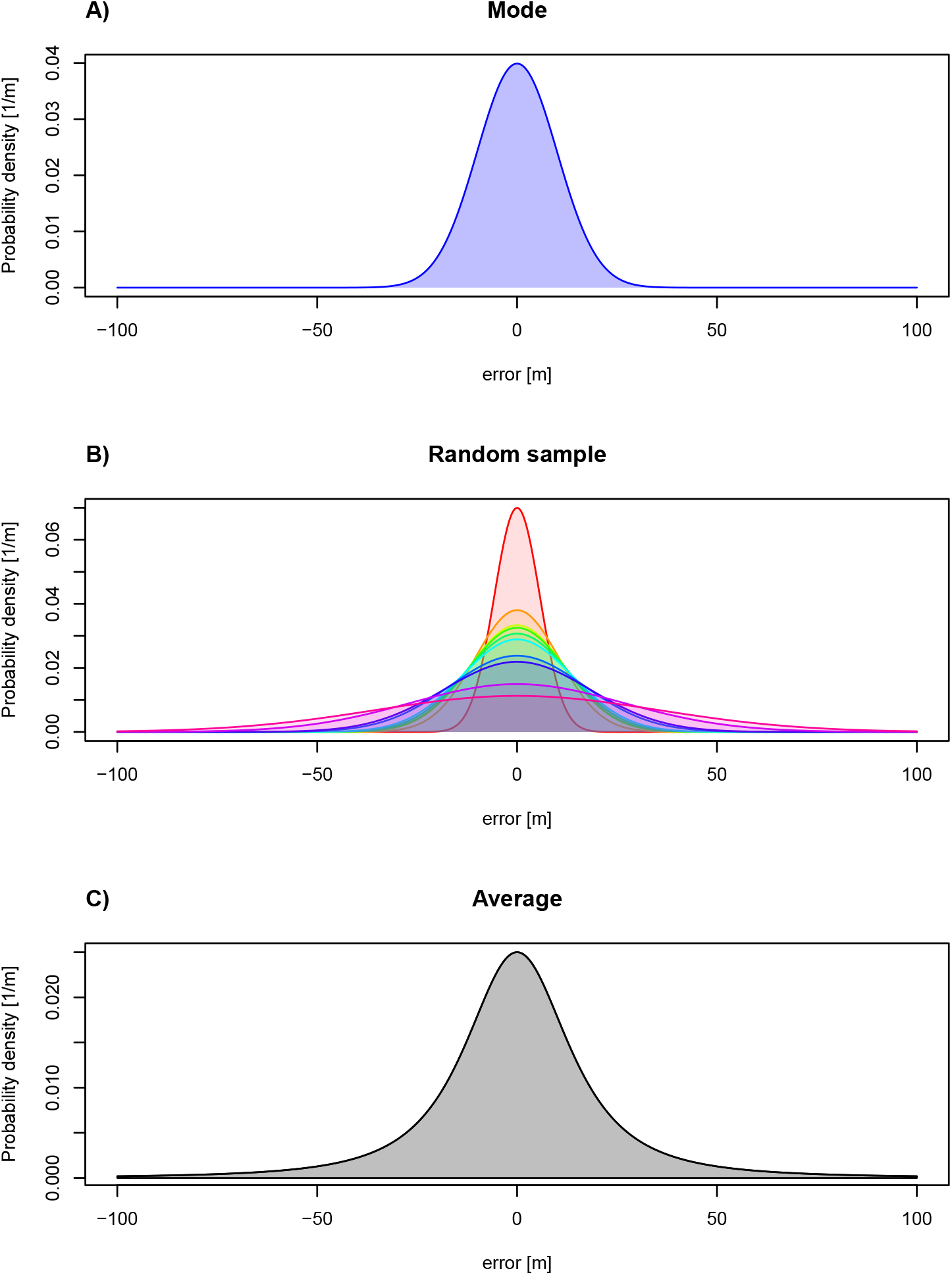
Demonstration of how ignored heteroskedasticity (time-varying variance) can lead to heavy-tailed distributions. In subpanel **A**, the location-error distribution is given for the most common variance, as a representative example. This distribution is Gaussian with 10-meter RMS error, and is light tailed. In subpanel **B**, 10 random HDOP values are drawn with resulting location-error distributions overlayed. Each individual distribution is light tailed, but the scale of location error varies dramatically among them. In subpanel **C**, the average location error is considered, which is proportionally *t*-distributed. This final distribution is what is considered when ignoring heteroskedasticity, and is heavy tailed.

As tracking devices can vary in behavior and poorly specified location-error models can lead to heavy-tailed UERE distributions, error-model selection can be important for making the best use of tracking data. AIC values serve as a model selection criterion that balances goodness of fit against parsimony (Burnham and Anderson, 2002) and can provide asymptotically optimal predictions (Yang, 2005). However, in situations where sample sizes are small and unbiased parameter estimates differ substantially from (biased) maximumlikelihood parameter estimates, ‘corrected’ AIC_C_ values can differ substantially from regular AIC values. Furthermore, it is not sufficient to adopt off-the-shelf AIC_C_ formulas, because they are both model and estimator dependent (Calabrese et al., 2018; Fleming et al., 2019), and the most commonly used AIC_C_ formula of 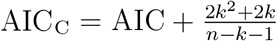 is derived specifically for linear regression models estimated via maximum likelihood (Burnham and Anderson, 2002). Therefore, our AIC_C_ model-selection criterion is based on newly derived formulas, specific to our model structures and debiased parameter estimators (App. S2.1.1 & S2.2.1).

While useful as a model selection criterion, AIC_C_ values scale with the amount of data and cannot be used to assess a model’s goodness-of-fit for comparing performance across devices (Burnham and Anderson, 2002). Therefore, we develop an absolute goodness-of-fit statistic—reduced *Z*^2^ or 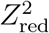—that can be compared across animals, devices, and studies. 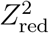 is an error-model analog to the well-known reduced chi-squared statistic, 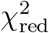, but based on Fisher’s *Z* distribution (App. S2.1.2 & S2.1.2). As with 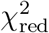 smaller values of 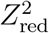 denote better performance and knowing the true model results in 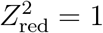 on average.

Historically, studies have either assumed the null model (1.1) or a model of homoskedastic location errors. However, we find that the best performing models can have more complicated structures. In App. S3 we introduce a number of novel error models for GPS devices with missing or incomplete DOP information. The predictors covered include the hierarchy of DOP values (HDOP, VDOP, PDOP and GDOP— Table 1), approximating DOP values with the number of satellites in reception, and the possibility of GPS fix type and time-to-fix as modifiers. We then apply these error models to 190 GPS, GSM, and VPS devices representing 27 device models from 14 manufacturers in App. S4, using our error-model-selection framework with `R` implementation in the `ctmm` package (Fleming and Calabrese, 2015; Calabrese et al., 2016).

For most of the devices we analyze in App. S4, `ctmm`’s null location-error model—based on the model ranking in App. S3—is the selected error model, and the calibration step is as simple as the `R` assignment

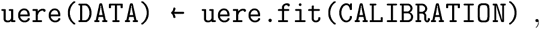

where `DATA` refers to the tracking data and `CALIBRATION` refers to the calibration data. For pre-calibrated location data, such as Argos Doppler-shift^2^ and triangulated VHF location estimates, no calibration steps need to be taken, while problematic devices exhibiting a statistically inefficient null model will require model selection. A worked example is given in `vignette(‘error’)` within `R` package `ctmm`.

### 2.3 Error or outlier?

A dual task to modeling location error is the classification of outliers, which are observations that defy the combined movement and error model (Fig. 4). For instance, a recorded flight altitude only slightly below ground with a large corresponding VDOP could be considered typical (*p*-value ≫ 0), while a recorded flight altitude deep below ground with a small corresponding VDOP should be extraordinarily rare (*p*-value ≈ 0) under normal circumstances and therefore considered an outlier. Ordinarily erroneous data are not outliers and can still contain meaningful information that should be retained, as long as the applied statistical framework incorporates an appropriate error model. Discarding data that are not true outliers can bias inference (Péron et al., 2020).

**Figure 4:**
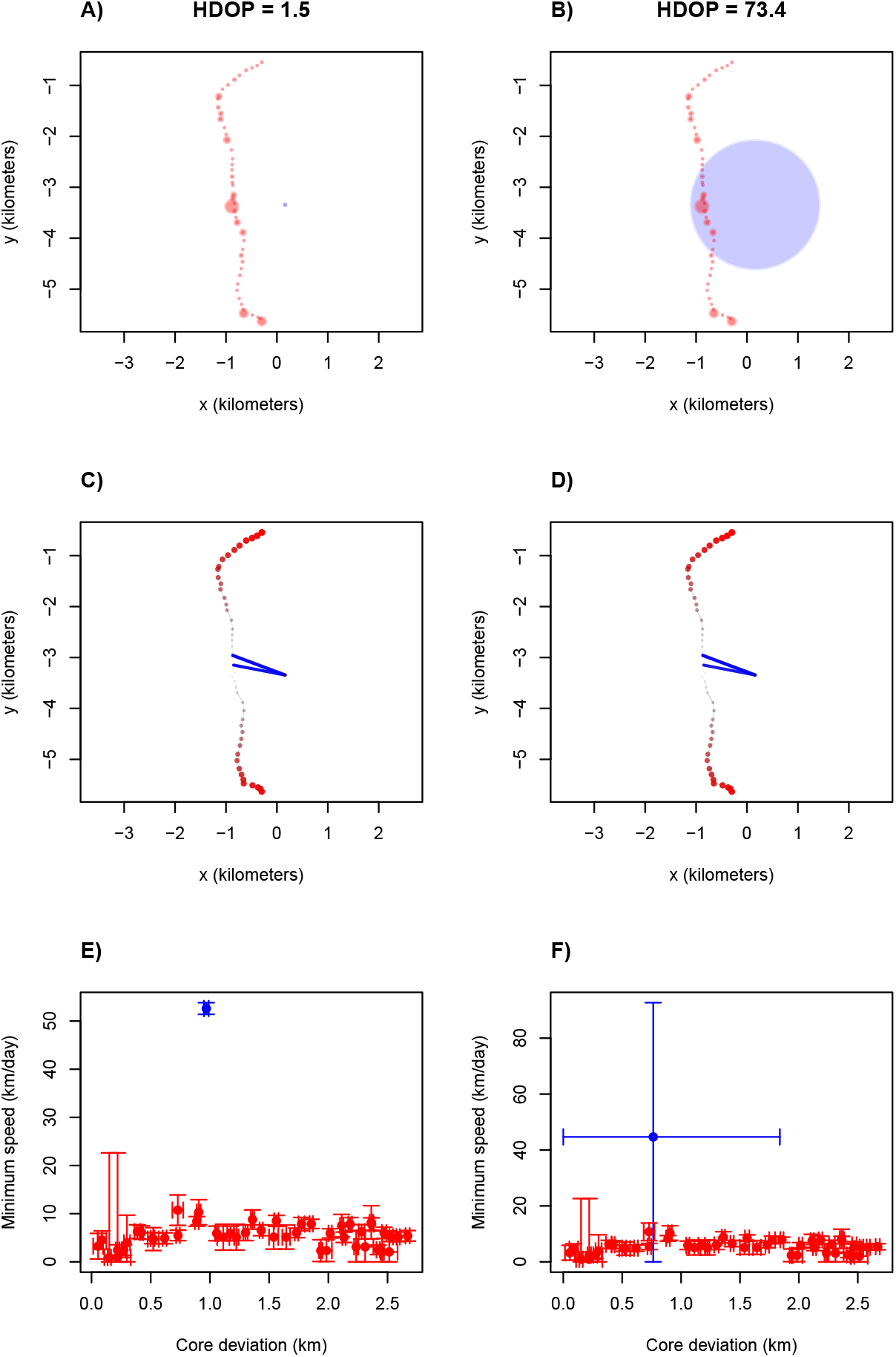
Simulated animal tracks with location error (**A-B**) and a visual analysis of outliers (**C-F**). In the first column, an outlier is introduced—colored blue in the first and third rows— where the corresponding horizontal dilution of precision (HDOP) value is underestimated by a factor of 50. The second column contains the same location estimates as the first column, but where all HDOP values are accurate. Scatter plots are provided in the first row, with 95% error circles. Error-informed speed and distance estimates are calculated and used to color the data in row two, where fast transits are more blue and remote locations are more red. Finally, the same speed and distance point estimates from the second row are given confidence intervals and plotted in the third row. ‘Minimum speed’ refers to the minimum speed capable of explaining the data, while ‘core deviation’ refers to distance from the median location (see App. S5 for precise explanation of these quantities). The obvious outlier in panels **A & E** is not an outlier in panels panels **B & F**.

The classification of outliers aside, the second problem is what to do with them. If the data contain many outliers, then the ideal solution is to improve the error model until said outliers can be considered normal (c.f., App. S4.5.2). However, if the number of outliers is small, then requisite error-model parameters may not be well resolved, and even so, the information gained by incorporating these data may be insubstantial. Therefore, removing outliers can be a pragmatic choice, especially in animal tracking data, which can be plagued with baffling abnormalities. In any case, these decisions should always be reported with resulting analyses.

It is standard practice to classify outliers in animal tracking data using simple metrics such as speed and distance, so that location estimates implying implausible movements can be ruled out (Douglas et al., 2012; Safi and Kranstauber, 2021). However, traditionally these outlier metrics have not been informed by location error, and so they can confound normal errors with true outliers. For example, straight-line distance is commonly used as an estimate of the minimal path-length requirement between sequential location estimates, yet location errors can cause this calculation to overestimate true distances (Ranacher et al., 2016; Noonan et al., 2019). In App. S5 we derive maximum-likelihood distance and speed estimators that account for location error and are minimally conditioned on the data. These estimators are implemented in the `ctmm` (Fleming and Calabrese, 2015; Calabrese et al., 2016) method `outlie ()`, and demonstrated in Fig 4. While the example in Fig. 4 is visually striking, outliers in finely sampled data are not generally obvious, and their implausibility cannot be properly discriminated without accounting for the associated location errors.

Location-error outliers can occur in both tracking and calibration data. Identifying outliers in calibration data, therefore, may require some degree of iteration, whereby researchers first assume a plausible RMS UERE value and then calculate a more accurate RMS UERE value after removing all contingent outliers. The updated RMS UERE can then be checked for consistency by applying it to the initial data (with outliers intact) and determining if the same classification of outliers is recovered (Fig. 1).

## 3 Empirical examples

We present two sets of empirical analyses—one set on an expansive collection of calibration data and another set on individual tracked animals. First, we analyze the quality of error information provided by tracking devices in Sec. 3.1. Second, we demonstrate the impact of proper error modeling and error-informed methods on biological inference in Secs. 3.2-3.4.

### 3.1 Device calibrations

Here we summarize the application of our error-model-selection techniques to 190 GPS, cellular, and VPS devices representing 27 device models from 14 manufacturers. These tracking devices are used on a number of species, including American black bear (*Ursus americanus*), Andean condor (*Vultur gryphus*), bald eagle (*Haliaeetus leucocephalus*), barn owl (*Tyto alba*), black-crowned night-heron (*Nycticorax nycticorax*), European perch (*Perea fluviatilis*), eastern and western Santa Cruz Island Galapagos giant tortoises (*Chelonoidis donfaustoi* and *Chelonoidis porteri*), eastern coyote (*Canis oriens*), fisher (*Pekania pennanti*), golden eagle (*Aquila chrysaetos*), northern pike (*Esox lucius*), raccoon (*Procyon lotor*), white-tailed deer (*Odocoileus virginianus*), and wood turtle (*Glyptemys insculpta*). For each device model, the selected error models and their goodness-of-fit statistics are reported in table 2, while the details of each individual model-selection procedure are provided in App. S4. With the following statistics on these 27 device models, we also report 95% binomial confidence intervals to make population-level inferences on devices of this era.

**Table 2:**
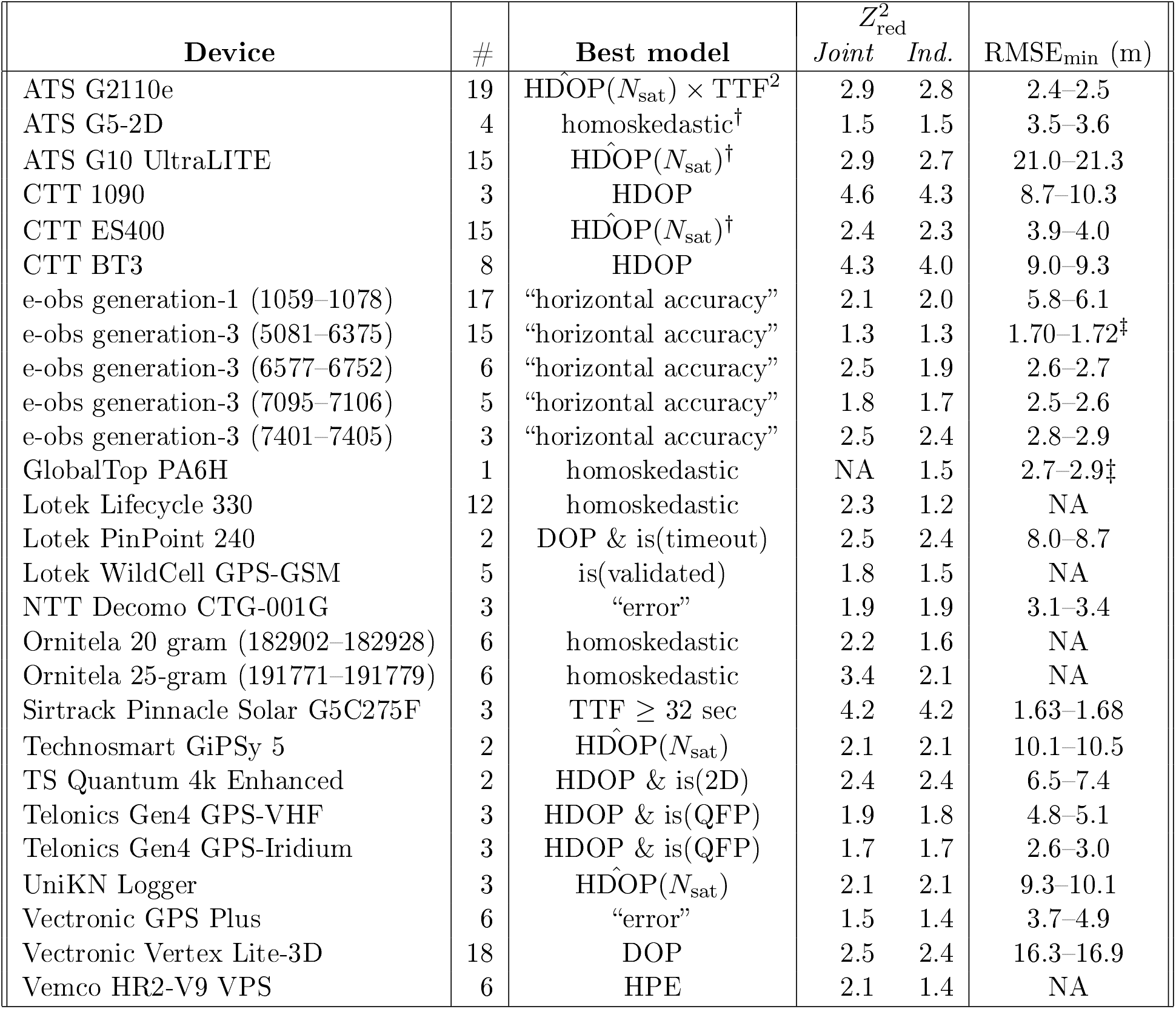
Selected horizontal error models, detailed in App. S4, with corresponding goodness-of-fit statistics 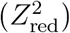 and root-mean-square location error given ideal reception (RMSE_min_) for 27 GPS and VPS device models, where ‘#’ denotes the number of devices sampled. The 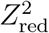 statistic assesses the performance of a device’s error model, where a lower value indicates more informative DOP values and a shorter tailed UERE distribution. 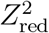 statistics are given for both the joint error model, assuming all devices of the same type are equivalent, and the individualized error model, assuming all devices of the same type have unrelated error parameters. If these two statistics are close, then error calibration can be assumed to be transferable among devices of this type. ^†^Performance of the selected model was not significantly better than the null HDOP model. ^‡^This performance was obtained in 1-Hz data.

In 56% (39%–75%) of device models, the reported data on location error were trustworthy enough that model selection was not necessary to obtain the best performing error model, if following the guidelines suggested in App. S3. This includes using the manufacturer supplied error estimates when provided, the most appropriate DOP values otherwise, and our number-of-satellites model (S3.1) as a last resort. In 77% (63%–9l%) of device models, one set of calibration parameters could safely be assumed valid for all devices of the same make and model. Only 7% (1%–24%) of device models had a selected error model that required substantial modeling effort. The Sirtrack Pinnacle Solar G5C275F had location errors that shrank by a factor of four after 32 seconds of “time on”, while the ATS G2110e had location errors that decreased smoothly with number of satellites and increased smoothly with time-to-fix. Both of these device models’ reported HDOP values that were either not informative or were marginally informative.

In terms of horizontal errors, an estimated 22% (9–42%) of device models had DOP values that were *misinformative*, in that they provided worse information on location error than a model of homoskedastic errors. All other properties held fixed, a larger 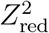 statistic indicates a heavier tailed UERE distribution. In several examples, the DOP values were so poor that their 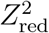 were comparable to that of a Cauchy distribution, which does not even have a finite variance. Furthermore, in terms of vertical errors, an estimated 0% (0%-52%) of device models had particularly informative VDOP values, in that they provided the best information on vertical location error, as they are expected to. Informative VDOP values are necessary to produce statistically optimal predictions in the vertical dimension, such as flight height distributions.

Finally, for each device we estimated its (horizontal) RMS location error under ideal conditions, which we denote as RMSE_min_. Ideal conditions typically amount to HDOP=1, in which case RMSE_min_ is equivalent to the RMS UERE calibration parameter, but some devices return DOP values below 1 or error estimates in meters rather than unitless DOP values. RMSE_min_ estimates ranged an order of magnitude, from 1.7 meters to 21 meters, with the more precise devices likely relying on newer dual-frequency GPS receivers or on-board Kálmán filtering. Similarly, RMS UERE calibration parameters also varied by an order of magnitude. Therefore, suggested DOP-value thresholds, like HDOP=10, can differ across device models by an order of magnitude in their physical implication. Furthermore, in the absence of calibration data or device specific knowledge, a blanket prior for GPS location-error parameters must span an order of magnitude (Fig. 1, method 2), which limits their capability and further motivates the use of calibration data (Fig. 1, method 1).

### 3.2 Error estimation of a GPS-tracked brown pelican

#### Methods

To demonstrate the benefit of an informative prior on location error parameters, when simultaneously fitting movement and error parameters in the absence of calibration data, we considered 5 months of GPS data, including HDOP values, collected at 90-minute intervals from a brown pelican (*Pelecanus Occidentalis*), tracked with a 65-gram Geotrak Solar GPS PTT. There were no associated calibration data. To this dataset we performed a nonresident continuous-time movement-model-selection procedure (Calabrese et al., 2016) with and without an informative prior on the location error parameters. When calibration date are unavailable, as is the case here, fitting with an informative prior corresponds to our proposed “fall-back” (Fig. 1, method 2). For the RMS UERE’s prior, we used a gamma distribution with 10-meter mean and 2 degrees-of-freedom, which has 95% credible intervals that are close to the range of estimates reported in Sec. 3.1. We then inspected the location-error estimates and corresponding occurrence distributions. Occurrence distributions quantify where the individual was located during the sampling period by probabilistically reconstructing the movement path (Fleming et al., 2016). The most common example of an occurrence distribution is the Brownian bridge (Horne et al., 2007), which assumes a movement model of Brownian motion. In contrast, here we selected a movement model via AIC. As the selected movement models here ultimately featured continuous velocities and a lack of range residence, we note that the final comparative analysis could similarly be performed with `crawl` (Johnson et al., 2008; Johnson, 2008), using the prior we obtain here, as `crawl` has the capability to specify a prior on the location-error parameter.

#### Results

When simultaneously fitting movement parameters without an informative prior, the RMS location error was estimated to be 780 meters (710–850 m) at HDOP=1, which is implausibly large for GPS location data, especially with an open view of the sky (Fig. 5A). In contrast, when employing an informative prior of 10 meters (1.6–19 m) at HDOP=1, based on the results of Sec. 3.1, the posterior location-error parameter estimate was 10 meters (1.4–20 m) at HDOP=1 (Fig. 5B), which was neither substantially nor significantly different from the prior estimate, indicating that these tracking data contained little information on the location-error process.

**Figure 5:**
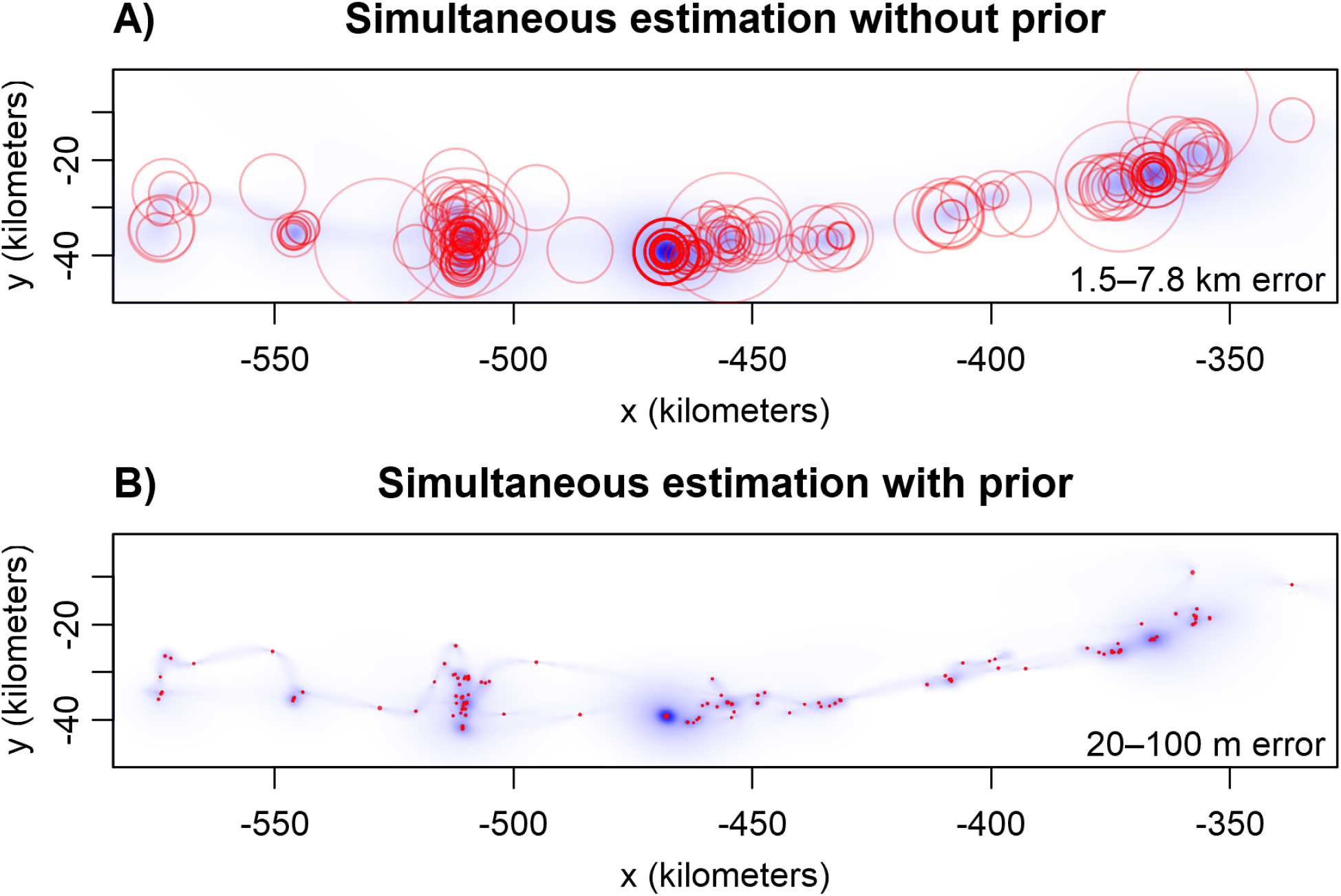
GPS data with 95% error circles (red) and occurrence distributions (blue) for a brown pelican (*Pelecanus Occidentalis*), as calculated from the AIC-best continuous-time movement model fit (**A**) simultaneously without a prior on the location-error parameters, and (**B**) with an informative prior on the location-error parameters. In each subpanel, the range of RMS location-error point estimates, corresponding to the error circles, are reported in the lower right corner of the plot. In this case, the scales of movement are much larger than typical GPS location errors and the simultaneously estimated location-error parameters in A are implausibly large.

### 3.3 Error estimation of a GPS-tracked common noddy

#### Methods

To demonstrate the impact of heteroskedastic location errors on the simultaneous fitting of movement and error parameters, we considered 4 days of common noddy (*Anous stolidus*) data, tracked with a custom GPS tag (Bouten et al., 2013) in 20-minute intervals. Locations were clipped from around the nest to isolate foraging behavior. For this dataset we performed a standard continuous-time movement-model-selection procedure (Calabrese et al., 2016) with four different error models—(A) a simultaneously fit homoskedastic location-error model, (B) a simultaneously fit PDOP-informed error model, (C) a PDOP-informed error model with an informative prior, and (D) a calibrated PDOP-informed error model. For each selected model we inspected the location-error estimates and corresponding occurrence distributions. Finally we cross validated the error-parameter estimate from each method against that of calibration data, which would not be subject to statistical identifiability issues. As the selected movement models here ultimately featured continuous velocities, and only a fine-scale analysis was performed, we note that a similar final comparison could also be made with `crawl` (Johnson et al., 2008; Johnson, 2008), by specifying the prior of the location-error parameter to be the likelihood of its calibration estimate, which we obtain here.

#### Results

When simultaneously fitting movement parameters with a homoskedastic error model, the RMS location error was estimated to be 500 meters (95% CI: 390–610 m), which, again, is implausibly large for GPS location data, especially with an open view of the sky (Fig. 6A). Simultaneously fitting movement parameters with a PDOP-informed error model improved the RMS location-error estimate by an order of magnitude to 44 meters (21–68 m) at PDOP=1 (Fig. 6B), but this estimate is still several times larger than expected, with a lower confidence interval that barely includes the largest minimum error reported in Sec. 3.1. We note that 44 meter location error at PDOP=1 may not seem large, but this implies 440 meter location error at PDOP=10, and DOP values tend to range from 1–10 or 1–20. By simultaneously fitting error and movement model parameters, these first two results correspond to the conventional application of state-space models. When including an informative prior, based on the results of Sec. 3.1, the RMS location-error estimate improved slightly to 37 meters (95% CI: 13–62 m) at PDOP=1 (Fig. 6B). Finally, when using a small sample of calibration data, we found the RMS location error to be only 3–4 meters at PDOP=1, which is consistent with a GPS device that has a dual-frequency GPS receiver or an on-board Kálmán filter. None of the simultaneously estimated location-error parameters cross-validated with the 3–4 meter calibration-derived estimate (*F*-test: *p*_A_ = 5.4 × 10^−72^; *p*_B_ = 2.9 × 10^−22^; *p*_C_ = 1.1 × 10^−16^), which avoids issues of statistical identifiability and movement-model misspecification. We note that this cross validation does not assume that the signal reception is comparable in the calibration and tracking datasets in any of the PDOP-informed error models (B–C). Estimation performed with the integrated (calibration — tracking data) model D did not significantly update the 3–4 meter minimum error estimate (Fig. 6C), which indicated that the tracking data had little information regarding the location-error process.

**Figure 6:**
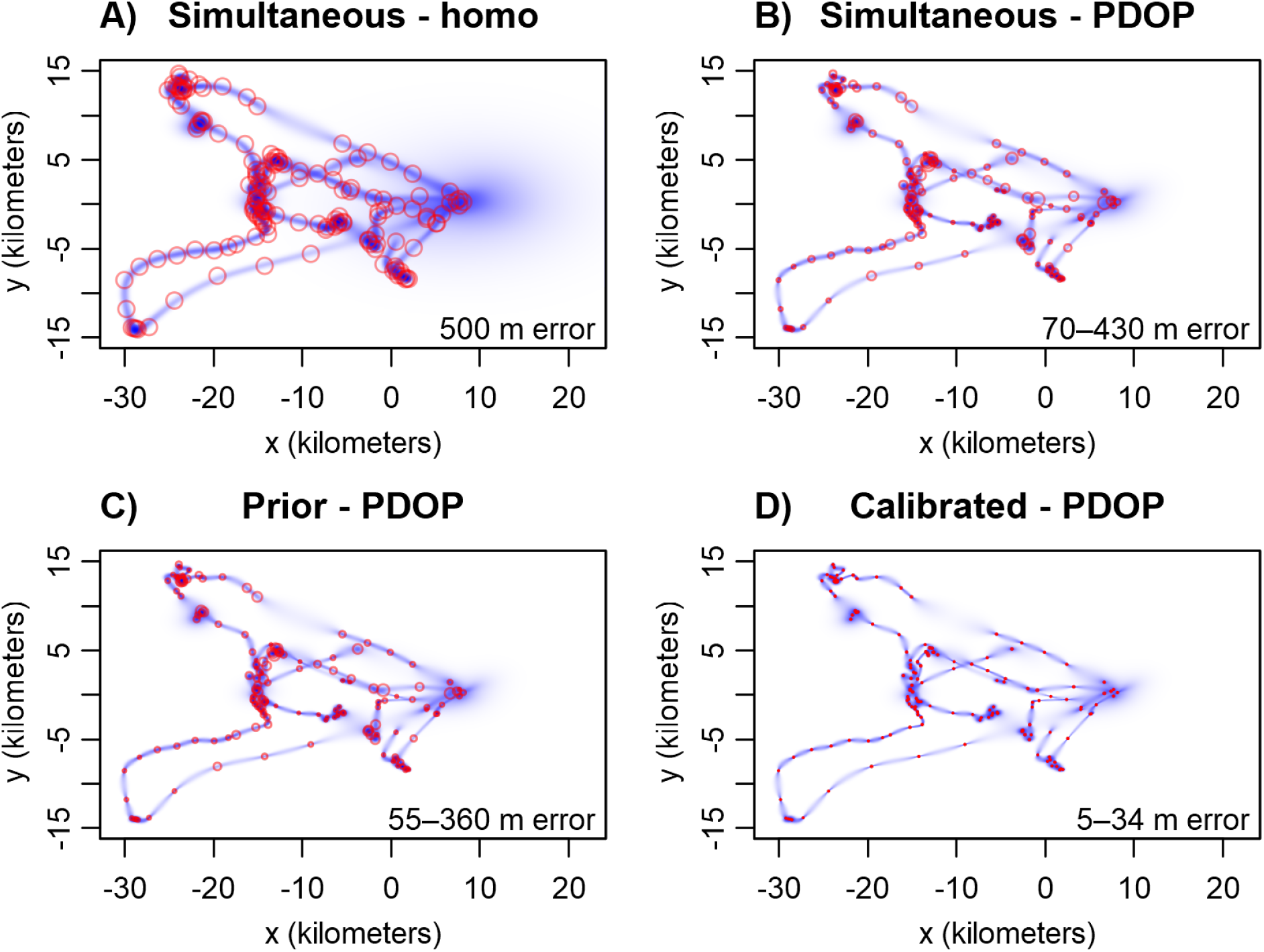
GPS data with 95% error circles (red) and occurrence distributions (blue) for a common noddy (*Anous stolidus*), as calculated from the AIC-best continuous-time movement model fit (**A**) simultaneously with a homoskedastic location-error model, (**B**) simultaneously with a PDOP-informed error model, (**C**) with a prior-informed PDOP error model, and (**D**) with a calibrated PDOP error model. In each subpanel, the range of RMS location-error point estimates, corresponding to the error circles, are reported in the lower right corner of the plot. Subpanels **A–B** represent the conventional application of state-space models, which simultaneously fit movement and error model parameters, while subpanel **D** represents the calibration approach we advocate and subpanel **C** represents the fall-back we suggest if no calibration data are available (Fig. 1, method 2). In this case, the scales of movement are much larger than typical GPS location errors and the simultaneously estimated location-error parameters in **A** are implausibly large. In **B**, structure is provided to the location-error model that allows for improved parameter estimates, though the results are only marginally credible, and then only marginally improved by an informative prior in **C**. None of the uncalibrated location-error estimates cross validate with the calibration data, which is leveraged in **D** and provides an estimate that is 1–2 orders of magnitude smaller than that of **A–C** under the same conditions.

### 3.4 Speed of GPS-tracked wood turtle

#### Methods

To demonstrate the impact of error-informed analysis on fine-scale inference, we considered 6.3 months of wood turtle (*Glyptemys insculpta*) data, tracked with a Lotek GPS tag in 1-hour intervals, for which we apply the location-error model selected in Sec. S4.5.2. We applied two mean-speed estimators: a conventional straight-line-distance estimator and a continuous-time speed and distance estimator that accounts for both location error and tortuosity (Noonan et al., 2019). For each estimator, we conditioned our estimates on data with HDOP values below a specific threshold, which we manipulated.

#### Results

While the straight-line distance estimates were extremely sensitive to the HDOP cutoff, the error-informed estimates were largely consistent across threshold levels (Fig. 7). With the conventional straight-line distance estimator, two primary sources of bias exist—positive bias from ignoring location error (Ranacher et al., 2016; Noonan et al., 2019) and negative bias from ignoring tortuosity (Rowcliffe et al., 2012; Noonan et al., 2019). In contrast to homerange estimation, discarding erroneous observations to mitigate against the first bias actually increases the second bias. Therefore, while in Fig. 7 the mean-speed estimate is improved by discarding erroneous data, in other cases with faster species the same manipulation can produce an increasingly net negative bias. Importantly, the “Goldilocks”—not too much, not too little—HDOP threshold required to balance these two biases cannot be known a priori, as it depends sensitively on the scales of movement, scales of location error, and sampling schedule.

**Figure 7:**
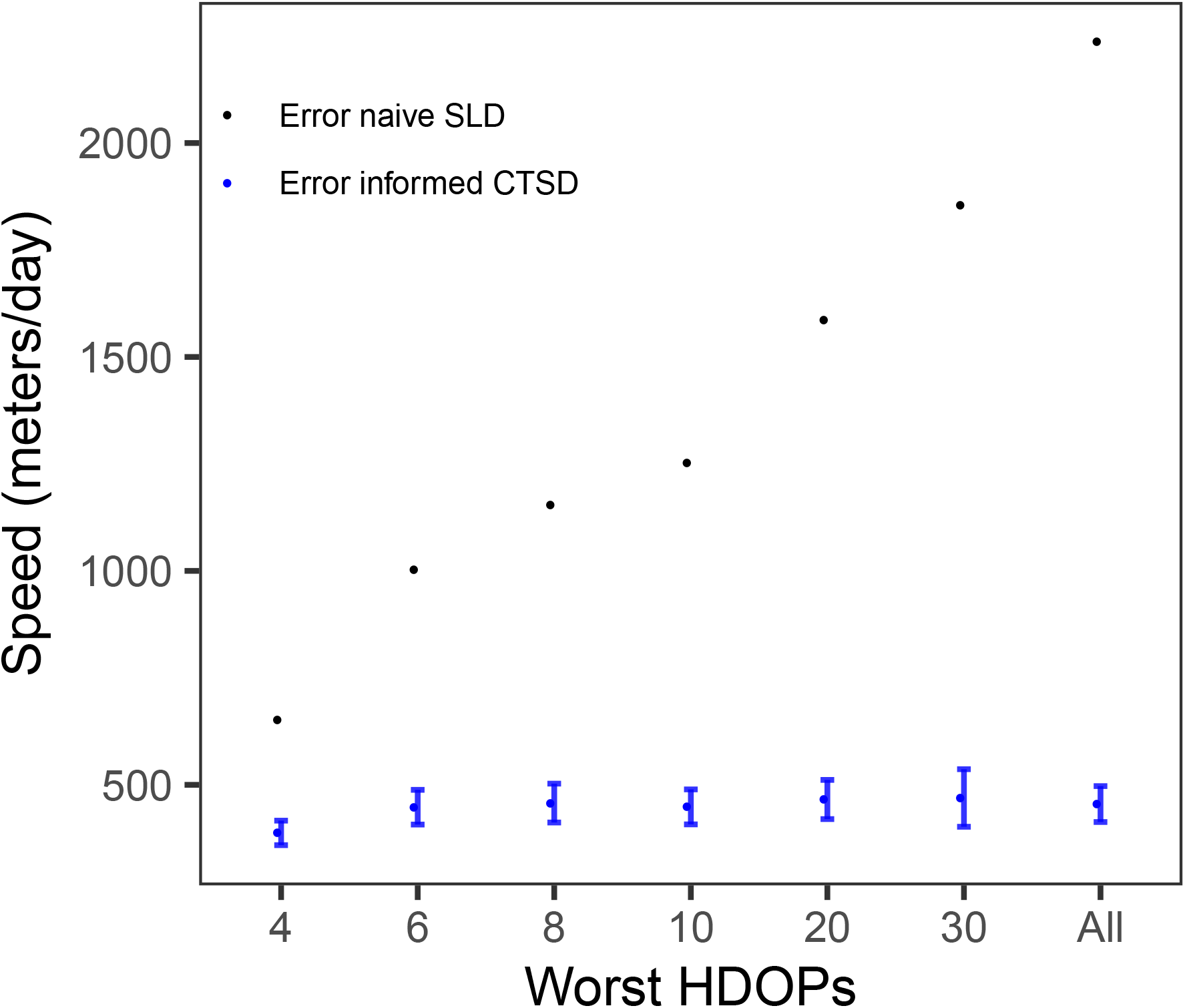
Mean speed estimates with 95% CIs (where available) for a GPS-tracked wood turtle versus the worst horizontal dilution of precision (HDOP) value included in the data. The black points correspond to conventional straight-line distance (SLD) estimates and the blue points correspond to an error-informed continuous-time speed and distance (CTSD) estimator based on time-series Kriging (Noonan et al., 2019). The first estimate only includes location fixes with HDOP ≤ 4, while the final estimate includes all of the data. Only the error-informed estimates are consistent across all included HDOP values, though the lowest HDOP locations will be biased towards open habitats.

### 3.5 Home-range area of an Argos-tracked night-heron

#### Methods

To demonstrate the impact of error-informed analysis on coarse-scale inference, we considered 6 months of data on a black-crowned night-heron (*Nycticorax nycticorax*), tracked with a Telonics TAV-2630 Argos Doppler-shift tag at ~60-minute intervals during its range-resident period—wherein the heron was not migrating and had a well established home range. These Argos Doppler-shift data were of the older variety that do not come pre-calibrated. Instead, each location estimate was accompanied with a location class—3,2,1,0,A,B,Z—for which we used the calibration results of Vincent et al. (2002) to approximate the error ellipses.^3^ We applied two home-range estimators, a conventional kernel density estimator (KDE) and an error-informed autocorrelated kernel density (AKDE) estimator with weights optimized to mitigate against irregular sampling (Fleming et al., 2015; Fleming and Calabrese, 2017; Fleming et al., 2018). In the latter method, both the autocorrelation estimator and bandwidth optimization tease apart variance due to movement and error, and the data are first Kriged to reduce location error before placing the kernels (Fleming et al., 2016, 2017). The estimator represents the default output of the `akde()` method in the `ctmm R` package when conditioning on an error-informed movement model. Moreover, for these data, AKDE without a location-error model also results in conventional KDE, because these data appear independent when ignoring error. Therefore, our comparison effectively includes both error-informed and non-informed AKDE.

With each estimator, we manipulated the quality of input data, from only including the best location class (3) to including all of the data (3-B). Researchers with Argos Doppler-shift data typically will threshold their data class-wise like this based on quality before applying methods that do not account for error. Therefore, our analysis tests the sensitivity to this choice of cutoff, in addition to biases that stem from not accounting for location error.

#### Results

The conventional KDE home-range area estimates were highly sensitive to the location-class threshold, and in each case were larger than error-informed estimates even when restricted to the highest quality data (Fig. 8). In contrast, error-informed AKDE produced consistent home-range area estimates, regardless of which location classes were included, though the effective sample size dropped precipitously when only including the highest quality fixes.

**Figure 8:**
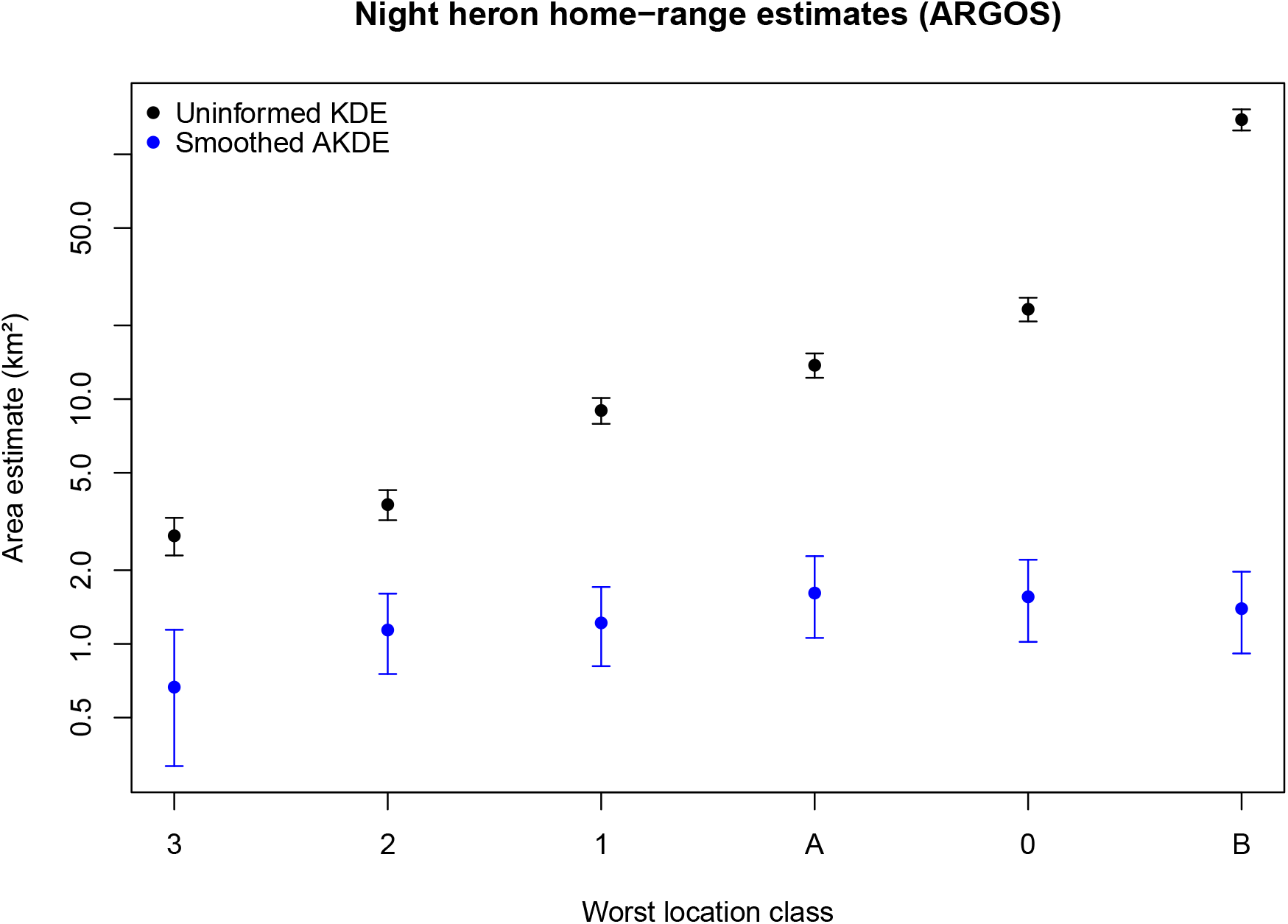
95% home-range estimates with 95% CIs for an Argos-tracked night-heron versus the worst location class included in the data, with less-accurate location classes censored for that estimate, where the black points correspond to conventional kernel density estimates (KDE) and the blue points correspond to error-informed autocorrelated kernel density estimates (AKDE). The Argos location classes 3,2,1,A,0,B are sorted by accuracy, and so the first estimates at ‘3’ only include data from the most accurate location class, and therefore have the smallest sample size, while the final estimates at ‘B’ include data from location classes 3-B, and are the largest when not accounting for location errors. Only the error-informed estimates are consistent across included location classes, though the effective sample size drops precipitously when only including location class 3. Note the logarithmic scale on the *y*-axis.

### 3.6 Home-range overlap of an Argos-tracked night-heron

#### Methods

To demonstrate the downstream impact of error-informed analysis on coarse-scale inference, we considered the degree of home-range overlap for a night-heron across two summers (2017, 2018), also tracked with a Telonics TAV-2630 Argos Doppler-shift tag, but complete with precalibrated error-ellipse information. We applied both conventional KDE and error-informed AKDE, and compared overlap (Winner et al., 2018) between summer ranges as a measure of year-to-year site fidelity. Again, these data appear independent when ignoring location error, and so AKDE without an error model results in conventional KDE. Therefore, our comparison effectively includes both error-informed and non-informed AKDE.

#### Results

According to the conventional KDEs, which is biased by location error to have inflated variance and thus inflated overlap, the 2017 and 2018 summer ranges were largely similar at 82% (78%-86%) overlap, whereas according to the error-informed AKDEs the 2017 and 2018 summer ranges were adjacent, but largely distinct, at 33% (22%–46%) overlap (Fig. 9). Therefore, in this case, the estimator used has a direct impact on biological inference, because the Argos Doppler-shift location errors are large relative to the home-range area of the nightheron.

**Figure 9:**
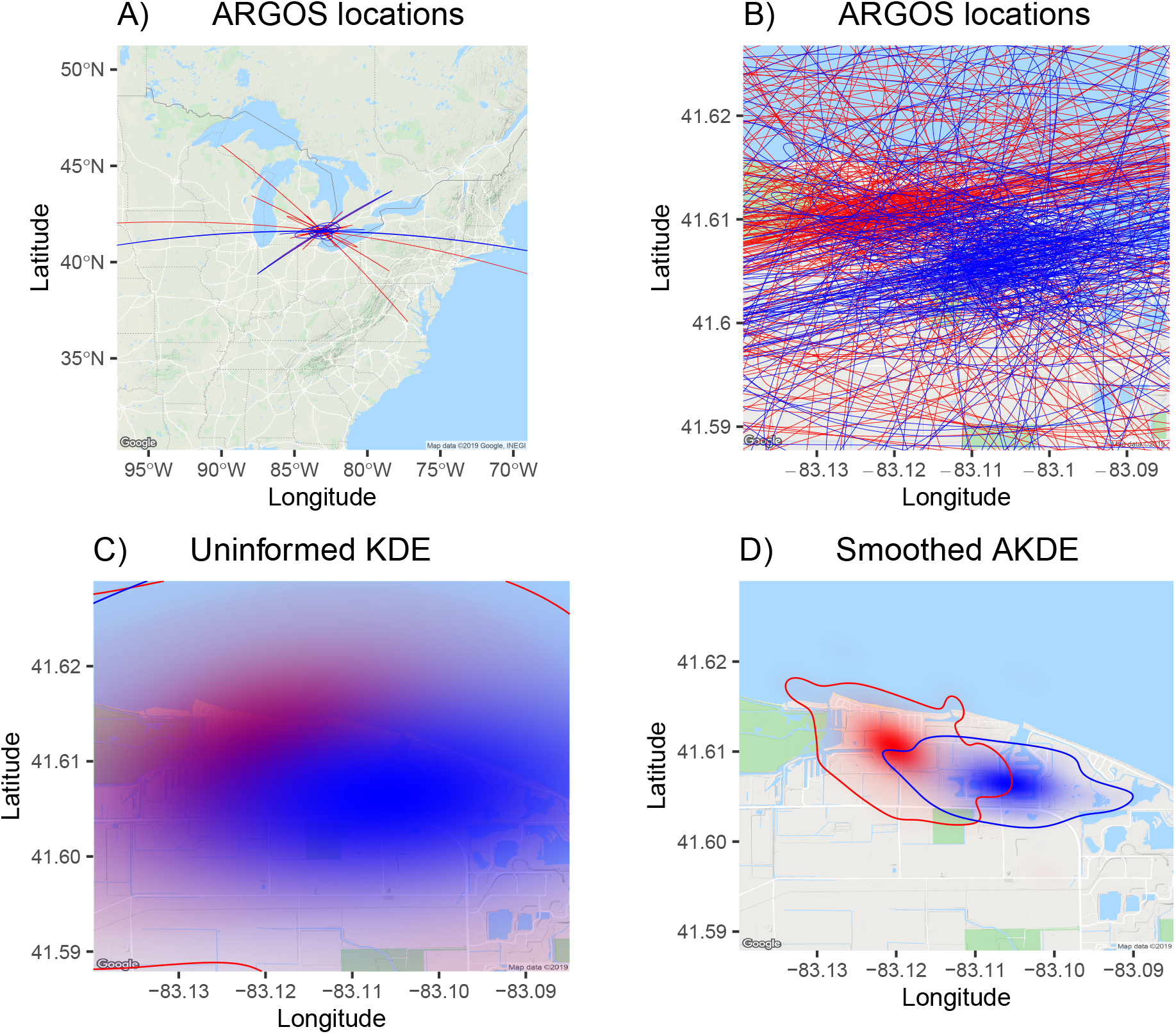
2017 (red) and 2018 (blue) summer ranges of a night-heron, as given by (**A**) Argos Doppler-shift error ellipses zoomed out to their largest location-error scale, (**B**) the same Argos error ellipses zoomed in to the inferred home-range scale, (**C**) conventional KDE home ranges, and (**D**) error-informed AKDE home range. All contours depict the 95% coverage areas. The conventional KDE estimates are inflated from location error, which makes the two summer ranges appear more similar than they are.

## 4 Discussion

We are currently in a golden age of biotelemetry, where more individuals and more taxa are being tracked than ever before (Kays et al., 2015). Tracking data have become highly desirable for a variety of ecological, evolutionary and conservation-related purposes. That is why it is important to use methods that can best make use of all of the information contained in tracking data without incurring spurious bias. A key component of that is properly accounting for location error, even with GPS data (Noonan et al., 2019). The widespread practice of discarding location fixes based on a DOP-value or location-class threshold can only serve as a partial mitigation against location error, at the cost of diminished sample size. Using empirical data, we have demonstrated that error-naive estimates can be very sensitive to the chosen threshold, and cannot be expected to converge to error-informed estimates, even when most of the data has been discarded (Figs 8–7). Without an appropriate error model, common movement metrics can easily be biased by an order of magnitude (also see Noonan et al., 2019). We have also demonstrated that device-specific errors can vary by an order of magnitude (Table 2), which means that a DOP value of 1 on one model device can be equivalent to a DOP value of 10 on another model device. On the other hand, simultaneously estimating both the movement and location-error parameters, which is also a common practice, is known to be problematic because the variances due to movement and to error cannot be reliably distinguished (Auger-Méthé et al., 2016; Noonan et al., 2019).

Motivated by these challenges, we have introduced a comprehensive guide and framework for addressing location error in animal tracking data. Our framework consists of a three-step solution to account for location error in movement analysis (Fig. 1) starting with the collection of calibration data and the application of formal model-selection techniques to construct a statistically efficient location-error model. Finally, with error-informed movement analyses, researchers can use all of their data to produce the best possible estimates. We have also added to the existing number of error-informed analyses with statistics for outlier detection and autocorrelation estimation.

Error-informed analysis on calibrated tracking data can be necessary for meaningful biological inference, as the amount of bias present in conventional estimates is a function of the sampling schedule, scales of movement, and scales of error. When making comparisons across time, across individuals, or across taxa, observed differences in estimated behavior may simply be the result of differential biases, when using traditional methods. For instance, if estimating energy expenditure via distance traveled by conspecifics at two different study sites, presumed differences could be the result of environmental factors driving hypothesized biological mechanisms, or there could a be non-biological explanation related to inaccuracy. Differences in distance estimates could be the result of two different model tracking devices with substantially different calibration parameters having been deployed at the two study sites, so that the individuals tracked with less precise receivers appear to move more tortuously and travel for longer distances. Differences in distance estimates could also be the result of the study sites corresponding to different habitats with inconsistent satellite reception, such as an open grassland versus a canopied woodland, and so the individuals that spend more time in woodland habitat appear to move more than their grassland counterparts. Unfortunately, there is no sure way to discriminate these possibilities with commonly used methods. Furthermore, because these biases also depend on the animals’ movement characteristics, it not generally sufficient to make error-naive comparisons even if the tracking device, sampling schedule, and habitat are all identical (e.g., Noonan et al., 2019, Fig. 4c). Differential bias is especially likely to happen in repeated studies over time, as older devices are replaced by newer device models, and in studies where different researchers collected data in different habitats and with different tracking devices. As movement data continue to amass and larger, collaborative studies become increasingly possible, this issue will only increase in importance. For instance, Morato et al. (2016) compared the movement behaviors of 44 jaguars across 5 biomes with 5 different model GPS tracking devices. Combinations of GPS and Argos Doppler-shift tracking data are also common in marine systems (e.g., Phalan et al., 2007; Block et al., 2011; Raymond et al., 2015; Citta et al., 2018).

As location data may be applied to a variety of purposes in ecology, we suggest that the collection and analysis of GPS calibration data needs to become standard practice for researchers and managers in the short term. The error-model-selection framework we have introduced here consists of statistically efficient parameter estimators, AIC_C_ formulas, and a goodness-of-fit statistic 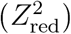—all custom-derived for working with error-laden location data. As we have shown in App. S4, GPS tracking devices output a variety of data related to location error and can exhibit a variety of error structures. At this time, model selection is often necessary to ensure that location error is accurately described, but consumer pressure on manufacturers could ease this burden considerably, as we detail below.

For existing and near-future tracking data, it would also be useful to have a repository of calibration data, sorted by device make and model, for researchers to draw from. For many historical datasets, the original tracking devices no longer exist or are no longer functional, and even today the choice to collect adequate calibration data can sometimes come at the cost of tracking fewer individuals. Our curated calibration dataset (Fleming et al., 2020), featuring 190 devices comprising 27 models from 14 manufacturers, is a first start toward this goal.

### Manufacturer Recommendations

We suggest that the task of calibration and error-model selection (Fig. 1) be performed by GPS device manufacturers, as is the case with Argos Doppler-shift horizontal errors, by reporting 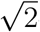-standard-deviation error circles or equivalent. At a minimum, all error information from the GPS module should be returned and calibration data should be provided— preferably per device and at different sampling rates if on-board error filtering varies. Moreover, complete information on version and build for all software, firmware, and hardware, are also necessary to know which calibration parameters can be expected to be applicable across devices.

On a lower level, we note that Argos offers both Kálmán-filtered Doppler-shift location estimates and unfiltered (least-squares) location estimates. This kind of choice would also be appealing to GPS consumers. Researchers that process their own data with software packages such as `crawl` and `ctmm` can potentially obtain better precision with unfiltered data, instead of processing the same data twice over. Error ellipses would also be beneficial for GPS tracking data in some cases and GSM tracking data in most cases.

### Device performance

It is straightforward to answer the question of which GPS devices have the smallest errors in a given environment. However, such a comparison would be limited to the specified environment. Our 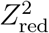 goodness-of-fit statistic provides a different comparison that addresses which device has the more predictable distribution of location errors. In other words, a small 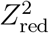 indicates that the device gives fair warning of large errors, with few false alarms. Moreover, 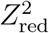 does not require that environmental conditions be controlled for, and so these statistics can be compared more generally. Smaller values of 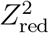 indicated better error-model (e.g., DOP-value) performance, and 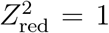 results when the location errors are properly calibrated and the resulting UEREs are normally distributed, which is not an unreasonable assumption (Kaplan and Hegarty, 2006). In general, less informative DOP values will fail to explain heteroskedasticity and produce heavier tailed UERE distributions and larger values of 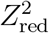. For the 27 GPS, cellular, and VPS device models we analyzed, the AIC_C_-best error models produced 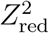 statistics ranging from 1.3–4.2, with a median value of 2.3 (Table 2).

Beyond the 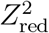 statistic, at least two other measures of error performance are necessary for a comprehensive comparison—the RMS location error under ideal conditions (RMSE_min_) and a standardized measure of DOP-value response to satellite-signal attenuation. Though the calibration data we analyzed were not collected for this task, we can report RMSE_min_ values between 1.7 and 21 meters (Table 2), with the smaller location errors likely being the result of dual-frequency GPS receivers and multiple location fixes processed by an onboard Kálmán filter. Specifically, devices may condition the current location estimate on past data, under an assumed movement and location-error model, possibly also leveraging Doppler-shift information. Some devices filter all location estimates, keeping the constituent data unreported at coarse sampling rates, while other devices only apply filtering when the sampling rate reaches a specific threshold (~1 Hz). On this point we also caution researchers that devices with on-board filtering may have smaller RMS UERE parameters at higher fix rates, and, as such devices warm up, the magnitude of these parameters can continue to shrink until they reach an asymptote.

For 22% (9-42%, population CI) of tracking device models, their recorded DOP values were misinformative, in that they provided worse information on location error than a simple model of homoskedastic errors, and so using them could actually have a deleterious effect on analysis. Because informative DOP values are necessary to account for the many sources of heteroskedasticity in tracking data, they can be important in many situations, such as if animals traverse through habitats of differing satellite reception. For instance, animals that burrow or rest under trees might appear to be scattering about ~100 meters when not moving, without informative DOP values to explain the increase in variance. In these cases, researchers should attempt to collect calibration data in different environments, as an approximate correction, but such calibration data are unlikely to exactly reproduce the same quality location estimates as the animal, and habitats may differ between the location estimate and the true location.

Finally, we note that *none* (0%–52%) of the 25 GPS tracking devices we analyzed had particularly informative VDOP values, where the null model based on VDOP outperformed models using HDOP values or our number-of-satellites model (S3.1) as proxies for vertical error, or even a homoskedastic model. Informative VDOP values are necessary for an accurate assessment of vertical errors, and 12 of the devices we tested were GPS tags, rather than collars, which would be applied more often to aerial or arboreal species. These data are increasingly needed in good quality, considering human impacts in the airspace, such as wind farms, buildings, planes, power lines, etc. (Lambertucci et al., 2015; Ross-Smith et al., 2016; Péron et al., 2017; Poessel et al., 2018a).

### Error model performance

Particularly with older Argos Doppler-shift location data, which only feature location class information (3,2,1,0,A,B,Z), several analyses have employed *t*-distributed location errors, as the error distributions within location classes are heavy tailed (Jonsen et al., 2005; Hoenner et al., 2012; Albertsen et al., 2015; Brost et al., 2015). Error models that fail to capture the heteroskedasticity of satellite location data will generally be heavier tailed (c.f., Fig. S4.2). This holds for both historical Argos Doppler-shift location data and GPS location data with inadequate error-model performance (e.g., data that lack informative DOP values). While *t*-distributed errors are readily available in `R` package Template Model Builder (`TMB`; Albertsen et al., 2015; Péron et al., 2017; Auger-Méthé et al., 2019), heavy-tailed error distributions eventually need to be incorporated into tracking specific `R` packages, such as `crawl` (Johnson et al., 2008; Johnson, 2008) and `ctmm` (Fleming and Calabrese, 2015; Calabrese et al., 2016). As pointed out by Albertsen et al. (2015), there are straightforward and reliable approximations for heavy tailed error distributions implemented in `TMB`, which can also be implemented in similar frameworks like `crawl` and `ctmm`.

On the other hand, technological advances, such as solar-powered GPS tags, now allow very high sampling rates to be sustained for long periods of time (App. S4.3.2). These new data introduce several complications when it is necessary to account for location errors. First, as the sampling interval falls below 1–2 minutes, GPS location errors become autocorrelated (Tao and Bonnifait, 2015). Second, many GPS devices employ on-board filters to post-process high-frequency location estimates and shrink their errors, which comes at the cost of increased location-error autocorrelation. As these types of data become increasingly common, the calibration models and estimators introduced here will need to be upgraded from independently sampled errors to autocorrelated errors. Otherwise, fine-scale prediction uncertainties, such as those of location and speed, will be underestimated when analyzing high frequency tracking data. The accommodation of location-error autocorrelation may also be necessary for VPS data (App. S4.14.1), though further testing is warranted. Generalizing the methods introduced here to incorporate autocorrelated location errors is a largely straightforward, though tedious, computational task.

When using the independent and identically distributed (IID) UERE methods presented here on high-resolution GPS tracking data, we can approximate the resulting biases as follows. If Δ*t* is the sampling interval, then effective sample sizes are overestimated by an inflation factor on the order of I ≡ (1 min)/Δ*t*. AlC_C_ differences will be overestimated by a factor on the order of *I*, which will overstate differences in model performance if not accounted for. UERE variance parameters will be relatively underestimated by a factor on the order of *I/N*, where *N* is the nominal sample size assuming independence. However, this bias is quite minor in calibration datasets that sample for an hour or more. In any case, we recommend collecting calibration data over the course of a day, if not many days.

### Simultaneous estimation when necessary

The simulation example considered by Auger-Méthé et al. (2016) represents a more extreme case of lack of identifiability between relatively featureless, discrete-time movement and error models, though model misspecification can produce worse outcomes. If estimating movement and location error simultaneously, we recommend (1) using an informative prior to penalize implausible error-parameter estimates and (2) enforcing as much distinguishing structure as possible on the two processes to increase identifiability in their parameters. First, as we have demonstrated with both simulation (Fig. 2) and empirical data (Fig. 5), if the scales of movement and location error are very different, then simultaneous estimation can produce wildly inappropriate outputs, where both processes model the larger variance. An informative prior, such as based on the results of Sec. 3.1, can prevent these grossly implausible estimates. However, the order-of-magnitude variation in GPS device performance (Table 2) makes calibration data preferable. Second, there are a variety of ways that movement model and location-error model structure can differ. One example of distinguishing structure is the heteroskedasticity of location error, as captured by informative DOP values (or calibration-derived relative variances, as in Jonsen et al., 2005). It is unlikely that an individual’s finescale displacements will scale with DOP values as well as location error, and therefore homoskedastic errors, as simulated in Auger-Méthé et al. (2016), are much more vulnerable to issues of identifiability than heteroskedastic errors (Fig. 6). Another feature that distinguishes the movement and error processes are their contrasting autocorrelation structures. GPS location errors are typically not substantially autocorrelated at sampling intervals > 1-2 minutes (Tao and Bonnifait, 2015), while the movement processes of large mammals are typically correlated for days to weeks (McNay et al., 1994; Rooney et al., 1998; Boyce et al., 2010; Fleming et al., 2014*a,b*; Morato et al., 2016). Because of this separation of scales, irregular sampling is helpful for teasing apart movement from error, given an appropriately autocorrelated movement model. In contrast, neglecting autocorrelation or shoehorning irregularly sampled data into a discrete-time framework will result in a loss of this useful information. A more extreme example of distinguishing structure is found in the practice of double tagged GPS and Argos Doppler-shift tracking data (Winship et al., 2012), which can function almost identically to calibration on the Argos Doppler-shift location estimates if the GPS locations are sampled near to the Argos Doppler-shift locations and the GPS location errors are able to be treated as insubstantial (and not simultaneously estimated).

### Error-informed analyses

The number of movement analyses based on continuous-time state-space models, which can allow for both irregular sampling and location error, is steadily expanding (e.g., Breed et al., 2017; Michelot and Blackwell, 2019; Jonsen et al., 2019). However, there remain several analytic tasks that have yet to be formulated in a way that accounts for location error. Primary among these tasks are resource selection and home-range estimation. Resource selection with uncertain location data is an outstanding yet straightforward challenge that can be addressed by incorporating an observation model and marginalizing over the unknown true locations. Non-parametric home-range estimation is less obvious. The more general problem of kernel density estimation with measurement error is conventionally approached by ‘deconvolving’ or de-smoothing the kernels according to the spread of the error (Stefanski and Carroll, 1990). However, it is straightforward to show that this approach will not generally produce an asymptotically consistent estimator, meaning that the estimate will not converge to the truth in the limit of infinite data, as the individual kernels remain biased in their spread. The method we applied here, which involves Gaussian smoothing of the data (Fleming et al., 2016) and error-informed bandwidth optimization, is relatively ad hoc but is ensured to produce a density estimate of appropriate scale. Future home-range estimators will need to both optimally weight the data, in the sense of Fleming et al. (2018), so that more erroneous location estimates are treated as less informative, and smooth the data conditional on the kernel density estimate, rather than on a normal approximation of the density function. The latter is needed to shrink the estimate in towards the density of the data, rather than, simply, the mean of the data. Finally, we note that the error-informed statistics for outlier detection introduced in App. S5 assume circular error distributions and could be improved for Argos Doppler-shift, triangulated VHF, and light-level geolocator-derived data that have more elongated error distributions.

User-friendly `R` software packages that can incorporate the kind of heteroskedastic location errors we model here include `crawl` (Johnson et al., 2008; Johnson, 2008), `ctmm` (Fleming and Calabrese, 2015; Calabrese et al., 2016), `move` (Kranstauber et al., 2012), `momentuHMM` (McClintock and Michelot, 2018), and `smoove` (Gurarie et al., 2017*b*). Other packages capable of performing simultaneous estimation, such as `foieGras` (Jonsen et al., 2019), could also be easily updated to allow for priors on the location-error parameters. The methods and models we have introduced are implemented in the `R` package `ctmm`, complete with long-form documentation `vignette(“error”)`. Based on the calibration results presented here (Apps. S2–S4), error model selection is obviated in `ctmm` for Argos Doppler-shift, e-obs, Telonics, Vemco, and many other device models, as we have tested the performance of their standardized error information. Going forward, we envision both error modeling and error-informed animal movement analysis becoming more necessary, more user friendly, and more accurate.

### Final remarks

Historically, GPS location error has rarely been modeled and accounted for, because the magnitude of these errors have often been negligible, relative to the scales of movement. However, improvements in the temporal resolution of tracking data, device miniaturization for smaller taxa (Kays et al., 2015), and an increase in multi-study analyses (Tucker et al., 2018, 2019; Noonan et al., 2020) all escalate the need for reliable statistical methods that can account for location error. We have presented a major step towards achieving this goal, by deriving a comprehensive framework—from experimental design to error-model selection, error-model performance evaluation, and error-informed analyses—with the capability to pin down the observation model of animal tracking data and account for its heteroskedastic errors. Our framework allows tracking data to more directly and more accurately inform ecological analyses, now and going forward. This work should provide strong guidelines for both researchers and manufacturers in wildlife ecology and conservation biology, as well as other fields where data represent processes of interest and error processes that are difficult to tease apart.

## Acknowledgments

We thank Dr. Todd E. Katzner for his many contributions to and comments on this manuscript. We thank Drs. Edward Hurme and Arliss J. Winship for their many comments on this manuscript. CHF, JMC, MJN, and WFF were supported by NSF ABI 1458748. CHF, JMC, and WFF were supported by NSF IIBR 1915347. MMC and RAM were supported by MN DNR SWG F14AP00028. NPG and CSD were supported by the Pittman-Robertson Federal Aid to Wildlife Restoration Grant, NCWRC and the NCSU Fisheries, Wildlife, and Conservation Biology program. ALH was supported by the FONZ Conservation Research Grant, ConocoPhillips Charitable giving Global Signature Program, and MD DNR. RK was supported by NSF IIBR 1914928. KNP was supported by the Winifred Violet Scott Charitable Trust. AR and RS were supported by SNSF grant 173178. KS and CMT were supported by Laura Kearns, the Ottawa National Wildlife Refuge, the Winous Point Marsh Conservancy, and the ODNR Division of Wildlife. Any use of trade, product, or firm names is for descriptive purposes only and does not imply endorsement by the U.S. Government.

## S1 Sensitivity analysis

### S1.l Sensitivity analysis via simulation of errors

If 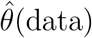 is the output of some analysis conditioned on the data that does not account for location error, and 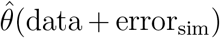 is the output of the same analysis, but with additional simulated location error added to the original data (atop its unknown location error), then the second-order variance induced by location error is simply the variance of the estimates 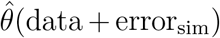, with respect to the ensemble of simulations, and is additive. The second-order bias induced by location error is given by the difference in the average simulated analysis and the unaltered analysis

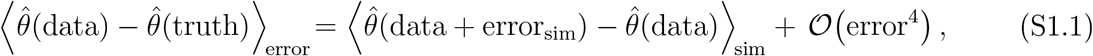

where 〈…〉_*X*_ denotes the expectation value with respect to the process *X* data = truth + error, and 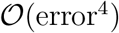 denotes higher-order corrections, including VAR [error]^2^, which require an error-informed analysis to determine. The only assumptions made here are that 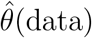 is analytic in the neighborhood of 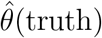, and that the distribution of errors is symmetric, which includes the errors being of mean zero.

### S1.2 Sensitivity analysis via simulation of trajectories

As a slight improvement, here we upgrade the previous analysis from estimates based on 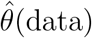 to estimates based on 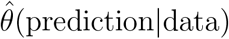, where ‘prediction’ refers to the Kriged location estimates, which are conditioned on the data and fitted movement model (Fleming et al., 2016). The second-order variance induced by location error is the variance of the estimates 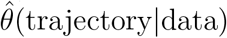, where ‘trajectory’ refer to simulated locations conditioned on the data and fitted movement model, which are centered on the conditional predictions (Fleming et al., 2017). The second-order bias induced by location error is given by the difference in the average simulated analysis and the analysis based on the Kriged estimates

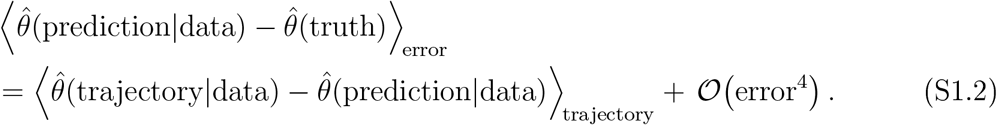

## S2 Error-model statistics

### S2.1 Single-UERE estimation

Here we derive the minimum variance unbiased (MVU) estimator for the mean-square UERE, (UERE^2^), given calibration data where the GPS tag/collar is at rest. Lack of bias and low variance is important so that multiple (possibly opportunistic) calibration datasets can be safely pooled regardless of size. Achieving the MVU property requires an assumption of normality in the distribution UEREs, which can still allow for a heavy-tailed distribution of location errors. For more general UERE distributions, these estimators are still unbiased.

In two dimensions, the RMS UERE, 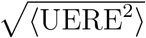, serves as the proportionality constant relating RMS location error and HDOP value

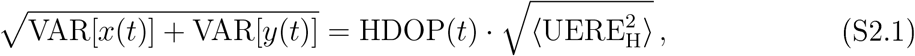

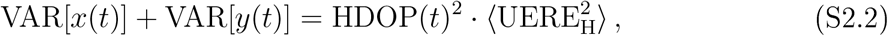

where UERE_H_ denotes horizontal UERE values. As GPS satellite coverage is relatively uniform, GPS location errors will generally be circular on average, in contrast to Argos Doppler-shift errors where there are only a few satellites in polar orbits. There are exceptions to the assumption of circular errors (Fig. S2.1), but GPS tracking devices do not record error-ellipse information, so there is little that can be done in these cases.

Given circular location errors, we have

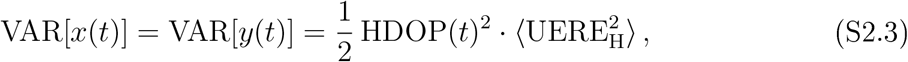

and similarly for the vertical dimension we have

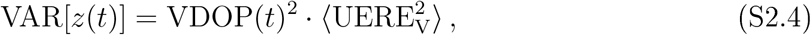

where in practice the horizontal and vertical RMS UERE values are not equal.

Let *k* index the *K* calibration datasets, with each corresponding to identical tags/collars at fixed locations ***μ**_k_*. The MVU mean-location estimators are then given by the weighted average

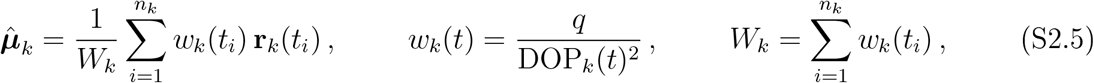

where **r**_*k*_(*t*) = (*x_k_*(*t*), *y_k_*(*t*)) in two dimensions. From hereon we keep our relations general with *q* = dim(**r**) denoting the number of spatial dimensions (*q* = 2 for horizontal and *q* = 1 for vertical. The MVU mean-square UERE estimator is

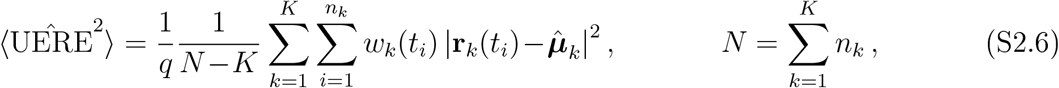

given *N* total sampled times for the K calibration datasets. The sampling distributions are given by

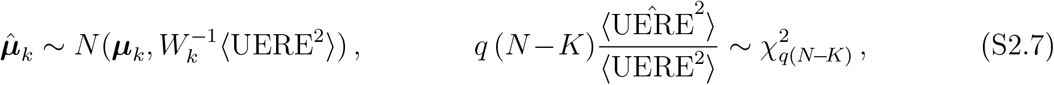

**Figure S2.1:**
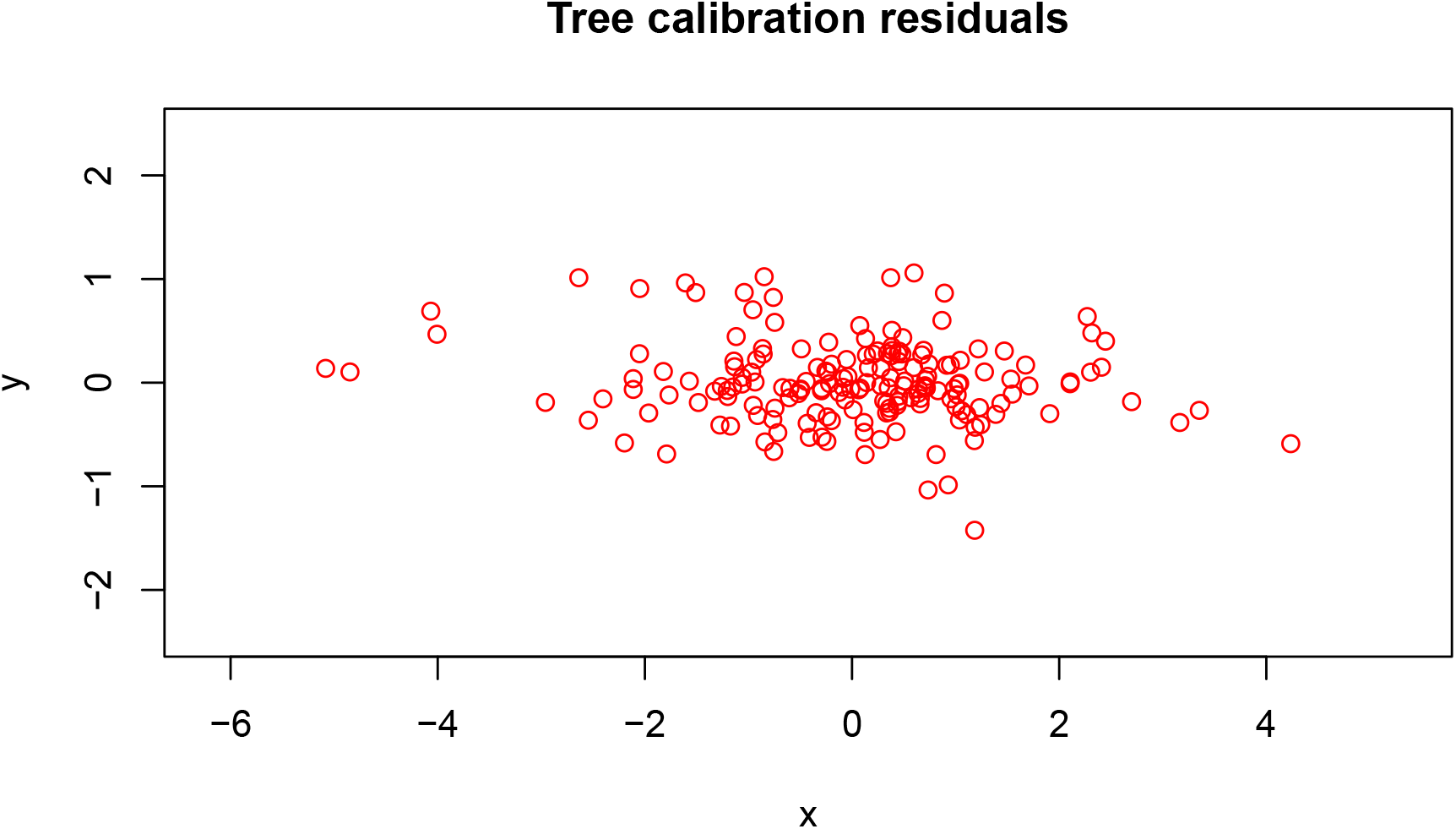
Error-model residuals from GPS calibration data where the tag was placed on the side of a tree. As the tree blocks out half of the sky, the spread in satellites is narrower along the axis perpendicular to the tree, which results in larger errors along that axis. Other situations where GPS location errors are elongated include when obtaining location fixes alongside cliffs or buildings.

and the standardized or “studentized” residuals are given by

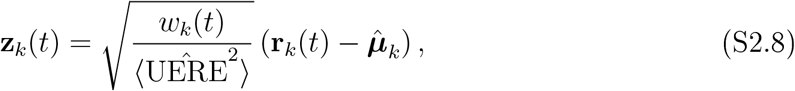

which can be checked for substantial deviance from normality or autocorrelation, as in Fig. S4.1.

#### S2.1.1 Single-UERE model selection

There are several situations where model selection is useful when calibrating data. First, while DOP values are usually informative, we do not always know the best proxy for the location-error variance (c.f., Sec. S3). Second, we may be unsure as to whether different devices share the same UERE parameters. Third, we may be unsure as to whether or not different location classes share the same UERE parameters. Therefore, it is useful to have AIC_C_ values to facilitate model selection. AIC values estimate the Kullback-Leibler divergence between the fitted model and true model (Burnham and Anderson, 2002). In situations where sample sizes are small and debiased parameter estimates differ substantially from maximum-likelihood (ML) parameter estimates, then AIC_C_ values can differ substantially from regular AIC values. Standard AIC_C_ formulas are specific to maximum-likelihood estimation with a simple error model, whereas here we do better than ML estimation with more complex error models.

Following analogous calculations in (Calabrese et al., 2018; Fleming et al., 2019), unbiased AIC_c_ values must satisfy the cross-validated log-likelihood, or double expectation 〈〈···〉〉 (over both the data and estimates calculated from independent data) of the log-likelihood function, regardless of whether or not the estimates are derived from maximizing said likelihood function.

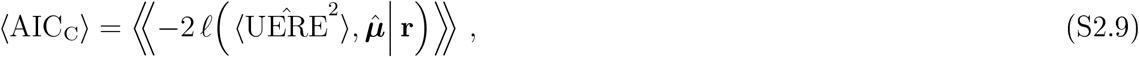

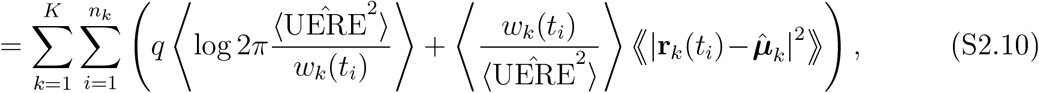

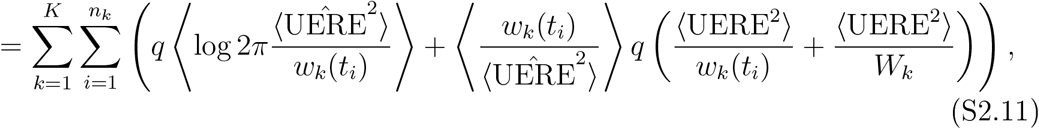

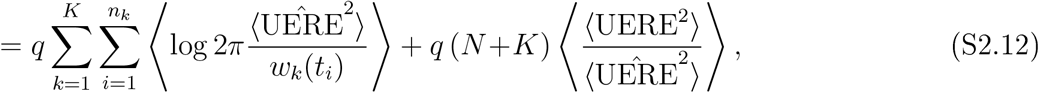

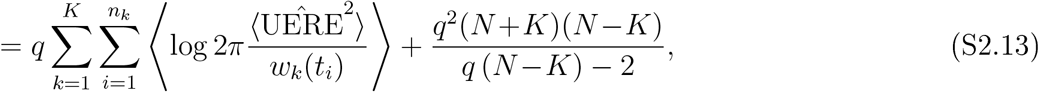

and so by the Lehmann-Scheffé theorem (Lehmann and Scheffé, 1950, 1955), the MVU AIC_C_ values must be given by

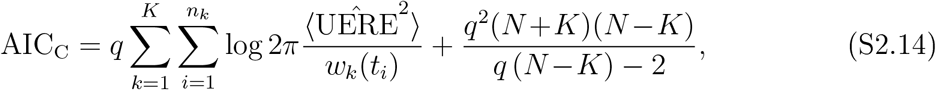

which one can check for asymptotic consistency with the standard AIC formula.

#### S2.1.2 Single-UERE goodness of fit

AIC_C_ differences allow us to rank our models’ expected predictive capability, but these differences scale with sample size and so they do not indicate how well the models perform in an absolute sense. If our variance model was fixed and our comparison was among trend models, ***μ***, then we might consider the reduced *χ*^2^ statistic to index goodness of fit. However, our trend model is fixed and we are comparing variance models, and so we derive a reduced *Z*^2^ statistic that is analogous to the familiar reduced *χ*^2^ statistic.

We start by considering the “internally” studentized residuals, as goodness of fit can be measured by their closeness to unity.

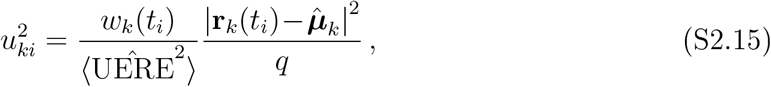

These square residuals are not exactly F-distributed because the numerators and denominators are not independent. Next we apply a trick due to Thompson (1935), decomposing all terms into independent variates.

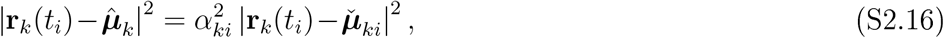

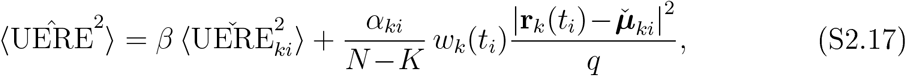

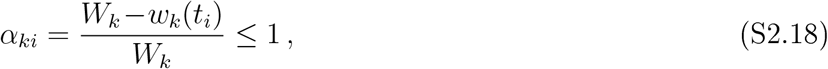

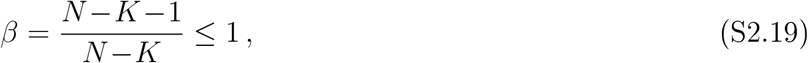

where all 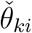-estimates are calculated leaving out the *ki*^th^ observation, and then the “externally” studentized residuals

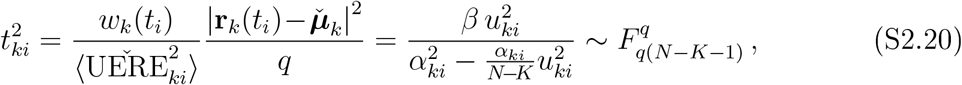

are exactly *F*-distributed. Importantly, there is a one-to-one correspondence between the internally studentized *u*^2^ and externally studentized *t*^2^ statistics, and so it does not matter which statistic we base our index on. Furthermore, relation (S2.49) is particularly useful because direct calculation of all 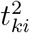 statistics via the leave-one-out 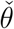-estimators has a computational cost of 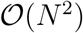, whereas calculation here via *u_ki_* has a computational cost of only 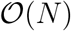.

Again, goodness of fit can be measured by *t*^2^ being close to unity. Both small values of *t*^2^, where the model variance is too large, and large values of *t*^2^, where model variance is too small, are equally bad and so we proceed to Fisher’s *Z*-distribution

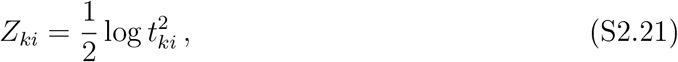

which places extreme values of *t*^2^ on the same relative scale, with a variance model that is a factor too small being just as bad as a variance model that is a factor too large. Goodness of fit is now determined by the proximity of *Z* to zero, and lack of performance can be inferred from the magnitude of *Z*^2^, which for all *ik* has an expectation value being only a function of the degrees of freedom. Our reduced *Z*^2^ statistic is given by

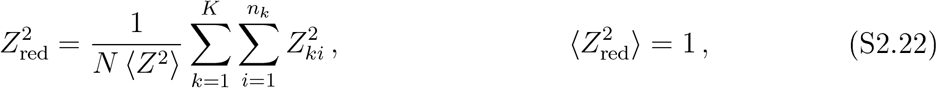

where, 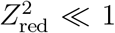 indicates an over-performing model, 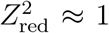 indicates a well-performing model, and 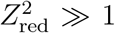 indicates an under-performing model, all relative to a true Gaussian model. If the true UERE distribution is not Gaussian, the reduced *Z*^2^ statistic can still be used to compare performance across error models, only the value of 1 no longer separates overperformance from under-performance. In this context, over-performance indicates that the square errors are exceptionally proportional to the model variances, while under-performance indicates a weaker relationship between the two quantities. For example, bivariate Laplace errors with correctly modeled variance will produce 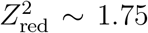 when using the Gaussian formula for 〈*Z*^2^〉 Even more extreme, bivariate Cauchy errors—which do not admit a finite variance—limit to 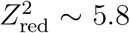 with large *N*. All else being equal, a more heavy-tailed distribution will give rise to a larger value of 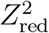 under the normal assumption.

Finally, for any 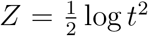, where 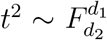 as above, the desired second moment 〈*Z*^2^〉 is straightforwardly calculated to be

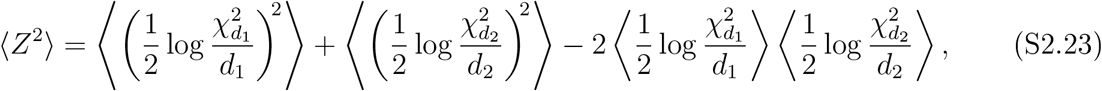

in terms of the normalized log-*χ*^2^ moments

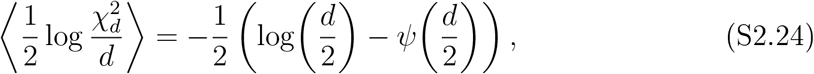

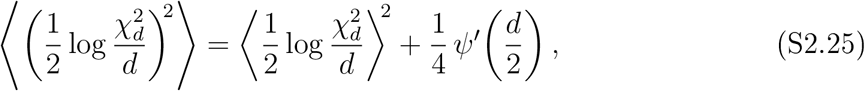

where 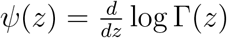 is the digamma function and Γ(*z*) = (*z* – 1)! is the gamma function.

#### S2.1.3 Mean-zero processes

The simplifications to the previous formulas for mean-zero process calibration, such as velocity, are

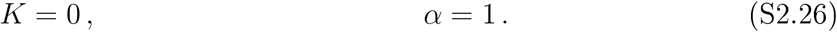

### S2.2 Multi-UERE estimation

There are several situations where multiple RMS UEREs need to be estimated. Many GPS modules record different quality locations, including 2D, 3D, “quick fixes” resolved and unresolved, but no HDOP information or their HDOP values derive from different methods and are not on the same scale. In some cases, unresolved location quality can only be inferred from missing attributes like speed and altitude, or from the fix time reaching its maximum value, which requires one RMS UERE value for fix times below the threshold value and a larger RMS UERE value for timed-out fixes.

When there are *C* > 1 location classes for which we need to estimate *C* RMS UERE parameters, we cannot derive MVU estimators, but we can derive residual maximum likelihood (REML) estimators that reduce to MVU when *C* = 1. Let *c* index the *C* location classes, and define the indicator function

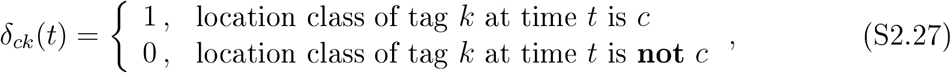

then the REML log-likelihood function is

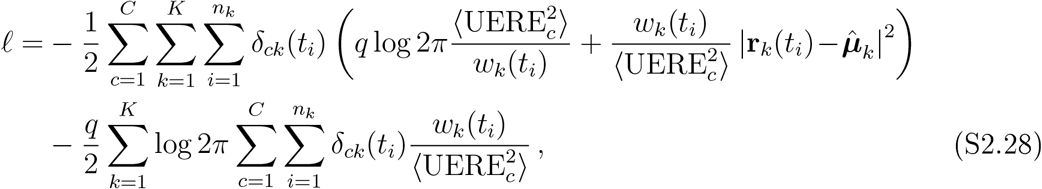

where the last term on the right hand side is the REML correction to the regular log-likelihood function.

The mean-square UERE and mean location estimating equations are given by

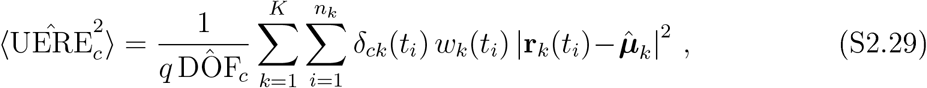

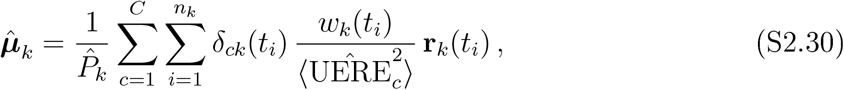

given the DOF and precision-weight estimates

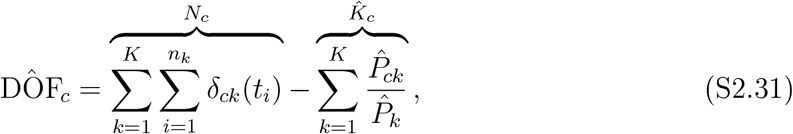

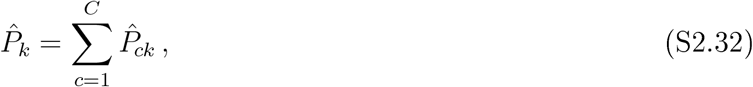

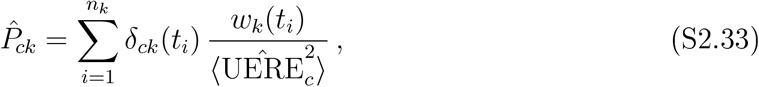

where the second term on the right-hand-side of (S2.31) is the REML correction to the MLE. We solve the estimating equations iteratively, by evaluating the above relations in reverse order, starting with an initial guess for the RMS UEREs. In the case of one location class (*C* = 1), this algorithm converges in one iteration. For *C* > 1, convergence is at worst quadratic and machine precision is typically achieved in a few iterations.

Effectively, the *K* lost degrees of freedom are distributed across the *C* RMS UERE estimates (*K_c_*), according to the precision weights *P_ck_*, as it is straightforward to show that

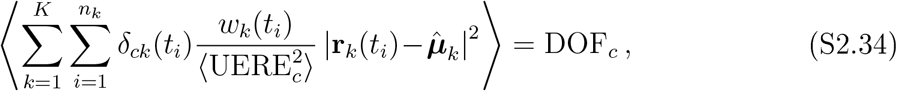

at the true RMS UERE values, and the total number of degrees of freedom are indeed *N* – *K*.

The residuals are given by

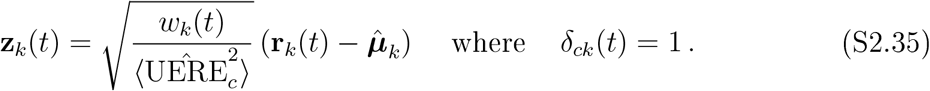

From the Hessian of the REML log-likelihood function, the sampling distributions of interest are asvmptotically given by

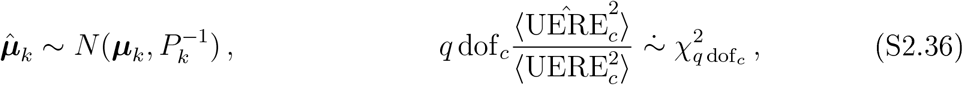

in terms of the degrees of freedom

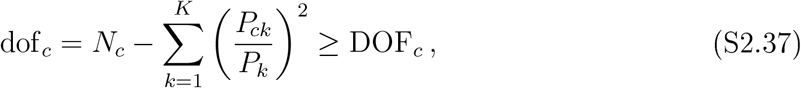

which is larger than the degrees of freedom in the estimating equations that decompose the variance of the residuals (S2.31), though still smaller than the biased maximum likelihood value. Both dof_c_ and DOF_c_ reduce to *N* – *K* when *C* =1.

#### S2.2.1 Multi-UERE model selection

Generalizing the results of Sec. S2.1.1, we again evaluate the cross-validated log-likelihood. However, instead of an exact calculation, we must make asymptotic approximations ignoring the correlation between the RMS UERE estimates and mean location estimates. These approximations result in an improved, though not fully unbiased, estimator of the Kullback-Leibler divergence (AIC).

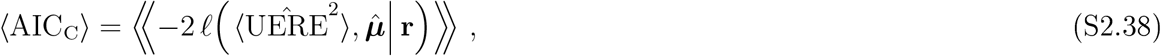

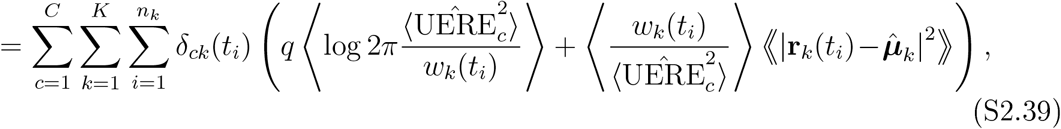

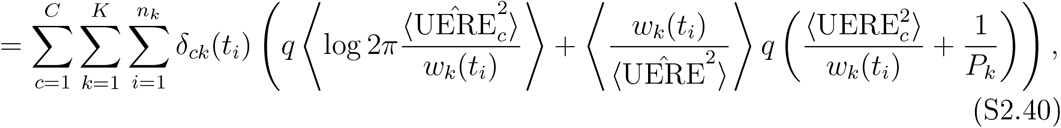

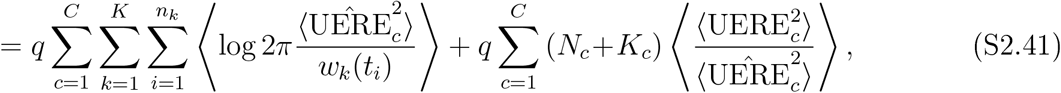

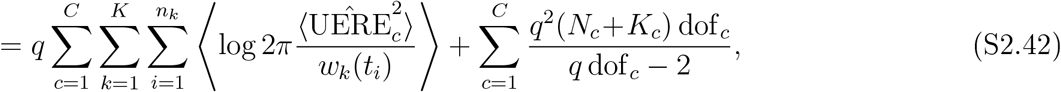

and so our second-order AIC_C_ values are given by (S2.43)

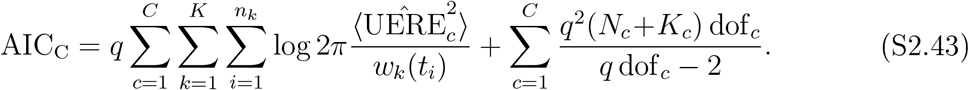

#### S2.2.2 Multi-UERE goodness of fit

Again we start with the internally studentized residuals

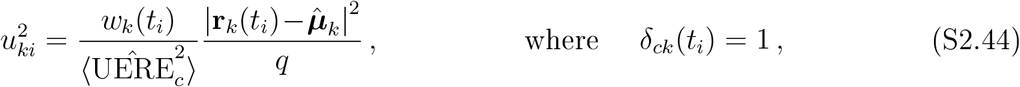

and apply the method of Thompson (1935)

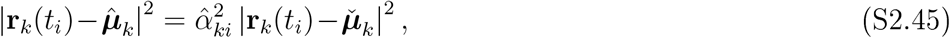

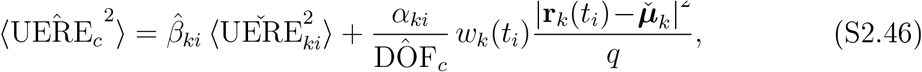

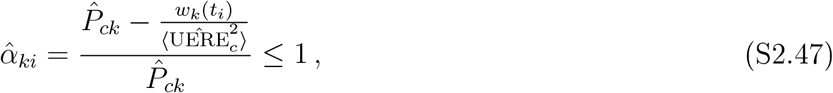

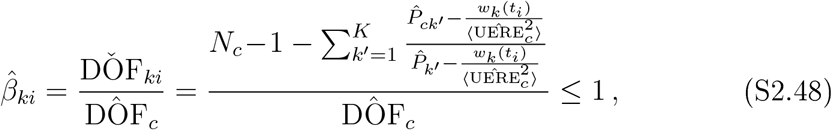

to transform into the externally studentized residuals

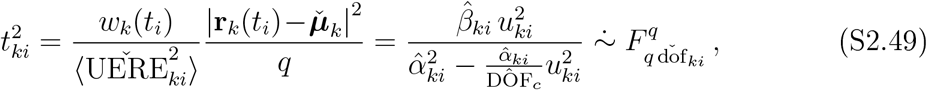

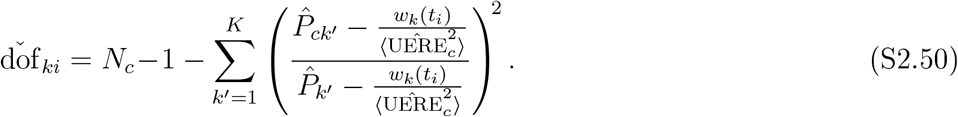

The relationship between the internally and externally studentized residuals is no longer as clear, but this remains a computationally efficient way of calculating the externally studentized residuals, for which we have a better approximation to the statistic’s distribution.

Fisher’s *Z*-statistic is then given by

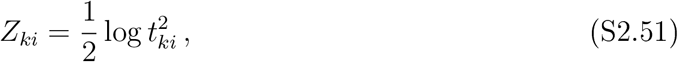

and our reduced *Z*^2^ statistic is given by

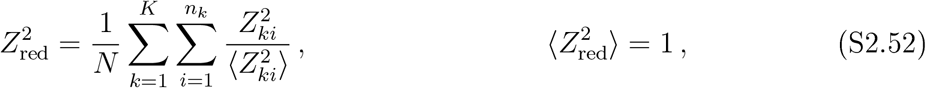

with the necessary formulas for 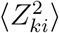 calculated in relation (S2.23).

#### S2.2.3 Mean-zero processes

The simplifications to the previous formulas for mean-zero process calibration, such as velocity, are

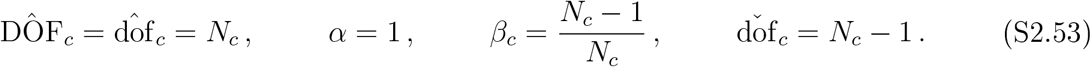

## S3 Precision models in practice

Here we detail precision models beyond null model (1.1), for GPS devices that do not supply informative DOP values or exhibit complications. In App. S4, these models are applied to a wide range of devices, demonstrating their coverage and utility.

### S3.1 DOP proxies

#### Co-opting DOP values

In lieu of HDOP values, some GPS devices only record “position DOP” (PDOP), “geometric DOP” (GDOP), or ambiguous DOP values. PDOP values combine HDOP and VDOP; GDOP values combine PDOP and “time DOP” (TDOP) values. As HDOP and VDOP values are both a function of satellite number and spread, they are correlated. Therefore, it is reasonable to co-opt or appropriate related DOP values in the absence of proper HDOP and VDOP values. Model selection can then determine whether alternative or ambiguous DOP values are more predictive than nothing (homoskedastic location errors).

By default, when importing data in to `ctmm` via the `as.telemetry()` function, and assuming the data are not of a special variety (Argos Doppler-shift or e-obs), then the `as.telemetry()` will first look for an HDOP value. If HDOP values are not found, then `as.telemetry()` will look for ambiguous DOP values, followed by PDOP and then GDOP values. If no DOP values are found, then `as.telemetry()` will look for the reported number of satellites, and apply model (S3.1) to approximate DOP values.

#### Number of Satellites

Some GPS data lack DOP values but do record the number of satellites used when triangulating location estimates. Therefore, as proxy for DOP, a positive, monotonically decreasing function of the number of satellites, *N*_sat_, can be used. The simplest error variance relation would decay with the density of the GPS satellite constellation, which is proportional to 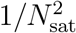. Furthermore, trilateralization is not possible with *N*_sat_ ≤ 2. Therefore, we can improve our simplest model to

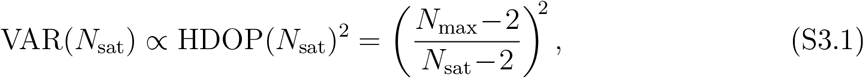

where *N*_max_ = 12, which fixes the minimum HDOP value to 1 for convenience. Fig. S3.1 depicts the relationship between HDOP and *N*_sat_, along with our best-fit model. GPS data was taken from 35 blue wildebeests (*Gorgon taurinus albojubatus*) and HDOP values were aggregated into RMS HDOP values and their variance, per value of *N*_sat_, as non-linear leastsquares regressions would not converge when using the full dataset (likely due to truncation error in the DOP values). To approximate an exact regression on the full dataset, we applied a weighted non-linear least squares regression to the RMS HDOP values under a log link. We considered all combinations of models with a power of 2 or unknown and a singularity at *N*_sat_ values of 0, 2, or unknown. In terms of both AIC and BIC, our simple model (S3.1) outperformed all others, save for the most complex model with both unknown power and unknown singularity. However, the more complex model could not maintain performance under different modeling choices—including HDOP summary statistic, regression weights, and link function—and its parameter estimates were not meaningful.

**Figure S3.1:**
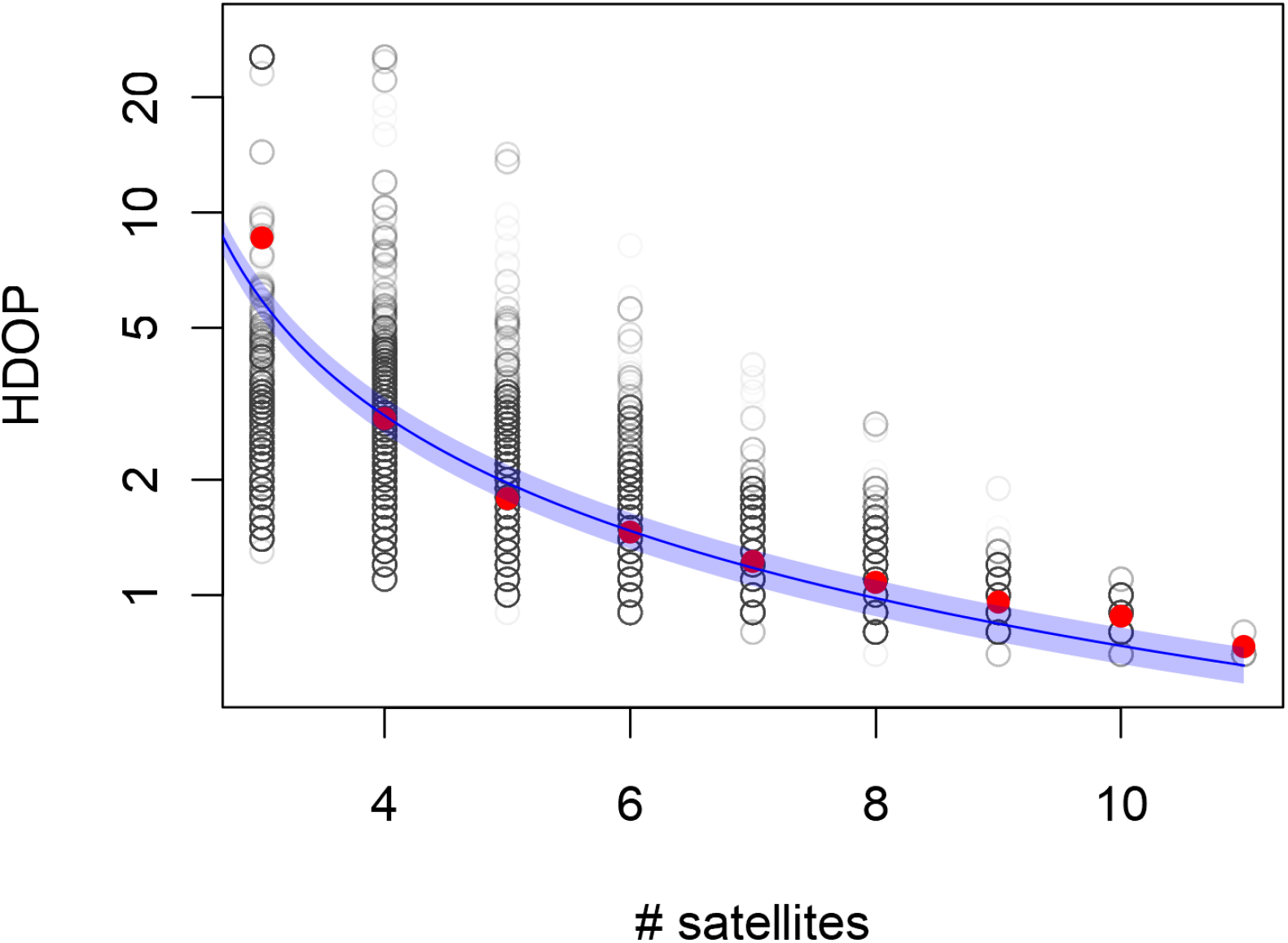
Horizontal dilution of precision (HDOP) versus number of satellites from 35 GPS collared wildebeests, with the RMS HDOP in red and approximate HDOP model (S3.1) in blue.

### S3.2 Precision modifiers

#### Fix Type

Some GPS devices specify a categorical “fix type” that distinguishes location estimates obtained via different algorithms or with different characteristics. The most common fix-type levels are 2D versus 3D, where the 2D location estimates are calculated with only 3 satellites and lack corresponding altitude estimates. In principle, HDOP values are supposed to account for the diminished accuracy of 2D location estimates, however, some GPS devices require distinct 2D/3D RMS UERE values (App. S4). Furthermore, some GPS devices require distinct 2D/3D parameters, yet do not provide an explicit “GPS fix-type” column. In this case, a fix-type column can be calculated from the logical test *N*_sat_ > 3.

By default, `ctmm`’s `as.telemetry()` function will assume that any GPS fix-type column is meaningful, and create a corresponding location class column in the output `telemetry` data object. Model selection can then determine whether or not extra location classes are supported by the calibration data, `as.telemetry()` also checks if some location estimates lack corresponding speed, altitude, and/or DOP values, in which case location classes will be created to distinguish the level of missingness, as this often corresponds to different location estimation algorithms.

#### Time to fix

GPS devices take some amount of time to acquire satellite signals and corresponding satellite orbital information before estimating a location. Location estimates can either be calculated with the full orbital information from each satellite in communication, or approximate orbital information from fewer satellites. In some GPS devices, this change in location estimate quality occurs at a maximum recorded “time to fix” value, when the device terminates its attempt to obtain full orbital information and proceeds to an approximate calculation. If this approximation is not accounted for in the device’s HDOP values, then separate “in time” and “timed out” RMS UERE values may be required (App. S4). Furthermore, as the level of approximation can vary considerably in “timed out” fixes, their UERE distribution may be heavy tailed.

In `ctmm`’s `as.telemetry()` function, the timeout argument can be used to create an additional location class for TTF values greater than or equal to the timeout value.

## S4 Error-model selection examples

### S4.1 Advanced Telemetry Systems

#### S4.1.1 ATS G2110e

We analyzed calibration data on nineteen Advanced Telemetry Systems (ATS) G2110e GPS/Iridium collars with two deployments each, where data were collected in 10 and 60 minute intervals for an average of 2.5 days. Before calibration, we removed all gross outliers, which constituted 0.3% of the total records. In our first round of analysis, we tested the veracity of the recorded HDOP values against our number-of-satellites model (S3.1) and homoskedastic location errors. We found the provided HDOP values to be more informative than nothing (homoskedastic errors), but inferior to our simple number-of-satellites model, which should not be the case. Performance of the initially best model, 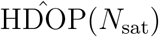, was poor 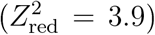, with a large variation among devices. Upon further investigation into other data provided by the GPS devices, we notice a smooth, positive trend between the calibration residuals and time to fix. Therefore we also included powers of (TTF/TTF_min_) in our HDOP model, with the selected model being a product of our number-of-satellites model and square TTF (Table S4.1), and where the expected RMS location error of this device is 2.4-2.5 meters under ideal conditions (*N*_sat_ = 12 TTF = TTF_min_), which is small enough to indicate a higher quality dual-frequency GPS receiver or on-board Kálmán filtering.

To assess variability among GPS devices, we compared our best joint model to the same class of model, but with calibration individualized by GPS device and by deployment. In either case, the increase in performance was not substantial.

**Table S4.1:** AIC_C_ differences and reduced *Z*^2^ values for error models fit to calibration data from nineteen ATS G2110e GPS collars with two deployments each.

| Model | $\Delta\text{AIC}_C$ | $Z_{\text{red}}^2$ |
| --- | --- | --- |
| $\hat{\text{HDOP}}(N_{\text{sat}}) \times \text{TTF}^2$ | 0.0 | 2.9 |
| $\hat{\text{HDOP}}(N_{\text{sat}}) \times \text{TTF}^3$ | 816.4 | 3.3 |
| $\text{HDOP} \times \text{TTF}^2$ | 1,432.7 | 3.2 |
| $\hat{\text{HDOP}}(N_{\text{sat}}) \times \text{TTF}^1$ | 1,716.0 | 3.0 |
| $\text{HDOP} \times \text{TTF}^3$ | 2,403.2 | 3.7 |
| $\text{HDOP} \times \text{TTF}^1$ | 2,945.7 | 3.3 |
| $\text{TTF}^2$ | 4,403.0 | 3.6 |
| $\text{TTF}^3$ | 5,111.6 | 4.0 |
| $\text{TTF}^1$ | 6,942.7 | 4.0 |
| $\hat{\text{HDOP}}(N_{\text{sat}})$ | 7,285.0 | 3.9 |
| HDOP | 8,060.7 | 4.1 |
| homoskedastic | 1,4863.6 | 5.3 |

#### S4.1.2 ATS G5-2D

We analyzed calibration data from four ATS G5-2D GPS/Iridium collars, with data collected in 10-minute intervals for 10-12 days. We pitted the reported HDOP values against our numbcr-of-satcllitcs model (S3.1) and a model of homoskedastic errors. Performance of all three error models was very similar (Table S4.2), though HDOP values only ranged from 0.6 to 2.1, which implies a consistently good signal during calibration. RMS error under ideal conditions (HDOP=0.6) was estimated to be 3.5-3.6 meters. Variation among devices was not substantial.

**Table S4.2:** AIC_C_ differences and reduced *Z*^2^ values for error models fit to calibration data from four ATS G5-2D GPS collars.

| Model | $\Delta\text{AIC}_C$ | $Z^2_{\text{red}}$ |
| --- | --- | --- |
| homoskedastic | 0.0 | 1.5 |
| HDOP | 34.7 | 1.5 |
| $\hat{\text{HDOP}}(N_{\text{sat}})$ | 39.6 | 1.5 |

In cases such as this, where the calibration data had consistently good reception yet the tracking data might not, we suggest performing error-model selection (but not parameter estimation) again with the tracking data. It could very well be the case that the HDOP or 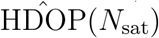 error models prove superior when faced with poor satellite reception.

#### S4.1.3 ATS G10 UltraLITE

We analyzed calibration data from 15 ATS G10 UltraLITE GPS tags deployed in 4 environments— deciduous forest, evergreen forest, forested wetland, and clearing—with data collected in 10-minute intervals for 2-15 days. For both horizontal and vertical errors we tested the provided HDOP, PDOP, and VDOP values against our numbcr-of-satcllitcs model (S3.1) and a model of homoskedastic errors (Table S4.3). Horizontal errors were comparably modeled by either HDOP value or our 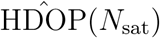 model, with the RMS error estimated to be 21.0–21.3 meters under ideal conditions (*N*_sat_ = 12). However, VDOP values were slightly worse than all other predictors for vertical errors. There was some variability in horizontal calibration parameters, with 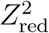 dropping from 2.9 to 2.7 upon individual calibration. Differences in performance among habitats was not as significant.

**Table S4.3:** AIC_C_ differences and reduced *Z*^2^ values for error models fit to calibration data from 15 ATS GIO UltraLITE GPS tags.

| Horizontal |  |  | Vertical |  |  |
| --- | --- | --- | --- | --- | --- |
| Model | $\Delta\text{AIC}_C$ | $Z^2_{\text{red}}$ | Model | $\Delta\text{AIC}_C$ | $Z^2_{\text{red}}$ |
| $\hat{\text{HDOP}}(N_{\text{sat}})$ | 0.0 | 2.9 | homoskedastic | 0.0 | 2.4 |
| HDOP | 10.5 | 2.9 | HDOP | 27.6 | 2.4 |
| PDOP | 5,993.9 | 3.4 | PDOP | 176.6 | 2.4 |
| homoskedastic | 6,174.9 | 3.4 | $\hat{\text{HDOP}}(N_{\text{sat}})$ | 343.6 | 2.4 |
| VDOP | 16,133.3 | 4.3 | VDOP | 2,331.3 | 2.5 |

### S4.2 Cellular Tracking Technologies

#### S4.2.1 CTT 1090

We analyzed test data from 3 Cellular Tracking Technologies (CTT) 1090 GPS-GSM tags, with data collected every 15 minutes for up to 36 hours. For both horizontal and vertical errors, we compared the performance of reported HDOP and VDOP values against a model of homoskedastic location errors after removing 3 gross outliers (Table S4.4). RMS horizontal location error was estimated to be 8.7–10.3 meters under ideal conditions (HDOP=1). Individualized calibration had some impact on performance, decreasing 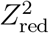 from 4.6 to 4.3.

**Table S4.4:** AIC_C_ differences and reduced *Z*^2^ values for error models fit to calibration data from three CTT 1090 GPS tags.

| Horizontal |  |  | Vertical |  |  |
| --- | --- | --- | --- | --- | --- |
| Model | $\Delta\text{AIC}_C$ | $Z_{\text{red}}^2$ | Model | $\Delta\text{AIC}_C$ | $Z_{\text{red}}^2$ |
| HDOP | 0.0 | 4.6 | HDOP | 0.0 | 2.1 |
| VDOP | 175.2 | 6.9 | homoskedastic | 69.6 | 1.6 |
| homoskedastic | 191.8 | 6.4 | VDOP | 70.0 | 1.8 |

#### S4.2.2 CTT BT3

We analyzed calibration data from 8 Cellular Tracking Technologies (CTT) BT3 GPS tags, with data collected in 10 and 15 minute intervals for up to one month. For both horizontal and vertical errors, we compared the performance of the reported HDOP and VDOP values to our 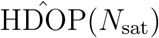 model (S3.1) and a model of homoskedastic location errors (Table S4.5). There was some variability in horizontal calibration parameters, with 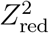 dropping from 4.3 to 4.0 upon individual calibration.

**Table S4.5:** AIC_C_ differences and reduced *Z*^2^ values for error models fit to calibration data from five CTT BT3 GPS tags.

| Horizontal |  |  | Vertical |  |  |
| --- | --- | --- | --- | --- | --- |
| Model | $\Delta\text{AIC}_C$ | $Z_{\text{red}}^2$ | Model | $\Delta\text{AIC}_C$ | $Z_{\text{red}}^2$ |
| HDOP | 0.0 | 4.3 | $\hat{\text{HDOP}}(N_{\text{sat}})$ | 0.0 | 2.3 |
| $\hat{\text{HDOP}}(N_{\text{sat}})$ | 1,528.0 | 4.7 | HDOP | 827.4 | 2.4 |
| VDOP | 3,685.0 | 5.1 | VDOP | 913.2 | 2.3 |
| homoskedastic | 4,300.5 | 5.0 | homoskedastic | 1,744.6 | 2.3 |

#### S4.2.3 CTT ES400

We analyzed calibration data from 15 Cellular Tracking Technologies (CTT) ES400 (4 ^th^ Gen) GPS-GSM solar powered telemetry units, with data collected in 10- and 15-minute intervals for two weeks. For both horizontal and vertical errors, we compared the performance of the reported HDOP and VDOP values to our 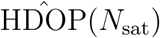 model (S3.1) and a model of homoskedastic location errors (Table S4.6). Location errors in both dimensions were best modeled by our 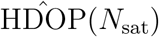 model, though HDOP was a close second. Curiously, the VDOP-based model was outperformed by both 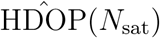 and HDOP for vertical errors. RMS horizontal error was estimated to be 3.9–4.0 meters under ideal conditions (*N*_sat_=12). There was slight variability among devices, with 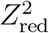 dropping from 2.4 to 2.3 upon individual horizontal calibration.

**Table S4.6:** AIC_C_ differences and reduced *Z*^2^ values for error models fit to calibration data from five CTT ES400 GPS collars.

| Horizontal |  |  | Vertical |  |  |
| --- | --- | --- | --- | --- | --- |
| Model | $\Delta\text{AIC}_C$ | $Z_{\text{red}}^2$ | Model | $\Delta\text{AIC}_C$ | $Z_{\text{red}}^2$ |
| $\widehat{\text{HDOP}}(N_{\text{sat}})$ | 0.0 | 2.4 | $\widehat{\text{HDOP}}(N_{\text{sat}})$ | 0.0 | 1.3 |
| HDOP | 1,527.8 | 2.4 | HDOP | 732.9 | 1.3 |
| homoskedastic | 10,895.3 | 2.9 | VDOP | 4,059.6 | 1.4 |
| VDOP | 16,075.8 | 3.2 | homoskedastic | 4,138.7 | 1.5 |

### S4.3 e-obs

e-obs devices generally do not provide DOP values, but a “horizontal accuracy estimate”, which, according to e-obs documentation is an *in*accuracy estimate measured in meters. From personal communications with e-obs, the statistical interpretation of this information— in terms of a specific number of standard deviations or quantile—is unknown. Based on the following results, `ctmm`’s `as.telemetry` function imports e-obs horizontal accuracy estimates as HDOP values.

#### S4.3.1 e-obs generation-1 GPS collars (1059-1078)

We analyzed calibration data collected from 17 first-generation e-obs GPS collars encircled around a tree and sampled at 10 minute intervals over the course of a day. To confirm the veracity of the e-obs error estimates we performed a null hypothesis test, with a null model of homoskedastic location errors of unknown variance, and an alternative model with error standard deviations proportional to the e-obs error estimate by treating the e-obs error estimates as dimensionful HDOP values. We found the e-obs error estimates to be highly informative (Table S4.7), and from these data the RMS “UERE” value was estimated to be 1.67 (1.63–1.70). RMS horizontal error was estimated to be 5.8–6.1 meters under ideal conditions. There was slight variability in the devices, with 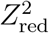 decreasing from 2.1 to 2.0 when using individualized horizontal calibration, and where collars with a higher percentage of missing data yielded smaller RMS UERE estimates.

**Table S4.7:** AIC_C_ differences and reduced *Z*^2^ values for error models fit to calibration data from 17 e-obs GPS tags.

| Horizontal |  |  | Vertical |  |  |
| --- | --- | --- | --- | --- | --- |
| Model | $\Delta\text{AIC}_C$ | $Z_{\text{red}}^2$ | Model | $\Delta\text{AIC}_C$ | $Z_{\text{red}}^2$ |
| horizontal-accuracy | 0.0 | 2.1 | horizontal-accuracy | 0.0 | 1.6 |
| homoskedastic | 2,442.9 | 3.1 | homoskedastic | 160.6 | 1.6 |

#### S4.3.2 e-obs generation-3 GPS tags (5081-6375)

We analyzed calibration data from 15 third-generation e-obs solar tags placed at a fixed location and sampled for 10-20 days with 5-20 minute sampling (depending on battery status) and 1-second sampling in direct sunlight for a minimum of 10 minutes. With these data, and using the same error model, the e-obs RMS UERE value was estimated to be 1.673 (1.672–1.674), consistent with our first-generation GPS collars and highly informative (Table S4.8). Variation among devices was not substantial.

**Table S4.8:** AIC_C_ differences and reduced *Z*^2^ values for error models fit to calibration data from 15 e-obs GPS tags. ^†^Due to autocorrelation, the AIC_C_ differences are exaggerated by a factor on the order of a hundred.

| Horizontal |  |  | Vertical |  |  |
| --- | --- | --- | --- | --- | --- |
| Model | $\Delta\text{AIC}_C^\dagger$ | $Z_{\text{red}}^2$ | Model | $\Delta\text{AIC}_C^\dagger$ | $Z_{\text{red}}^2$ |
| horizontal-accuracy | 0.0 | 1.3 | horizontal-accuracy | 0.0 | 1.1 |
| homoskedastic | 6,167,049 | 3.2 | homoskedastic | 2,430,763 | 1.8 |

Due to the presence of substantial autocorrelation in these data, which we did not account for in estimating the RMS UERE, both the point estimate and standard error are downwardly biased. We roughly approximate the influence of autocorrelation by inspection of the correlogram (Fig. S4.1), where we note that most of the autocorrelation is gone within 2 minutes of time lag. Following the analysis of Fleming et al. (2019), 10 days of sampling at 2-minute intervals would yield an effective sample size on the order of 7200 (10 days ? 2 minutes), and a resulting bias of 1/7200, which is not substantial. The standard errors, however, are on the order of 11 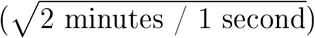 times too small. Therefore, a corrected RMS UERE estimate would be closer to 1.67 (1.66-1.68), which is still highly consistent with our first calibration results. Similarly, the RMS horizontal error would then be 1.70–1.72 meters under ideal conditions, which includes the data being collected at 1 hertz.

Inspection of the 1-second solar-tag error residuals and error estimates versus time since the unit was “cold” (not collecting 1-second fixes) revealed RMS location errors that shrink over time, from 30 meters initially to 3 meters after two minutes, which is evidence of on-board error-filtering of the location estimates. Likely related to the e-obs error filter, we found two further issues in the 1-second solar tag data not present in the 10-minute collar data. First, the e-obs error estimates were more substantially autocorrelated for a more prolonged period of time (Fig. S4.1). Second, the first few seconds of cold fixes had substantially underestimated location errors (~ 25%).

#### S4.3.3 e-obs generation-3 24-gram GPS tags (6577-6752)

We analyzed calibration data from 6 third-generation e-obs 24-gram GPS tags, with location fixes collected every 2 minutes for 3 days. Three of the tags were placed under forest canopy and three were placed on a roof. We compared the e-obs “horizontal accuracy estimate” to a model of homoskedastic errors. The e-obs horizontal accuracy estimate proved to be informative (Table S4.9), but, in contrast to older e-obs models, the RMS UERE was estimated to be 0.94 (0.93–0.95). RMS horizontal error was estimated to be 2.6–2.7 meters under ideal conditions, which suggests a dual-frequency GPS receiver or on-board Kálmán filter. There was substantial heterogeneity among devices, with AIC_C_ decreasing by 4,903.9 and Z2d dropping from 2.5 to 1.9 when using individiualized horizontal calibration. This heterogeneity could not be explained by habitat (ΔAIC_C_ = 4,846.9).

**Figure S4.1:**
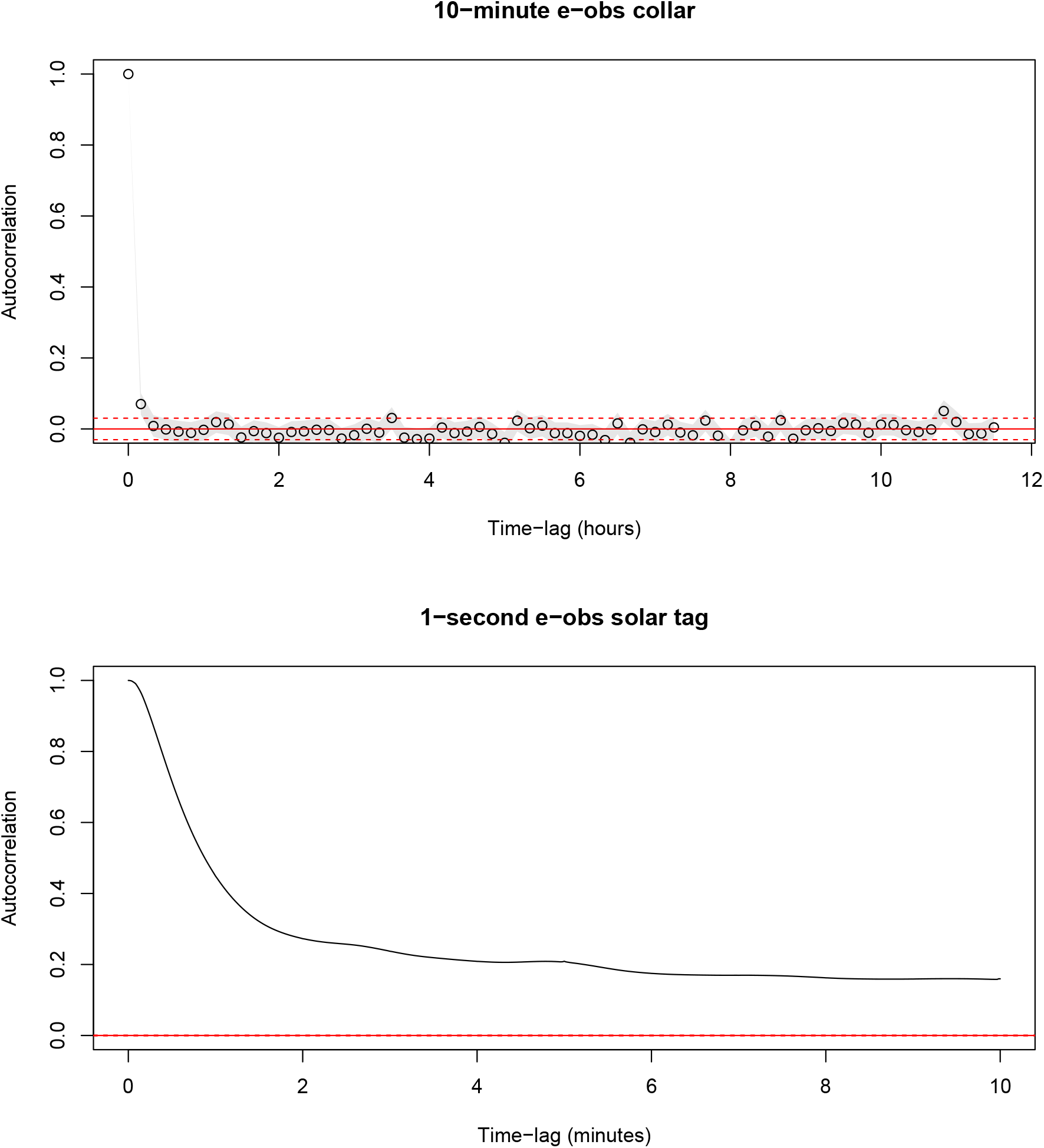
Joint correlograms of standardized calibration residuals from (**A**) 17 e-obs collars sampled at 10-minute intervals and (**B**) 15 e-obs solar tags sampled at 1-second intervals. Autocorrelation is not very substantial in the 10-minute calibration residuals (< 10%), though it is significant at the first lag. In contrast, the 1-second e-obs data are substantially autocorrelated even after 10 minutes of time lag.

**Table S4.9:** AIC_C_ differences and reduced *Z*^2^ values for error models fit to calibration data from 6 third-generation e-obs GPS tags.

| Horizontal |  |  | Vertical |  |  |
| --- | --- | --- | --- | --- | --- |
| Model | $\Delta AIC_C$ | $Z_{red}^2$ | Model | $\Delta AIC_C$ | $Z_{red}^2$ |
| horizontal-accuracy | 0.0 | 2.5 | homoskedastic | 0.0 | 1.8 |
| homoskedastic | 14,234.7 | 3.8 | horizontal-accuracy | 614.9 | 1.8 |

#### S4.3.4 e-obs generation-3 42-gram GPS tag (7095-7106)

We analyzed calibration data from 5 third-generation e-obs 42-gram GPS tags, with location fixes collected every 2 minutes for 3 days. Three of the tags were placed under forest canopy and two were placed on a roof. We compared the e-obs “horizontal accuracy estimate” to recorded HDOP values, our number-of-satellites error model (*N*_sat_; S3.1), and a model of homoskedastic location errors (Table S4.10). The e-obs “horizontal accuracy estimate” proved to be informative, with the RMS UERE estimated to be 0.99 (0.98–1.00). The RMS UERE value of ~1 is close to some other third-generation e-obs calibration values and consistent with the interpretation of the e-obs “horizontal accuracy estimate” being an estimate of the root-mean-square location error. RMS horizontal error was estimated to be 2.5–2.6 meters under ideal conditions, which suggests a dual-frequency GPS receiver or on-board Kálmán filter. There was slight heterogeneity among devices, with 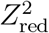 dropping from 1.8 to 1.7 when using individualized calibration.

**Table S4.10:** AIC_C_ differences and reduced *Z*^2^ values for error models fit to calibration data from 5 third-generation e-obs GPS tags.

| Horizontal |  |  | Vertical |  |  |
| --- | --- | --- | --- | --- | --- |
| Model | $\Delta AIC_C$ | $Z_{red}^2$ | Model | $\Delta AIC_C$ | $Z_{red}^2$ |
| horizontal-accuracy | 0.0 | 1.8 | horizontal-accuracy | 0.0 | 1.6 |
| $\widehat{HDOP}(N_{sat})$ | 12,123.4 | 3.3 | HDOP | 635.3 | 1.8 |
| homoskedastic | 12,166.1 | 3.2 | $\widehat{HDOP}(N_{sat})$ | 1,119.8 | 1.8 |
| HDOP | 12,630.7 | 3.5 | homoskedastic | 1,520.5 | 1.8 |

#### S4.3.5 e-obs generation-3 GPS tag (7401-7405)

We analyzed calibration data from 3 third-generation e-obs GPS tags, with location fixes collected every half hour or hour for 3-6 days in four different environments—on an open deck, beneath a tree, indoors, and beneath a house. For both horizontal and vertical errors, we compared the e-obs “horizontal accuracy estimate” to ambiguous DOP values, our number-of-satellites error model (*N*_sat_; S3.1), and a model of homoskedastic location errors (Table S4.11). Again, the e-obs “horizontal accuracy estimate” proved to be the most informative predictor of horizontal errors, but with the RMS UERE estimated to be 0.86 (0.84-0.88), which is smaller than that of previous e-obs devices. There was no substantial difference in performance when using per-device calibration, however there were substantial differences across habitat, with a clear trend of the RMS UERE increasing with signal quality, from 0.35-0.39 beneath the house to 0.95-1.00 in an open setting, which is similar to the trend we noticed in generation-1 e-obs GPS collars (App. S4.3.1). It appears that, while the e-obs error estimates are proportional to the location-error variance with good satellite reception, they overestimate variance with poor satellite reception. We leave further examination to future studies.

**Table S4.11:**
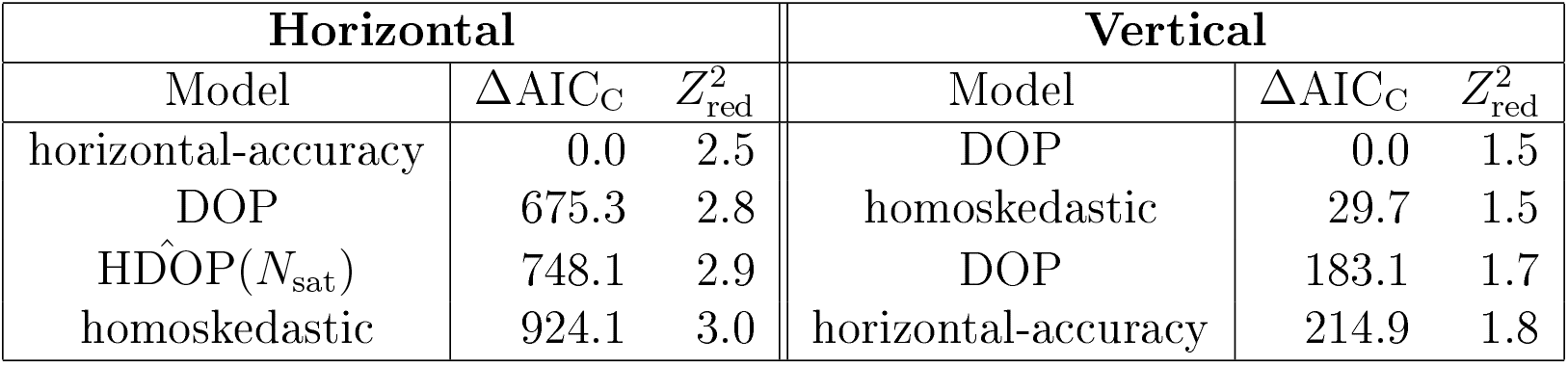
AIC_C_ differences and reduced *Z*^2^ values for error models fit to calibration data from 3 third-generation e-obs GPS tags.

### S4.4 GlobalTop

#### S4.4.1 GlobalTop PA6H

We analyzed calibration data from a GlobalTop PA6H GPS module, with location fixes taken every second, overnight. The PA6H GPS module returned all relevant DOP values and the number of satellites, which we compared to a model of homoskedastic location errors (Table S4.12). In terms of goodness of fit, homoskedastic errors were the best performing and VDOP values were worst performing, even for vertical errors.

**Table S4.12:** AIC_C_ differences and reduced *Z*^2^ values for error models fit to calibration data from a GlobalTop PA6H GPS module. Error models either consist of a single RMS UERE value, or a different RMS UERE for in-time and timed-out fixes. ^†^Due to autocorrelation, the AIC_C_ differences are exaggerated by a factor on the order of a hundred.

| Horizontal |  |  | Vertical |  |  |
| --- | --- | --- | --- | --- | --- |
| Model | $\Delta\text{AIC}_C^\dagger$ | $Z_{\text{red}}^2$ | Model | $\Delta\text{AIC}_C^\dagger$ | $Z_{\text{red}}^2$ |
| HDOP | 0.0 | 1.5 | homoskedastic | 0.0 | 0.6 |
| homoskedastic | 4,897.2 | 1.3 | HDOP | 155.7 | 0.6 |
| $\text{HDOP}(N_{\text{sat}})$ | 7,143.4 | 1.5 | $\text{HDOP}(N_{\text{sat}})$ | 1,273.5 | 0.7 |
| PDOP | 11,333.7 | 1.5 | PDOP | 3,484.8 | 0.6 |
| VDOP | 17,776.3 | 1.5 | VDOP | 6,395.1 | 0.7 |

### S4.5 Lotek

#### S4.5.1 Lotek Lifecycle 330

We recovered opportunistic calibration data for Lotek Lifecycle 330 GPS collars from twelve deceased Mongolian gazelles, all sampled in 23-hour intervals for 5 days to 8 months after death. This model Lotek collar records a fix type of either 3D or 3D-V, but insufficient 3D fixes were obtained for calibrating both location classes. The only other location-error information included was an ambiguous DOP value, which we tested against a model of homoskedastic location errors. The homoskedastic error model proved to be more predictive than the DOP values (Table S4.13), but there was substantial variation among GPS collars, with 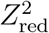 dropping from 2.3 to 1.2 when using individualized horizontal calibration.

**Table S4.13:** AIC_C_ differences and reduced *Z*^2^ values for error models fit to opportunistic calibration data from 12 Lotek Lifecycle 330 GPS collars.

| Model | $\Delta\text{AIC}_C$ | $Z_{\text{red}}^2$ |
| --- | --- | --- |
| homoskedastic | 0.0 | 2.3 |
| DOP | 50.9 | 2.5 |

#### S4.5.2 Lotek PinPoint 240

In addition to ambiguous ‘DOP’ values, Lotek PinPoint 240 GPS tags record a “duration” column, corresponding to the number of seconds lapsed before the GPS fix is completed or “time to fix” (TTF). When using these tags to track wood turtles, a large number of outliers were observed wherein recorded locations implied biologically implausible speeds, even when an using error-informed speed estimator on calibrated data (c.f., Sec. 2.3). Upon further inspection it was noticed that most of these outliers had a maximized fix duration of 70 seconds, which motivated our choice of candidate models.

We performed model selection using calibration data from two tags left under a tree for a day, with 10-minute sampling intervals. In the candidate models we compared homoskedastic location errors, DOP values, and our number-of-satellites precision model (*N*_sat_; Sec. S3.1). Furthermore, we also test for a larger RMS UERE value in the timed-out fixes. Our results are summarized in Table S4.14.

For the horizontal data, the combination of DOP value and distinct in-time/timed-out UERE parameters was the clear winner. The horizontal RMS UERE of the timed-out fixes was four times larger than that of the regular fixes, transforming most of the previously labeled outliers into normal data (Fig. S4.2). Results were slightly different for the vertical data, with our number-of-satellites precision model (*N*_sat_; S3.1) outperforming the DOP model. This suggests that the ambiguous DOP value is likely an HDOP value, and demonstrates the utility of our *N*_sat_ model—not only is it capable of outperforming a homoskedastic error model, but it can also outperform co-opted DOP values. RMS horizontal error was estimated to be 8.0–8.7 meters under ideal conditions (HDOP=1 & in-time). For the horizontal errors, there was slight heterogeneity among devices, with 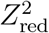 dropping from 2.5 to 2.4 when using individualized calibration.

**Figure S4.2:**
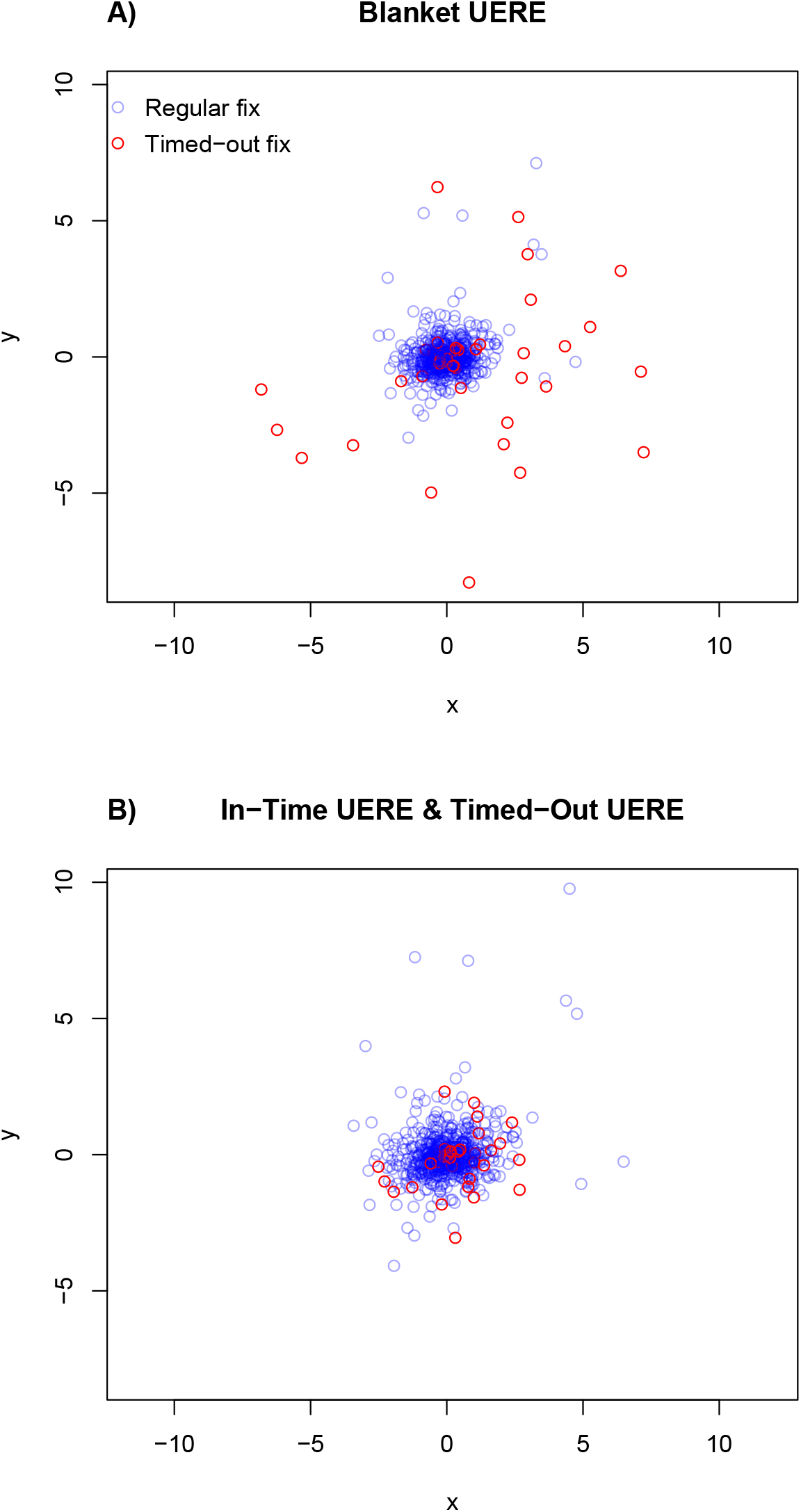
Residuals of calibration data for Lotek tags when **A**) considering all location fixes to share the same RMS UERE value and **B**) considering regular location fixes and timed-out location fixes as having distinct RMS UERE values. A shorter tailed residual distribution indicates a better performing error model.

**Table S4.14:** AIC_C_ differences and reduced *Z*^2^ values for error models fit to calibration data from 2 Lotek PinPoint 240 GPS tags. Error models either consist of a single RMS UERE value, or a different RMS UERE for in-time and timed-out fixes.

| Horizontal |  |  | Vertical |  |  |
| --- | --- | --- | --- | --- | --- |
| Model | $\Delta AIC_C$ | $Z_{red}^2$ | Model | $\Delta AIC_C$ | $Z_{red}^2$ |
| DOP & is(timeout) | 0.0 | 2.5 | $\hat{HDOP}(N_{sat})$ & is(timeout) | 0.0 | 1.7 |
| $\hat{HDOP}(N_{sat})$ & is(timeout) | 571.0 | 3.2 | DOP & is(timeout) | 57.4 | 2.0 |
| DOP | 602.2 | 3.3 | $\hat{HDOP}(N_{sat})$ | 140.0 | 1.9 |
| is(timeout) | 1,416.6 | 5.1 | is(timeout) | 279.7 | 2.2 |
| $\hat{HDOP}(N_{sat})$ | 1,434.0 | 4.6 | DOP | 452.0 | 2.4 |
| homoskedastic | 3,724.0 | 9.6 | homoskedastic | 773.1 | 2.3 |

#### S4.5.3 Lotek WildCell GPS-GSM

We collected over three days worth of hourly calibration data from five Lotek WildCell GPS-GSM collars. These data recorded an ambiguous DOP value and whether or not the location fix was “validated”. The “validated” column proved to be marginally informative, with validated location fixes having 8.5-9.4 meter RMS horizontal error and non-validated fixes having 1-3 times that amount, but the DOP values were not found to be informative (Table S4.15. There was moderate heterogeneity among devices, with the horizontal 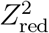 dropping from 1.8 to 1.5 upon individualized calibration.

**Table S4.15:** AIC_C_ differences and reduced *Z*^2^ values for error models fit to calibration data from 5 Lotek WildCell GPS-GSM tags. Error models either consist of a single RMS UERE value, or a different RMS UERE for validated and non-validated fixes.

| Horizontal |  |  | Vertical |  |  |
| --- | --- | --- | --- | --- | --- |
| Model | $\Delta AIC_C$ | $Z_{red}^2$ | Model | $\Delta AIC_C$ | $Z_{red}^2$ |
| is(validated) | 0.0 | 1.8 | DOP | 0.0 | 1.3 |
| homoskedastic | 8.0 | 1.8 | homoskedastic | 1.4 | 1.3 |
| DOP & is(validated) | 91.5 | 2.1 | DOP & is(validated) | 2.0 | 1.4 |
| DOP | 102.6 | 2.1 | is(validated) | 3.2 | 1.3 |

### S4.6 NTT Docomo

#### S4.6.1 NTT Docomo CTG-001G

We collected a weeks’ worth of hourly calibration data from three NTT Docomo CTG-001G units. The CTG-001G is a cellular-assisted GPS device that can triangulate with cellular towers, using Japan’s GSM equivalent–‘Freedom of Mobile Multimedia Access’ (FOMO) (Ishii et al., 2019). This device provided an error estimate in meters, which we tested against a model of homoskedastic location errors (Table S4.16). A location class was also provided, but the low-accuracy classes were too sparsely populated for analysis. We found the CTG-001G error estimates to be highly informative, and there was no substantial variation in device performance. The RMS location error was estimated to be 3.1–3.4 meters under ideal conditions.

**Table S4.16:** AIC_C_ differences and reduced *Z*^2^ values for error models fit to opportunistic calibration data from three NTT Docomo CTG-001G GPS device.

| Model | $\Delta\text{AIC}_C$ | $Z_{\text{red}}^2$ |
| --- | --- | --- |
| “error” | 0.0 | 1.9 |
| homoskedastic | 1,835.8 | 5.7 |

### S4.7 Ornitela

#### S4.7.1 Ornitela 20-gram tag (182902–182928)

We placed three 20-gram Ornitela GPS tags on a roof and three in a forest for 2 days, collecting data in 2-minute intervals. We tested joint error models using the reported HDOP values and number of satellites, but both of these models failed to outperform one of homoskedastic location errors (Table S4.17). Because the HDOP values and number of satellites were not informative, and because habitats differed among devices, individual calibrations were highly heterogeneous, with AIC_C_ decreasing 4,116.2 and 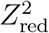 dropping from 2.2 to 1.6. 99% of the AIC_C_ difference was solely due to habitat, with forested location errors being twice that of rooftop errors. In principle, HDOP values can account for signal loss due to a wooded environment, but, again, the stated HDOP values were not informative.

**Table S4.17:** AIC_C_ differences and reduced *Z*^2^ values for error models fit to calibration data from six 20-gram Ornitela GPS tags.

| Horizontal |  |  | Vertical |  |  |
| --- | --- | --- | --- | --- | --- |
| Model | $\Delta\text{AIC}_C$ | $Z_{\text{red}}^2$ | Model | $\Delta\text{AIC}_C$ | $Z_{\text{red}}^2$ |
| homoskedastic | 0.0 | 2.2 | homoskedastic | 0.0 | 1.6 |
| HDOP | 763.7 | 2.4 | HDOP | 374.6 | 1.5 |
| $\text{HDOP}(N_{\text{sat}})$ | 1,717.8 | 2.5 | $\text{HDOP}(N_{\text{sat}})$ | 526.2 | 1.7 |

#### S4.7.2 Ornitela 25-gram tag (191771–191779)

We placed three 25-gram Ornitela GPS tags on a roof and three in a forest for 2 days, collecting data in 2-minute intervals. We tested joint error models using the reported HDOP values and number of satellites, but both of these models failed to outperform one of homoskedastic location errors (Table S4.18). The tags were very heterogeneous, as individualized calibration caused 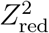 to drop from 3.4 to 2.1. Habitat alone could not explain these differences (ΔAIC_C_=2,357.8).

### S4.8 Sirtrack

#### S4.8.1 Sirtrack Pinnacle Solar G5C275F

We recovered opportunistic calibration data for Sirtrack Pinnacle Solar G5C275F GPS-Iridium collars from three deceased Mongolian gazelles, all sampled hourly for 3–12 months after death. After a first round of calibration, pitting HDOP versus our number-of-satellites model (*N*_sat_; S3.1) and homoskedastic location errors, we noticed a precipitous drop in the RMS error at precisely 32 seconds of “time on”, which we assume to be the ‘time to fix’ (TTF). Therefore, we included among our candidate models variants with separate location-estimate classes for TTF<32 seconds and TTF≥32 seconds (Table S4.19). TTF proved to be more informative than HDOP, which was not predictive in the Mongolian grasslands where the collars were used, and the median HDOP value was 1.2. The best-fit RMS UERE was 1.66 meters (1.63–1.68 meters) for location estimates with 32 seconds or more of “time on”, and 6.56 meters (6.48–6.64 meters) for location estimates with less than 32 seconds of “time on”. The sub-dekameter scale of the former location errors suggest on-board processing of the location estimates with sufficient “time on”, even though the scheduled sampling interval was only one hour. Most likely, this device aggregates multiple location fixes into one reported estimate. Negligible heterogeneity was observed among devices.

**Table S4.18:** AIC_C_ differences and reduced *Z*^2^ values for error models fit to calibration data from six 25-gram Ornitela GPS tags.

| Horizontal |  |  | Vertical |  |  |
| --- | --- | --- | --- | --- | --- |
| Model | $\Delta\text{AIC}_C$ | $Z^2_{\text{red}}$ | Model | $\Delta\text{AIC}_C$ | $Z^2_{\text{red}}$ |
| homoskedastic | 0.0 | 3.4 | homoskedastic | 0.0 | 1.7 |
| HDOP | 334.0 | 3.6 | HDOP | 578.6 | 1.8 |
| $\hat{\text{HDOP}}(N_{\text{sat}})$ | 1,028.8 | 3.5 | $\hat{\text{HDOP}}(N_{\text{sat}})$ | 2,310.3 | 2.1 |

**Table S4.19:** AIC_C_ differences and reduced *Z*^2^ values for error models fit to opportunistic calibration data from 3 Sirtrack GPS collars. Error models either consist of a single RMS UERE value or separate RMS UERE values for location estimates resolved before or after 32 seconds.

| Model | $\Delta\text{AIC}_C$ | $Z^2_{\text{red}}$ |
| --- | --- | --- |
| is($\text{TTF} < 32$ sec) | 0.0 | 4.2 |
| $\hat{\text{HDOP}}(N_{\text{sat}})$ & is($\text{TTF} < 32$ sec) | 206.8 | 4.1 |
| HDOP & is($\text{TTF} < 32$ sec) | 1,707.6 | 4.3 |
| $\hat{\text{HDOP}}(N_{\text{sat}})$ | 11,343.1 | 4.8 |
| homoskedastic | 13,681.6 | 5.5 |
| HDOP | 13,810.6 | 5.0 |

### S4.9 Technosmart

#### S4.9.1 Technosmart GiPSy 5

We collected calibration data from two Technosmart GiPSy 5 tags, both sampled in 10 second intervals for 30 minutes and 9 hours. Although this device came with informative HDOP values, they were outperformed by our number-of-satellites model (*N*_sat_; S3.1), as summarized in table S4.20. RMS horizontal error was estimated to be 10.1–10.5 meters under ideal conditions (*N*_sat_=12). Differences in tag calibration were not substantial.

**Table S4.20:**
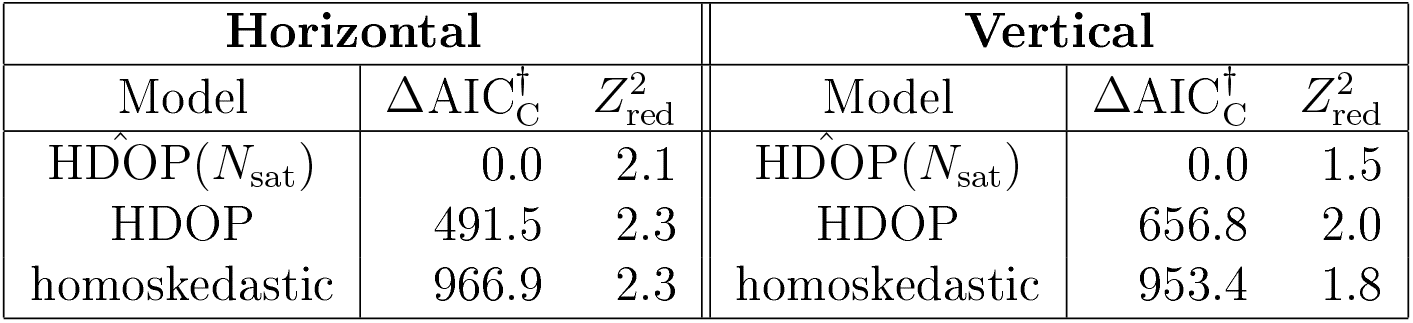
AIC_C_ differences and reduced *Z*^2^ values for error models fit to calibration data from 2 Technosmart GiPSy GPS tags. ^†^Due to autocorrelation, the AIC_C_ differences are exaggerated by a factor on the order of ten.

| Horizontal |  |  | Vertical |  |  |
| --- | --- | --- | --- | --- | --- |
| Model | $\Delta\text{AIC}_C^\dagger$ | $Z_{\text{red}}^2$ | Model | $\Delta\text{AIC}_C^\dagger$ | $Z_{\text{red}}^2$ |
| $\text{HDOP}(N_{\text{sat}})$ | 0.0 | 2.1 | $\text{HDOP}(N_{\text{sat}})$ | 0.0 | 1.5 |
| HDOP | 491.5 | 2.3 | HDOP | 656.8 | 2.0 |
| homoskedastic | 966.9 | 2.3 | homoskedastic | 953.4 | 1.8 |

### S4.10 Telemetry Solutions

#### S4.10.1 TS Quantum 4000 Enhanced

Model Quantum 4000 Enhanced Telemetry Solutions (TS) GPS tags report an HDOP, number of satellites, and location fix type, which is either 2D or 3D depending on whether or not three or more more satellites were in reception. Both the 3D and 2D fixes are accompanied with HDOP values, and the 2D HDOP values were on-average larger than the 3D HDOP values, which is expected to be the case if the 3D and 2D HDOP values are on the same scale.

We performed model selection on calibration data from two tags left under a tree for a month, with 2-hour sampling intervals. We considered models with 1-2 RMS UERE values for the 3D/2D fix types, and we also pitted the HDOP values against homoskedastic location errors and our number-of-satellites precision model (*N*_sat_; S3.1). Our results are summarized in Table S4.21. The highest performing model featured HDOP-informed errors, but different RMS UERE values for the 3D and 2D fixes. The 2D RMS UERE was approximately four times larger than the 3D RMS UERE. RMS horizontal error was estimated to be 6.5–7.4 meters under ideal conditions (HDOP=1 & 3D). There was no significant heterogeneity between the two devices.

**Table S4.21:** AIC_C_ differences and reduced *Z*^2^ values for error models fit to calibration data from 2 Telemetry Solutions GPS tags. Error models either consist of a single RMS UERE value or separate RMS UERE values for 2D and 3D location estimates.

| Model | $\Delta\text{AIC}_C$ | $Z_{\text{red}}^2$ |
| --- | --- | --- |
| HDOP & is(2D) | 0.0 | 2.4 |
| $\text{HDOP}(N_{\text{sat}})$ & is(2D) | 225.4 | 3.3 |
| $\text{HDOP}(N_{\text{sat}})$ | 248.6 | 3.6 |
| HDOP | 286.6 | 3.9 |
| is(2D) | 418.5 | 3.7 |
| homoskedastic | 637.3 | 4.7 |

### S4.11 Telonics

“Gcn4” Telonics data come standard with both an HDOP value and a “horizontal error” estimate in meters, with the two values not proportional and instead corresponding to error estimates of differing quality. Telonics documentation describes the new horizontal error estimate as a 100% confidence radius, but this is unlikely to be the case. In personal communications with Telonics, the specific quantile or number of standard deviations was unknown. Given that the Gen4 horizontal error estimates are measured in meters, we considered a model where they are proportional to the RMS horizontal error. Furthermore, Gen4 Telonics data contain two distinct location classes—regular GPS location fixes and “quick-fix points” (QFPs). The quick fixes do not come with a corresponding error estimate and Telonics documentation explicitly states that their RMS UERE value is larger than that of the ordinary fixes. Based on the following results, `ctmm`’s `as.telemetry` function ignores the Telonics Gen4 error estimate and instead uses HDOP values. Location classes are also designated, based on whether or not the location fix is a QFP.

#### S4.11.1 Telonics Gen4 GPS-VHF

To determine the calibration parameters of Telonics Gen4 GPS-VHF collars, we used opportunistic calibration data collected from one deceased lowland tapir (*Tapirus terrestris*) and two collars that detached from lowland tapir, all sampled hourly in the Pantanal for 2 to 25 days. We pitted the Telonics Gen4 “horizontal error” estimate against HDOP values, our number-of-satellites model (*N*_sat_; S3.1), and homoskedastic errors (Table S4.22). The HDOP values proved to be the most predictive, with an RMS UERE of 6.2 meters (6.0–6.4 meters). However, the HDOP model was not substantially different from a model of homoskedastic location errors, and, surprisingly, the “horizontal error” estimates did not appear proportional to RMS location errors. RMS horizontal error was estimated to be 4.8–5.1 meters under ideal conditions (HDOP=0.8). Variation among devices was not substantial.

Only one of the three devices recorded QFPs. By fitting an error model with two RMS UERE values—one for regular location estimates and another for QFPs—we found the QPF RMS UERE to be about twice that of the regular RMS UERE, which is consistent with Telonics documentation.

**Table S4.22:** AIC_C_ differences and reduced *Z*^2^ values for error models fit to calibration data from 3 Telonics Gen4 VHF GPS collars.

| Horizontal |  |  | Vertical |  |  |
| --- | --- | --- | --- | --- | --- |
| Model | $\Delta\text{AIC}_C$ | $Z^2_{\text{red}}$ | Model | $\Delta\text{AIC}_C$ | $Z^2_{\text{red}}$ |
| HDOP | 0.0 | 1.9 | homoskedastic | 0.0 | 1.3 |
| $\widehat{\text{HDOP}}(N_{\text{sat}})$ | 77.9 | 2.0 | HDOP | 52.9 | 1.3 |
| homoskedastic | 79.5 | 1.8 | $\widehat{\text{HDOP}}(N_{\text{sat}})$ | 58.0 | 1.3 |
| horizontal-error | 437.8 | 2.8 | horizontal-error | 418.0 | 1.8 |

#### S4.11.2 Telonics Gen4 GPS-Iridium

We performed a follow-up analysis with newer Telonics Gen4 GPS-Iridium collars, using opportunistic calibration data from three detached collars, all sampled hourly in the Cerrado for 2-3 days. Our model comparison resulted in the same model ranking for horizontal errors (Table S4.23), with the selected model based on HDOP, though the GPS-Iridium RMS UERE value of 6.9 meters (6.4–7.4 meters) was slightly larger than that of the GPS-VHF model (6.0–6.4 meters). RMS horizontal error was estimated to be 2.6–3.0 meters under ideal conditions (HDOP=0.4). Variation among devices was not substantial.

**Table S4.23:** AIC_C_ differences and reduced *Z*^2^ values for error models fit to calibration data from 3 Telonics Gcn4 Iridium GPS collars.

| Horizontal |  |  | Vertical |  |  |
| --- | --- | --- | --- | --- | --- |
| Model | $\Delta\text{AIC}_C$ | $Z_{\text{red}}^2$ | Model | $\Delta\text{AIC}_C$ | $Z_{\text{red}}^2$ |
| HDOP | 0.0 | 1.7 | HDOP | 0.0 | 1.5 |
| $\text{HDOP}(N_{\text{sat}})$ | 23.8 | 1.9 | $\text{HDOP}(N_{\text{sat}})$ | 6.1 | 1.2 |
| homoskedastic | 49.7 | 2.0 | homoskedastic | 18.0 | 1.2 |
| horizontal-error | 171.7 | 2.6 | horizontal-error | 71.1 | 1.4 |

### S4.12 UniKN

#### S4.12.1 UniKN Logger GPS tag

We placed three UniKN Logger solar tags on a roof for two weeks, where under sunlight conditions they were able to obtain location fixes approximately every 16 minutes. This model UniKN Logger tag did not report an HDOP value, but it did report the number of satellites, with which we applied number-of-satellites precision model (S3.1). After removing one gross outlier, the best performing model was our number-of-satellites model (Table S4.24). RMS horizontal error was estimated to be 9.3–10.1 meters under ideal conditions (*N*_sat_=12). There was no significant variability among devices.

**Table S4.24:** AIC_C_ differences and reduced *Z*^2^ values for error models fit to calibration data from 3 UniKN Logger GPS tags.

| Horizontal |  |  | Vertical |  |  |
| --- | --- | --- | --- | --- | --- |
| Model | $\Delta\text{AIC}_C$ | $Z_{\text{red}}^2$ | Model | $\Delta\text{AIC}_C$ | $Z_{\text{red}}^2$ |
| $\text{HDOP}(N_{\text{sat}})$ | 0 | 2.1 | $\text{HDOP}(N_{\text{sat}})$ | 0 | 1.1 |
| homoskedastic | 66.9 | 2.3 | homoskedastic | 57.0 | 1.2 |

### S4.13 Vectronic

#### S4.13.1 Vectronic GPS Plus

We collected testing data on 6 Vectronic GPS Plus collars, with fixes taken in 15-minute intervals for 2 hours. With less statistically efficient methods this might have been too little data to analyze, but with our minimum-variance unbiased (MVU) parameter estimators and AIC_C_ values, and the ability to pool the 6 datasets into a joint estimate, this was enough data to provide 50 degrees of freedom in the RMS UERE estimates. In addition to ambiguous DOP values, the devices also recorded an error estimate in meters, which was described in Vectronic’s documentation as “the difference [m] between the real position and the transmitted position”. We compared (ambiguous) DOP values and recorded error estimates (in meters) to a model of homoskedastic location errors, and found the on-board error estimates to be the most predictive (Table S4.25). Differences in performance among devices were not very significant. The estimated RMS UERE for the recorded error estimates was 0.80-1.06, which is consistent with these being estimates of the RMS location error.

**Table S4.25:** AIC_C_ differences and reduced *Z*^2^ values for error models fit to calibration data from 6 Vectronic GPS Plus collars.

| Horizontal |  |  | Vertical |  |  |
| --- | --- | --- | --- | --- | --- |
| Model | $\Delta\text{AIC}_C$ | $Z_{\text{red}}^2$ | Model | $\Delta\text{AIC}_C$ | $Z_{\text{red}}^2$ |
| “error” | 0 | 1.5 | “error” | 0 | 0.9 |
| DOP | 31.4 | 1.7 | DOP | 39.6 | 1.2 |
| homoskedastic | 49.3 | 2.8 | homoskedastic | 46.4 | 2.0 |

#### S4.13.2 Vectronic Vertex Lite-3D GPS-Iridium

We collected calibration data on 18 Vectronic Vertex Lite-3D GPS-Iridium collars, with fixes taken in 15-minute and 4-hour intervals for 11 days. We compared the reported (ambiguous) DOP values to a model of homoskedastic location error, and found them to be only mildly informative (Table S4.26). RMS horizontal error was estimated to be 16.3–16.9 meters under ideal conditions (HDOP=0.8). There was slight variability among devices, with 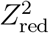 dropping from 2.5 to 2.4 upon individualized calibration.

**Table S4.26:** AIC_C_ differences and reduced *Z*^2^ values for error models fit to calibration data from 18 Vectronic Vertex Lite GPS collars.

| Horizontal |  |  | Vertical |  |  |
| --- | --- | --- | --- | --- | --- |
| Model | $\Delta\text{AIC}_C$ | $Z_{\text{red}}^2$ | Model | $\Delta\text{AIC}_C$ | $Z_{\text{red}}^2$ |
| DOP | 0 | 2.5 | DOP | 0 | 1.5 |
| homoskedastic | 513.0 | 2.6 | homoskedastic | 281.6 | 1.6 |

### S4.14 Vemco

Vemco positioning system (VPS) devices are analogous to GPS, but with acoustic communication to an array of fixed receivers instead of electromagnetic communication to a constellation of satellites, so that they can function underwater where GPS signals rapidly attenuate. VPS location estimates come with an ?PE’ value, which Vemco documentation describes as being analogous to GPS HDOP and requiring site specific calibration.

#### S4.14.1 Vemco HR2-V9 VPS

We attached 6 V9 180-kHz VPS tags to static moorings (metal poles fixed in cement anchors) and collected data over the span of 3 months in a Vemco HR2 high-residence-receiver network. The median sampling interval was 28 seconds, but some location estimates were obtained within a fraction of a second. Inspection of the residual correlogram (when using the ?PE null model) revealed that approximately half of the autocorrelation decayed within minutes, which is similar in behavior to GPS. The remaining autocorrelation took on the order of a day to decay, which could be due to movement in the calibration tags, filtering in the VPS location estimates, or some other property of VPS fixes.

We considered three candidate error models: a model of homoskedastic location errors, a null model of HDOP = HPE, and a model of HDOP = R.MSE, where R.MSE was an unknown column in the data that we hoped might correspond to an RMS location-error estimate. We found HPE to provide the best error model (Table S4.27), with an RMS UERE of 5.8 meters. There was substantial variability among locations, with RMS UEREs ranging 3–9 meters and the reduced *Z*^2^ statistic shrinking from 2.1 to 1.4 upon individualized calibration, but there was no significant dependence on depth.

**Table S4.27:**
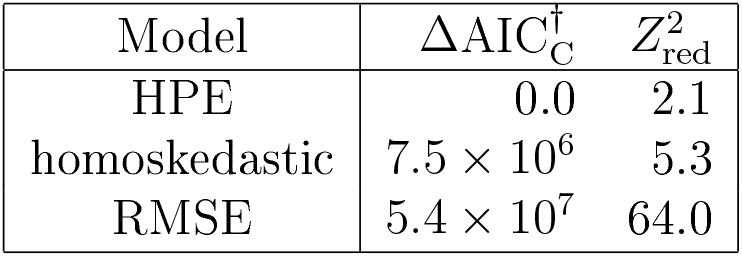
AIC_C_ differences and reduced *Z*^2^ values for error models fit to calibration data from 6 VPS tags. ^†^Due to autocorrelation, the AIC_C_ differences are exaggerated by a factor on the order of ten.

## S5 Error-informed statistics for outlier estimation

### S5.1 Distance statistics for outlier estimation

First we consider a location sample **r** = (*x, y*) and corresponding displacement vector **d** = **r** – **μ**_0_, as measured from a reference point **μ**_0_. For the reference point **μ**_0_ we consider the geometric median, which can be robustly estimated, and we ignore its relatively small 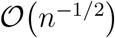 error, but other choices are also possible. In considering a normal distribution for the sampled location

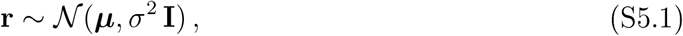

with unknown true location ***μ*** and known location error variance *σ*^2^, which we do not assume to be stationary in time, the distribution of the displacement vector **d** is then distributed according to

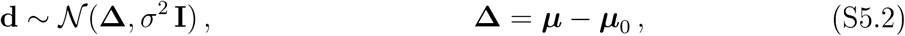

where **Δ** is the true displacement vector. Choosing a coordinate system with the *x*-axis parallel to **Δ**, our distribution can be represented in polar coordinates with **d** = (*d* cos *θ*, *d* sin *θ*) as

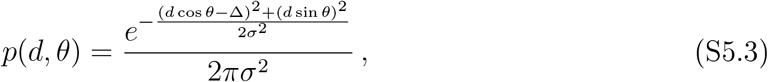

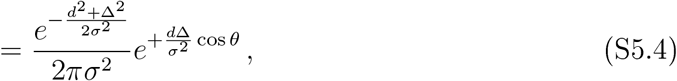

where the circular error assumption is critical to obtaining this tractable relation. The marginal distribution of *d* is then

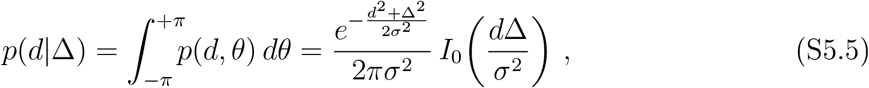

where *I_n_* is the modified Bessel function of the first kind. The log-likelihood function is then given by

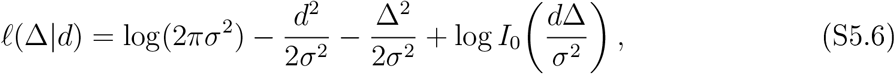

and so by differentiation the maximum likelihood estimate 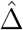 of the true distance Δ satisfies the transcendental equation

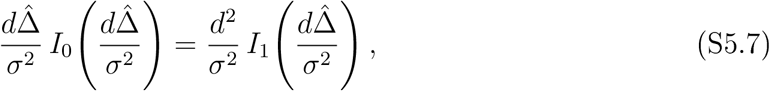

or equivalently

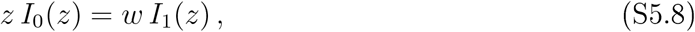

where 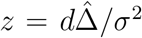 and *w* = *d*^2^/*σ*^2^. We solve this equation by first Taylor expanding from *z* = 0 or *z* = *w*, whichever is closer, and then Newton-Raphson iterating, by expanding the Bessel functions around the current best estimate. It can be shown that 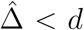, as should result from the inclusion of a location-error model, and 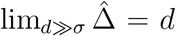, which also must be the case. Interestingly, for all *d*^2^ < 2*σ*^2^, 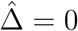.

Finally, from the Hessian of the log-likelihood function, and after some algebraic simplification using (S5.7), we have the uncertainty estimate

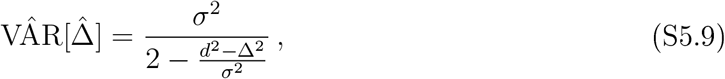

which becomes larger than *σ*^2^/2 as the improved distance estimate 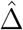 becomes smaller than the naive distance estimate, *d*.

### S5.2 Speed statistics for outlier estimation

For speed-based outlier detection, we want to avoid accidentally ruling out realistically fast displacements during which motion would be ballistic. Therefore, it is sufficient to consider speed as distance Δ = |***μ***_2_ – ***μ***_1_| divided by time, ignoring tortuosity. To condition on as few locations as possible, we only consider sequential pairs. For the distance estimate, we use the results of the previous section, where the variance is now the sum of the variances of the two locations, or 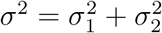. Otherwise, all relations there are the same.

In dividing distance by the sampled time 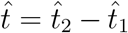, there are situations where roundoff or truncation errors in t result in large values of the distance/time estimate, with otherwise usable data (permitting a location-error model). To accommodate truncated times, it is easier to model the imparted location error, when considering the rounded time as true, than it is to directly consider the temporal error caused by rounding. In the `outlie` method, we use a greatest common divisor routine to attempt to automatically detect the temporal sampling resolution *δ*, which is usually 1 second, but frequently a minute, hour, or day. Then we count the maximum *M* = max *M_i_* of the number of location fixes *M_i_* per truncated time 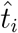. The minimum time to fix is then estimated by *δ/M*, which we subsequently use in place of zero for our speed estimates. Note that none of these crude estimates are used in movement analysis, but only for outlier detection.

Thus far, we have estimated speeds over the intervals, such as (*t*_1_, *t*_2_) and not specifically at the sampled times. To clean the data, we assign the highest speeds to the most suspect times. We proceed in ordered fashion from the smallest speed estimate to the largest speed estimate and assign each speed estimate to the adjacent time with the larger adjacent speeds, with the newer (larger) speed estimates overwriting the older (smaller) speed estimates. At the endpoints we compare to both the first (adjacent) times and the second times out. For convenience, we precede this loop by a reverse-ordered loop with reverse (low-speed) assignment to ensure the low-speed times are populated as well. If any outliers are removed, the speeds can be re-estimated to test for consecutive outliers. This speed assignment heuristic has substantial advantages over simpler heuristics, such as assigning the average adjacent speed estimate, which can result in false positives, or assigning the minimum adjacent speed speed estimate, which can result in false negatives.

### S5.3 Height statistics for outlier estimation

Here we consider the same problem but with vertical location estimates, *z*, which may represent height relative to the ellipsoid, sea level, terrain, or some other quantity. Given calibrated data with known variance *σ*^2^ and unknown true height *μ*

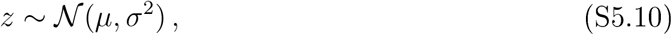

then after marginalizing over the two directions—up and down—the distribution of |*z*| can be expressed

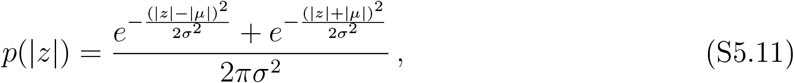

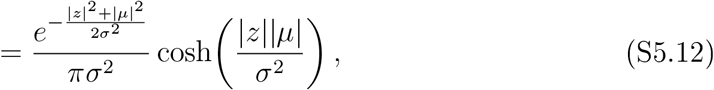

so that the log-likelihood of |*μ*| is given by

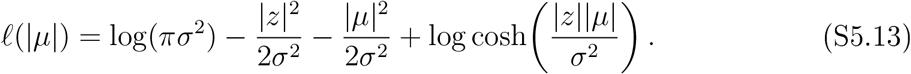

Next, by differentiation, the maximum likelihood estimate of |*μ*| must satisfy the transcendental equation

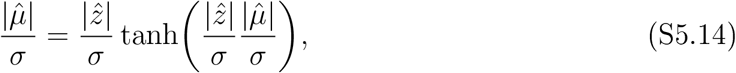

or equivalently

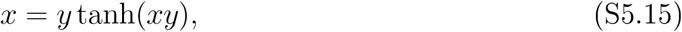

where 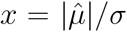 and 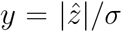. We solve this transcendental equation by expanding from *x* = 0 or *x* = *y*, whichever is closer, and then Newton-Raphson iterating, by expanding the hyperbolic tangent around the current best estimate. For 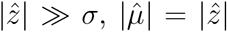, as expected, and for 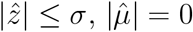.

Finally from the Hessian of the log-likelihood function, we have the uncertainty estimate

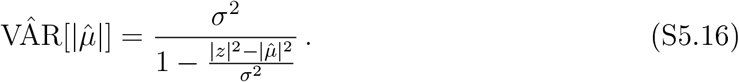

## S6 Errors in variograms

### S6.1 Fast variogram of errors

For evenly sampled data *z*(*t*), the empirical variogram is given by

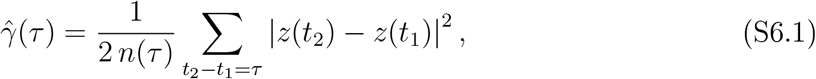

here in one dimension and where 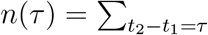 denotes the number of location pairs with time-lag *τ* between them. If location errors are additive, mean zero and uncorrelated, then the “observed” empirical variogram decomposes into the semi-variance function of the true movement plus the semi-variance of the location error.

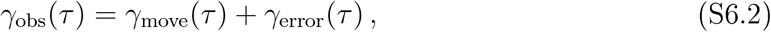

The data provides us with *γ*_obs_(*τ*), and the fitted movement model provides us with *γ*_move_(*τ*). Therefore, it is useful to have an estimate of *γ*_error_(*τ*) to finalize our comparison between the model and data.

The error variogram is given by

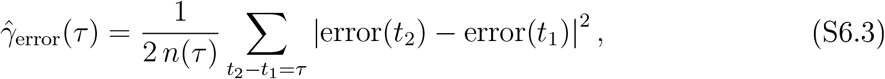

where the location errors are unknown, but can safely be considered as independent if the data are not sampled at exceptionally high frequencies (i.e., ≥ 1/min). Therefore, for positive time lags we have simply

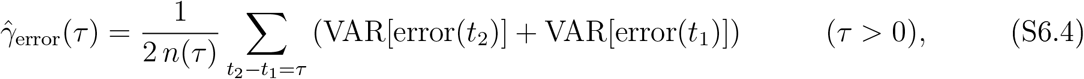

which we can estimate from DOP and RMS UERE values.

Naively, relation (S6.4) involves a sum for each lag, which would imply an 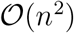 compu-tational cost for *n* sampled locations. In (Marcotte, 1996), an 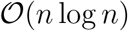 algorithm is given for the ordinary variogram (S6.1), by leveraging the fast Fourier transform (FFT). Here, we provide an 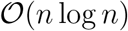 algorithm FFT algorithm for the error variogram, under the assumption that we have the timeseries VAR[error(*t*)]. The trick begins by including an indicator function *χ*(*t*) that evaluates to 1 if time t is sampled and 0 otherwise.

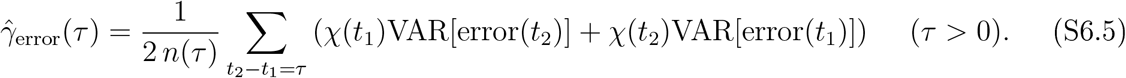

Then we may express the constrained sum with a Kronecker delta function as

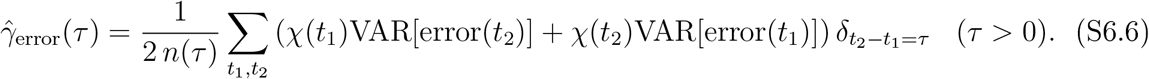

Next, by expanding the Kronecker delta functions in a suitable harmonic basis (Péron et al., 2016), we have for positive lags (S6.7) (S6.8)

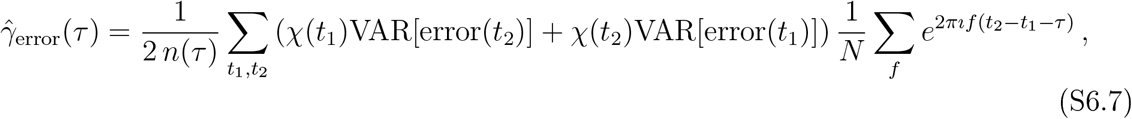

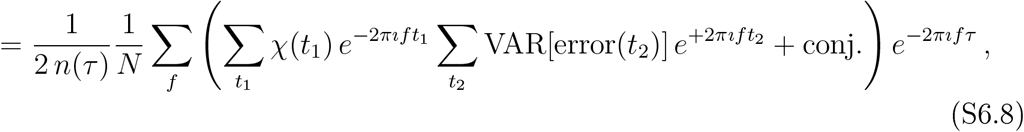

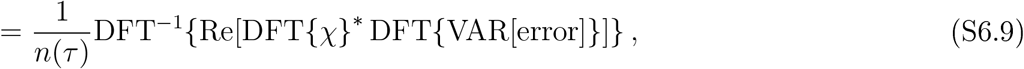

in terms of discrete Fourier transform operations.

### S6.2 Error-adjusted variogram

Here, instead of estimating the influence of error on the conventional variogram, we provide a semi-variance estimator that accounts for error. Because our error adjustment cannot be implemented with fast Fourier transforms, we consider the weighted variogram

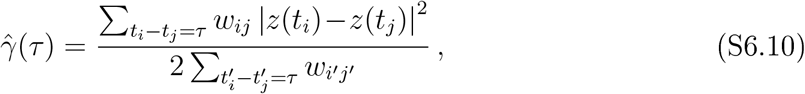

which is more robust to irregular sampling (Fleming et al., 2014a; Fleming and Calabrese, 2015). It is straightforward to show that this estimator maximizes the *χ*^2^ log-likelihood function

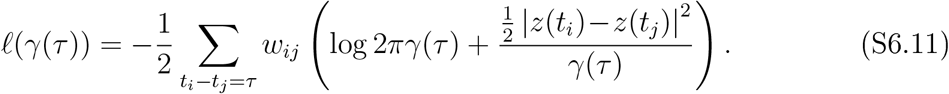

Therefore, we can generalize this model to erroneous observations 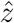 with observation error.

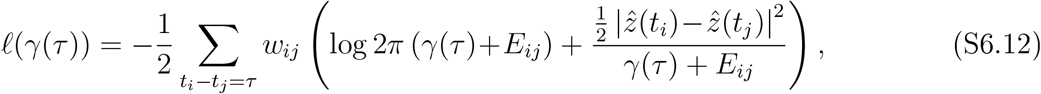

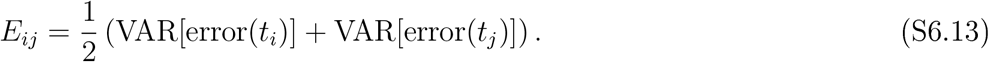

The maximum-likelihood estimate then satisfies the relation

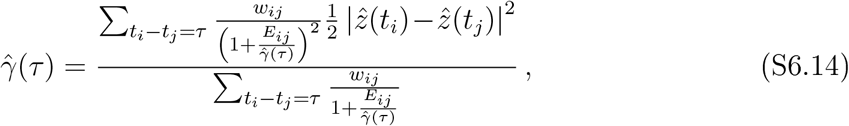

which can be used to solve for 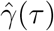 iteratively. An example calculation is given in Fig S6.1.

**Figure S6.1:**
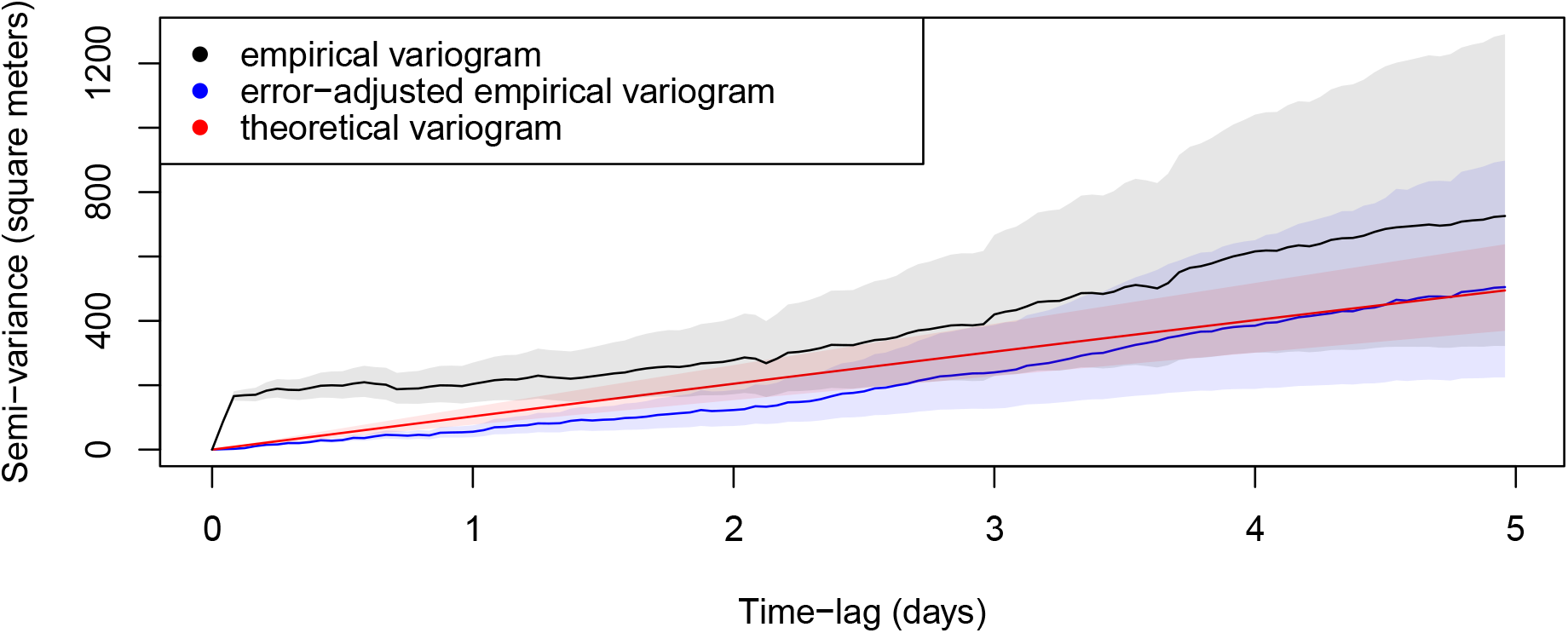
Conventional (black) and error-adjusted (blue) empirical variograms and AIC-best theoretical semivariance function (red) for a GPS-tracked wood turtle. The variogram is an unbiased measure of autocorrelation structure that estimates the average square displacement (*y*-axis) between two locations that are some time lag apart (*x*-axis), modulo a factor of ¼ in two dimensions. In the conventional variogram (black) this square displacement is a combination of movement and location error, which causes an initial discontinuity or ‘nugget effect’. In this case, two locations sampled closely together in time will be almost 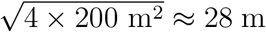 apart (RMS average), because of substantial location error from underbrush and canopy. The error-adjusted variogram, directly based on the data, and theoretical semivariance function, based on the AIG-best movement model, both agree that it actually takes the wood turtle ~3 days, on average, to be found this far from the previous location.

## Footnotes

1 By ‘Argos Doppler-shift’, we specifically refer to location estimates derived from communication with the Argos satellite system and not GPS location data that arc transmitted via the Argos satellite system.

2 This is only true if the Argos error ellipses arc to be trusted, which is at least approximately true.

3 Wo arc not fixing the relative variances of the location classes and estimating an overall variance parameter as in Jonsen et al. (2005), nor making any comparison with that approach here.

